# TECPR2 is a Rab5 effector that regulates endosomal cargo recycling

**DOI:** 10.1101/2024.10.03.616509

**Authors:** Sankalita Paul, Rajat Pant, Poonam Sharma, Kshitiz Walia, Suhasi Gupta, Adhil Aseem, Kamlesh Kumari Bajwa, Ruben George, Yudish Varma, Tripta Bhatia, Rajesh Ramachandran, Amit Tuli, Mahak Sharma

## Abstract

Small GTP-binding (G) proteins of the Rabs, Arfs, and Arls (Arf-like) family mediate the recruitment of their effectors to subcellular membrane-bound compartments, which in turn mediates vesicle budding, motility, and tethering. Here, we report that Tectonin-β-propeller repeat containing protein 2 (TECPR2), a protein mutated in a form of hereditary sensory and autonomic neuropathy (HSAN), is an effector of early endosomal Rab protein, Rab5, and interacts with Rab5 via its C-terminal TECPR repeats. The HSAN-associated TECPR2 (R1336W) missense variant was deficient in Rab5-binding and, consequently, in membrane recruitment. TECPR2-depletion led to perinuclear collapse of recycling endosomes and increased overlap of sorting and degradative subdomain markers on early endosomes. Consistent with a possible role in endocytic recycling, we observed impaired recycling and increased lysosomal degradation of α5β1 integrin receptors upon TECPR2 knockdown. TECPR2 regulates the early endosomal localization of the cargo adaptor for β1 integrins, SNX17, and its-associated protein complex WASH, which mediates actin polymerization on early endosomes. Finally, TECPR2 depletion in the zebrafish model resulted in delayed motility and changes in the neuromuscular junction. Our study supports an early endosomal role for TECPR2 in cargo recycling and provides insights into how its loss-of-function results in a neurodegenerative genetic disorder.

## Introduction

Vesicular transport consists of several steps, including vesicle budding at the donor membranes, vesicle motility on the microtubule tracks, tethering with the acceptor membranes, and finally membrane fusion. Rabs, Arfs, and Arls (Arf-like) are small GTP-binding (G) proteins of the Ras superfamily that act as master regulators of vesicular transport to and from these membranes. When bound to GTP, small G proteins localize to specific subcellular compartments and recruit effector proteins to these membranes. These effector proteins control different steps of vesicular transport (Grosshans et al., 2006).

Rab5 and Rab7 are two key Rab proteins that orchestrate the various steps of early to late endosomal transport. Rab5 promotes homotypic fusion and motility of early endosomes and recruits the Rab7 GEF, Mon1-CCZ1, to early endosomes, which in turn mediates Rab7 activation and maturation of early endosomes to late endosomes (Poteryaev et al., 2010; Rink et al., 2005). Early to late endosome maturation is coordinated with the retrieval of cargo receptors destined for recycling to the cell surface or towards the trans-Golgi network (TGN) from maturing early endosomes (Naslavsky and Caplan, 2018; Naslavsky and Caplan, 2023). The late endosomal G protein Rab7 recruits the microtubule motor dynein-dynactin complex via its effector RILP to mediate retrograde motility of late endosomes towards the perinuclear region (Zhang et al., 2009). Tethering and fusion of late endosomes with lysosomes is mediated by Rab7 and Arl8b, an Arl family member that localize to lysosomes. Arl8b interacts with the heterohexameric tethering factor, <u>HO</u>motypic fusion and <u>P</u>rotein <u>S</u>orting (HOPS) complex, and Rab7 effector PLEKHM1, a multidomain adaptor protein that mediates lysosome fusion with late endosomes and amphisomes (Khatter et al., 2015; Marwaha et al., 2017; Schleinitz et al., 2023). Rab2, another Rab protein, interacts with the HOPS complex and regulates tethering and fusion of amphisomes and late endosomes with lysosomes (Lorincz et al., 2017; Schleinitz et al., 2023).

Our ongoing interest in how membrane fusion machinery on lysosomes is regulated led us to a prior study describing the interaction of the HOPS complex with a protein known as Tectonin- β-propeller repeat containing protein 2 (TECPR2) (Stadel et al., 2015). TECPR2 was initially identified as an interacting partner of human autophagy-related 8 family members (LC3 and GABARAPs) (Behrends et al., 2010). In a subsequent study, a frameshift deletion in TECPR2 (L1139Rfs*75) was identified in the autosomal-recessive form of an ultra-rare genetic disorder, hereditary sensory and autonomic neuropathy (HSAN) 9 (previously known as spastic paraplegia- 49; OMIM 615031) (Oz-Levi et al., 2012). HSANs are neurodevelopmental and neurodegenerative disorders that are genetically and clinically heterogeneous and are characterized by the progressive loss of autonomic and sensory peripheral nervous system functions.

The N-terminal region (1-357 a.a.) of TECPR2 contains a WD40 domain predicted to fold into a seven-bladed β-propeller fold. The middle region (358-801 a.a.) lacks discernible domains, whereas the C-terminal region (802-1407 a.a.) contains six TECPR-repeats, which are predicted to form a double β-propeller motif. The last four residues of TECPR2 (WEVI; 1407-1411 a.a.) constitute a conventional W-type LC3-interacting region (LIR) motif that binds LC3 and GABARAP proteins. (Fig. 1A). Frameshift founder mutations in TECPR2 (L1139Rfs*75 and L440Rfs*19) lead to the formation of a truncated protein, which is not stable and is degraded in cells (Nalbach et al., 2023; Oz-Levi et al., 2012). To date, several HSAN disease-associated mutations in TECPR2 have been identified, as illustrated in Fig. 1A (Neuser et al., 2021). Notably, the pathogenic mutations in TECPR2 map to conserved residues in the N-terminal WD40 and C-terminal TECPR repeats, indicating that these regions play critical roles in regulating TECPR2 function (Fig. 1A).

**Figure 1:**
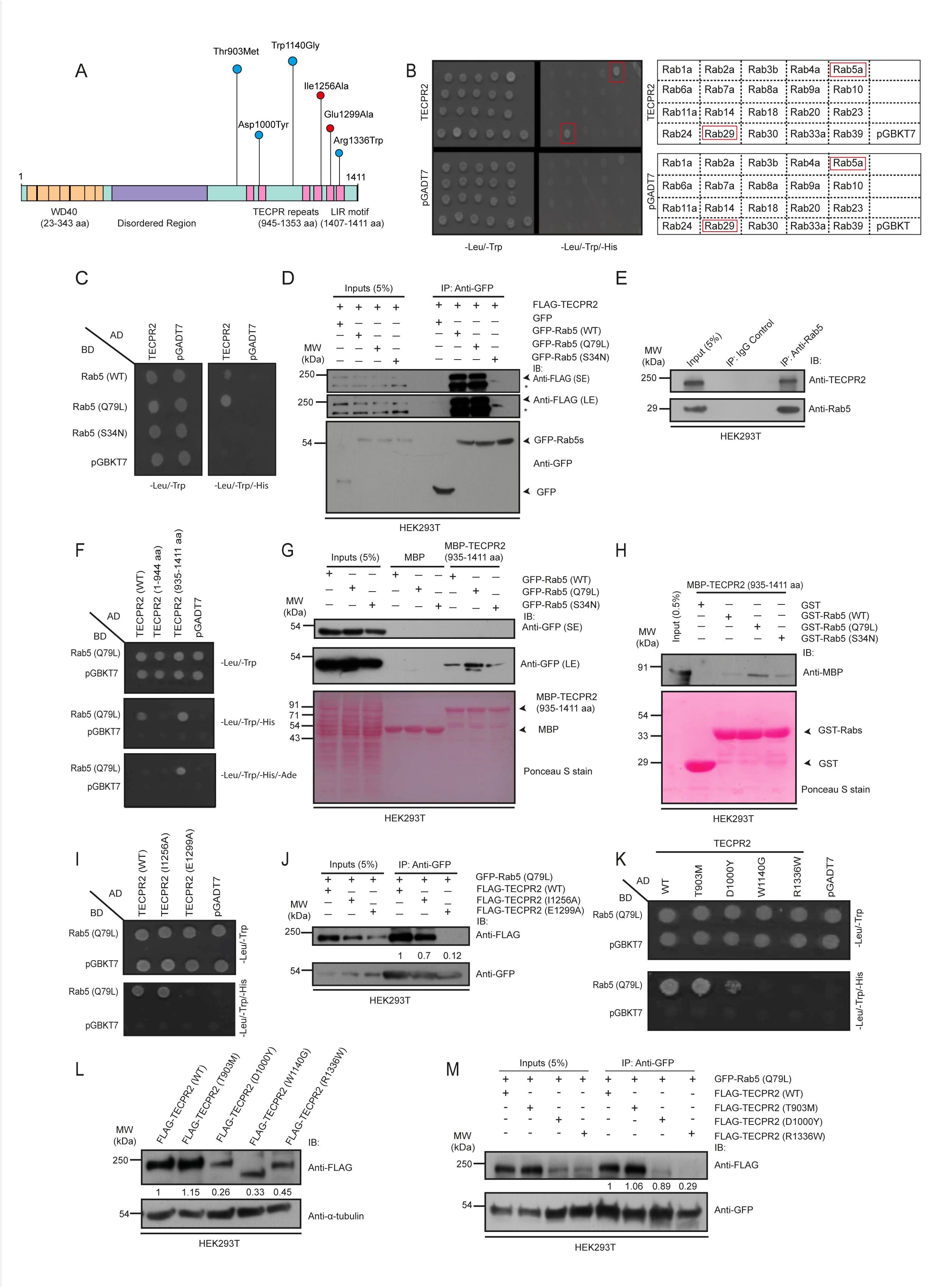
TECPR2 directly binds to small G protein Rab5 via its C-terminal TECPR repeats. A) A schematic of domain architecture of TECPR2 showing the WD40 repeat domain at the N- terminus, the middle-disordered region, and the TECPR repeats at the C-terminus, followed by the LC3-interacting region (LIR). Blue and red dots, respectively, indicate the HSAN9-associated variants and AlphaFold predicted Rab5-binding defective mutants of TECPR2. The height of the line connected with the dots representing the mutants is only for clarity purposes. B) Interaction analysis of TECPR2 with the GTP-locked mutants of selected Rab proteins using yeast-two hybrid assay. Co-transformants were spotted on -Leu/-Trp to confirm viability and -Leu/-Trp/-His media to detect interactions. C) Yeast two-hybrid assay to detect interaction of TECPR2 with wild-type (WT)-, GTP (Q79L)-, and GDP (S34N)-locked mutants of Rab5. Co-transformants were spotted on -Leu/-Trp to confirm viability and -Leu/-Trp/-His media to detect interactions. D) Lysates of HEK293T cells co-transfected with FLAG-TECPR2 and GFP or GFP-Rab5 mutants were incubated with an anti-GFP nanobody. The precipitates were immunoblotted (IB) with the indicated antibodies. The anti-FLAG antibody detects a FLAG-TECPR2 band signal at ∼180 kDa (marked with an arrowhead) and a non-specific band signal as marked with an asterisk (*). E) Co- immunoprecipitation of endogenous TECPR2 with Rab5 was performed by incubating the HEK293T cell lysates with an anti-Rab5 antibody conjugated-resin or mouse IgG-conjugated- resin (as a control), and the resultant precipitates were IB with indicated antibodies. F) Yeast two- hybrid interaction analysis of different domain deletion mutants of TECPR2 with the GTP-locked form of Rab5. Co-transformants were spotted on -Leu/-Trp to confirm viability, and -Leu/-Trp/- His and -Leu/-Trp/-His/-Ade media to detect interactions. G) Recombinant MBP and MBP- TECPR2 (935-1411 aa) proteins immobilized on amylose resin were incubated with lysates of HEK293T cells expressing indicated GFP-Rab5 (WT/Q79L/S34N) proteins. The precipitates were IB with the indicated antibodies, and Ponceau S staining was done to visualize purified proteins. H) Recombinant GST and GST-Rab5 (WT/Q79L/S34N) proteins immobilized on glutathione- agarose beads were incubated with purified MBP-TECPR2 (935–1411 aa) protein. The precipitates were IB with an anti-MBP antibody, and Ponceau S staining was done to visualize the purified proteins. I) Yeast two-hybrid assay of AlphaFold predicted Rab5-binding defective mutants of TECPR2 with the GTP-locked (Q79L) form of Rab5. Co-transformants were spotted on -Leu/-Trp to confirm viability and -Leu/-Trp/-His media to detect interactions. J) Lysates of HEK293T cells co-transfected with GFP-Rab5 (Q79L) and FLAG-TECPR2 (WT) or with indicated AlphaFold predicted Rab5-binding defective mutants of TECPR2 were incubated with an anti-GFP nanobody. The precipitates were IB with the indicated antibodies. The values represent the densitometric analysis of the FLAG-TECPR2 band intensity normalized to the input and direct IP of GFP-Rab5 (Q79L). K) Yeast two-hybrid analysis of the indicated HSAN9- associated variants of TECPR2 with the Rab5 (Q79L) form. Co-transformants were spotted on - Leu/-Trp to confirm viability and -Leu/-Trp/-His media to detect interactions. L) Whole cell lysates of HEK293T cells transfected with the indicated FLAG-tagged TECPR2 mutants were IB with the indicated antibodies. The values represent the densitometric analysis of the FLAG- TECPR2 band intensity normalized to α-tubulin band intensity. M) Lysates of HEK293T cells co- transfected with GFP-Rab5 (Q79L) and FLAG-TECPR2 (WT) or with the indicated HSAN9- associated variants of TECPR2 were incubated with an anti-GFP nanobody. The precipitates were IB with the indicated antibodies. The values represent the densitometric analysis of the FLAG- TECPR2 band intensity normalized to the input and direct IP of GFP-Rab5 (Q79L).

Previous studies have shown that TECPR2 regulates the stability of its interaction partners, including HOPS and BLOC-1 complexes and the COPII coat subunits SEC24D-SEC23, although the functional significance of TECPR2 interaction with HOPS, and BLOC-1 complexes remains unclear (Stadel et al., 2015). TECPR2 localizes to membrane fractions containing ER, ERES, and ER-Golgi intermediate compartment (ERGIC) protein markers (Nalbach et al., 2023). TECPR2- depleted cells showed reduced functional ER exit sites (ERES) and delayed secretory cargo export from the ER to the Golgi. Consistent with this, a subsequent study showed that the plasma membrane proteome, secreted proteome, and lysosome composition were also altered upon the loss of TECPR2 expression (Nalbach et al., 2023; Stadel et al., 2015). TECPR2 has also been suggested to play a role in autophagosome biogenesis, via LC3C-mediated recruitment to the phagophore membrane wherein it may regulate membrane export from ER (Stadel et al., 2015). However, However, recent studies have suggested a role for TECPR2 in the later steps of autophagy, i.e., in autophagosome-lysosome fusion. Autophagosome accumulation has been reported in SPG49/HSAN9 patient fibroblasts, and reduced LC3 degradation has been observed in cells lacking TECPR2 expression (Fraiberg et al., 2021). In line with this, accumulation of autophagosomes was also observed in a tecpr2 knockout (tecpr2^-/-^) mouse model, indicating a defect in autophagosome-lysosome fusion (Tamim-Yecheskel et al., 2021).

Our attempts to characterize TECPR2 localization were inconclusive due to the lack of commercially available antibodies that detect the endogenous protein in immunofluorescence assays. Furthermore, consistent with previous findings, overexpression of epitope-tagged full-length TECPR2 showed a mostly cytosolic distribution (Fig. 2A) (Fraiberg et al., 2021; Stadel et al., 2015). As small G proteins of the Rab family are key regulators of vesicular transport that mediate recruitment of their downstream effectors on intracellular membranes, we tested whether a Rab protein could serve as an interaction partner for TECPR2 and regulate its membrane recruitment and function.

**Figure 2:**
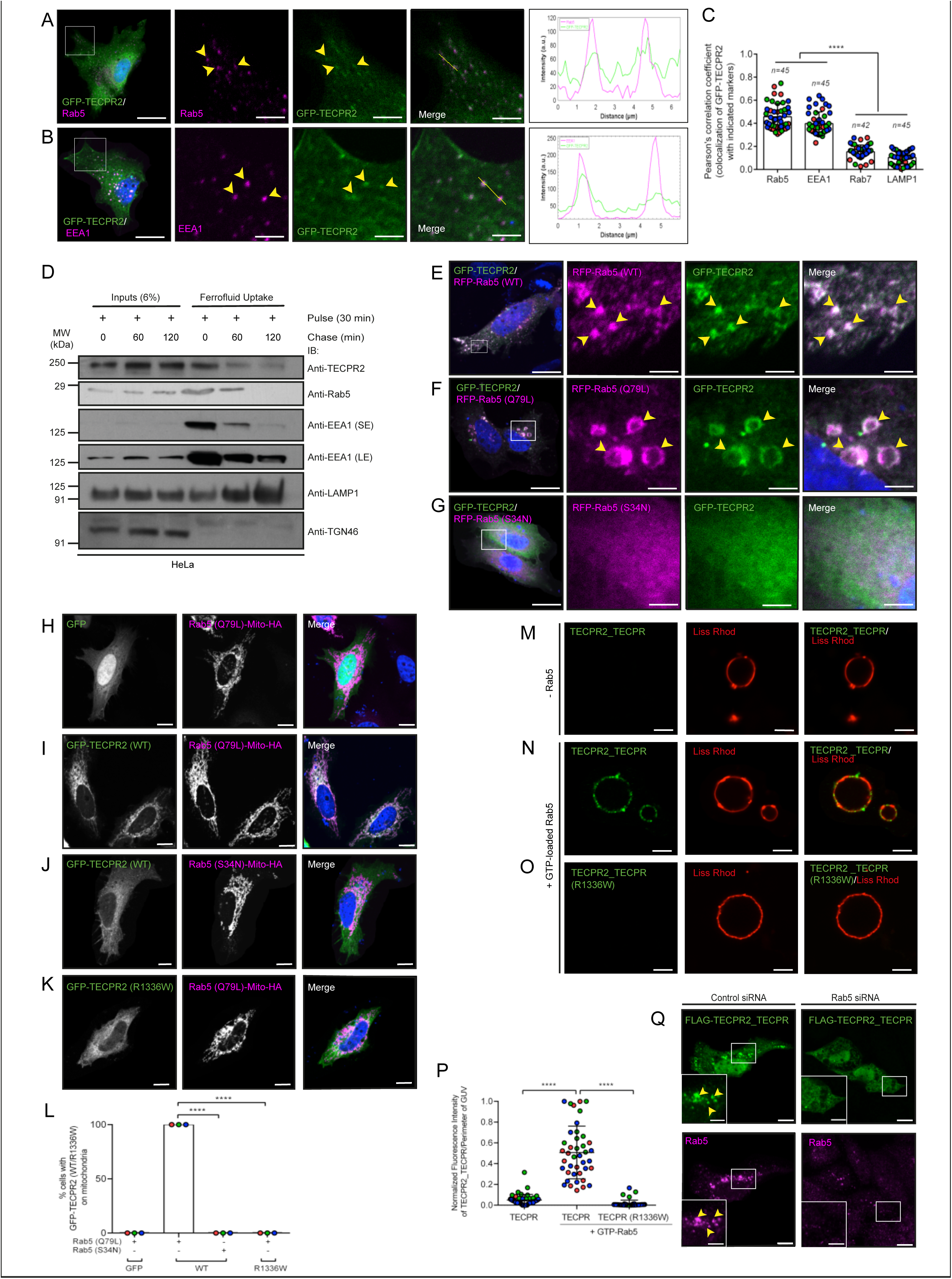
TECPR2 localizes to early endosomes in a Rab5-dependent manner, a process disrupted in the HSAN-associated R1336W variant. A and B) Representative confocal micrographs of HeLa cells transfected with GFP-TECPR2 and immunostained for endogenous Rab5 and EEA1. The line profiles on the right indicate fluorescence intensity along the yellow lines for both channels: GFP-TECPR2 (green) and Rab5 (magenta) (A) or EEA1 (magenta) (B). Arrowheads in the insets denote the colocalized pixels. Bars: 10 µm (main); 5 µm (inset). C) The quantification of the Pearson’s colocalization coefficient (PCC) for GFP-TECPR2 with the indicated markers is shown. n denotes the total number of cells examined in three independent experiments. Experiments are color-coded, and each dot represents the individual data points from each experiment (****p<0.0001; unpaired two-tailed Student’s t test). D) Ferrofluid uptake was performed in HeLa cells for 30 min (pulse) at 37°C, followed by chase for indicated time periods. The cells were homogenized at the indicated time points, and the ferrofluid-containing compartments were purified and IB for the indicated proteins. E-G) Representative confocal micrographs of HeLa cells co-transfected with GFP-TECPR2 and RFP-Rab5 (WT) (E), RFP-Rab5 (Q79L) (F), or RFP-Rab5 (S34N) (G). Arrowheads in the insets denote the colocalized pixels. Bars: 10 µm (main); 5 µm (inset). H-K) Immunofluorescence microscopy of HeLa cells co- transfected with GFP alone, GFP-TECPR2 (WT), or GFP-TECPR2 (R1336W) HSAN mutant with Rab5 (Q79L)-Mito-HA or Rab5 (S34N)-Mito-HA. Cells were fixed and immunostained with anti- HA antibodies. Scale bars: 10 μm. L) Quantification of the percentage of cells in which GFP- TECPR2 (WT) or R1336W mutant proteins were re-localized to mitochondria upon co- transfection with Mito-HA-tagged GTP (Q79L)- or GDP (S34N)-locked forms of Rab5 (****p< 0.0001; unpaired two-tailed Student’s t test). M-O) Confocal micrographs of Liss-Rhodamine- labeled GUVs incubated with purified TECPR2_TECPR (WT; 935–1411 aa) or with TECPR2_TECPR (R1336W; 935–1411 aa) in the absence (M) or presence (N-O) of purified GTP- bound Rab5. The GUVs were immunostained using an anti-TECPR2 antibody (shown in green) to determine the recruitment of TECPR2_TECPR onto GTP-Rab5-loaded GUVs. Scale bars: 10 μm. P) Quantification of fluorescence intensity of TECPR2_TECPR (WT or R1336W mutant) per perimeter of GUV in the absence or presence of GTP-bound Rab5 (****p<0.0001; unpaired two- tailed Student’s t test). Q) Confocal micrograph images of HeLa cells treated with control siRNA and Rab5 siRNA, followed by transfection with FLAG-TECPR2_TECPR. Cells were fixed and immunostained with anti-FLAG and anti-Rab5 antibodies. Arrowheads in the insets denote the colocalized pixels. Scale bars: 10 µm (main); 5 µm (inset).

Here, we report that TECPR2 is a Rab5 effector and that TECPR2 localizes to early endosomes in a Rab5-dependent manner. HSAN-associated missense mutation R1336W disrupts TECPR2 interaction with Rab5, thereby disrupting TECPR2 membrane recruitment. Depletion of TECPR2 led to perinuclear collapse of early and recycling endosomes and increased the overlap of sorting and degradative subdomain markers on early endosomes. TECPR2 depleted cells failed to efficiently recycle endocytosed α5β1 integrin receptors back to the cell surface, resulting in a reduced number of focal adhesions and, in turn, reduced cell spreading. TECPR2 interacts with and regulates the localization of β1 integrin cargo adaptor, sorting nexin (SNX) 17, to early endosomes. Concurrently, TECPR2 depletion, similar to SNX17 depletion, results in impaired retrieval of β1 integrin from the lysosomal degradation pathway. TECPR2-depleted cells also had reduced levels of Arp2/3-associated actin nucleation-promoting factor, WASH complex on Rab5- positive early endosomes. We propose that TECPR2 regulates the recruitment of endosomal sorting adaptors and their associated protein complexes on early endosomes, ensuring the retrieval of recycling cargo from a lysosomal degradation fate.

## Results

### TECPR2 directly binds with small G protein Rab5 via its C-terminal TECPR repeats

To test whether any of the Rab G proteins could interact with TECPR2, we performed yeast two-hybrid screening of TECPR2 with GTP-locked mutants of the selected candidate Rab proteins. As a positive control, we first confirmed the interaction of TECPR2 with its known binding partners, LC3B and the HOPS complex. As shown in Supplementary Fig. S1A, TECPR2 interacted with LC3B and HOPS subunits Vps41, Vps39, Vps33a, and Vps16, and very weakly with Vps11. Although Vps33a and Vps39 showed self-activation, their growth with TECPR2 WT was more than that with the pGADT7 empty vector. For Rab screening, we took 20 of the 70 mammalian Rab proteins in their constitutively active form, which are functionally characterized and are available in our laboratory (Fig. 1B). Interestingly, among these subsets of Rab proteins, Rab5 (Q79L) and Rab29 (Q67L) interacted with TECPR2. Interestingly, two previous studies had reported TECPR2 as a potential Rab5 interaction partner using proximity-dependent biotinylation approaches to identify Rab5 interactors (Gillingham et al., 2019; Go et al., 2021). Given that Rab5 is an extensively studied Rab protein, however, association of Rab5 with TECPR2 was not known, we wanted to confirm TECPR2 interaction with Rab5 and the significance of this interaction.

We validated that Rab5 binds to TECPR2 in its GTP-bound, but not GDP-bound conformation using constitutively dominant-active and -negative Rab5 point mutants (Q79L and S34N, respectively) in a yeast two-hybrid assay (Fig. 1C). Rab5 (Q79L) mutant bound to TECPR2 qualitatively more strongly than Rab5 (WT), as is known for other Rab effectors. We also performed co-immunoprecipitation assays that showed Rab5 interacts with full-length epitope-tagged TECPR2 in its WT and constitutively GTP-bound state, but only a weak interaction was observed with the constitutively GDP-bound Rab5 mutant (Fig. 1D). Finally, using co-immunoprecipitation assays, we confirmed that Rab5 interacts with TECPR2 under endogenous conditions (Fig. 1E).

Next, we investigated the binding region of TECPR2 required for its interaction with Rab5. We created TECPR2 mutants either lacking or containing the C-terminal TECPR repeats, i.e., TECPR2 (1-944 a.a.) and TECPR2 (935-1411 a.a.), and tested their ability to interact with Rab5 in a yeast two-hybrid assay. As shown in Fig. 1F, TECPR2 (935-1411 a.a.; henceforth labeled as TECPR2_TECPR) was both essential and sufficient for binding to Rab5 (Q79L), while no interaction was observed with the N-terminal region of TECPR2 (containing 1-944 a.a.). Notably, the TECPR2_TECPR fragment showed qualitatively more binding to Rab5 than TECPR2 (WT), suggesting that the N-terminal region may play an autoinhibitory role in regulating Rab5 binding. Next, we recombinantly expressed and purified MBP-tagged TECPR2_TECPR and incubated the purified protein with cell lysates expressing GFP-tagged WT and constitutively GTP-bound and GDP-bound forms of Rab5. As shown in Fig. 1G, the pulldown assay revealed that Rab5 interacts with the TECPR2_TECPR region of TECPR2 in a GTP-dependent manner.

To test whether there is direct binding of TECPR2_TECPR with GTP-bound Rab5, we performed a protein-protein interaction assay using recombinantly expressed and purified proteins. As shown in Fig. 1H, the TECPR2_TECPR fragment directly bound to GST-tagged Rab5 (Q79L) and a significantly weaker interaction was observed with GST-tagged Rab5 (S34N), confirming a physical interaction between the C-terminal region of TECPR2 and GTP-bound Rab5. Next, we employed ColabFold, a program that permits AlphaFold2-based predictions of protein complexes, to determine the binding interface residues between TECPR2_TECPR and Rab5 (Mirdita et al., 2022). Analysis of the predicted complex identified Met 1251, Ile 1256, and Glu 1299 as binding interface residues in TECPR2 (Supplementary Fig. S1B). These three residues are also highly conserved across evolution, increasing their likelihood of being relevant for TECPR2 function (Supplementary Fig. S1C). We used yeast two-hybrid and co-immunoprecipitation assays to test the significance of the TECPR2 I1256 and E1299 residues for binding to Rab5. As shown in Fig. 1I-J, mutation of E1299 but not I1256 to Alanine disrupted TECPR2 binding to Rab5, suggesting that residue E1299 might play a direct role in binding to Rab5.

HSAN-associated non-synonymous coding variants within the C-terminal TECPR repeats disrupt TECPR2 binding to Rab5 Previous studies have identified that the HSAN-associated missense clinical variants of TECPR2 map to the conserved residues within the N-terminal WD40 domain and the C-terminal TECPR repeats, suggesting that these residues regulate TECPR2 stability and/or function (Fig. 1A and Supplementary Fig. S1C) (Neuser et al., 2021). As we found that Rab5 directly interacts with the C-terminal TECPR repeats (935-1411 a.a.), we next evaluated the Rab5 binding potential of HSAN-associated clinical variants within this region. To this end, we created previously known non-synonymous coding variants in TECRP2: T903M, D1000Y, W1140G, and R1336W, all of which mark residues that are conserved across evolution (Supplementary Fig. S1C) (Neuser et al., 2021). AlphaMissense (AM), a technology designed to predict the pathogenicity of missense variants, showed that the TECPR2 clinical variants D1000Y, W1140G, and R1336W had high AM scores and were categorized as pathogenic variants, whereas T903M had a low AM score and was categorized as a benign variant (Supplementary Table I) (Cheng et al., 2023; Tordai et al., 2024). Consistent with the AM predictions, TECPR2 D1000Y, W1140G, and R1336W variants showed highly reduced or no binding to Rab5, whereas the T903M variant continued to interact with Rab5, similar to the WT protein (Fig. 1K). We noted that the TECPR2 variants D1000Y, W1140G, and R1336W had highly reduced expression in cell lysates, compared to the WT protein, while T903M variant expression was similar to WT (Fig. 1L). The W1140G variant of TECPR2 did not show a band corresponding to the WT molecular weight (∼180 kDa) but showed an unexplained downward mobility shift; thus, this variant was not used for further analysis. Consistent with the yeast two-hybrid assay, D1000Y and R1336W variants did not interact with Rab5, whereas the T903M variant continued to interact with Rab5 in a co-immunoprecipitation assay (Fig. 1M). Notably, a previous study showed that the non-synonymous coding variant R1337W in canine TECPR2 (residue R1336 in humans) was associated with juvenile-onset neuroaxonal dystrophy (NAD) in Spanish water dogs (Hahn et al., 2015). Our results suggest that while the R1336W mutation may destabilize TECPR2, leading to its reduced expression, the minimally expressed version is also defective in Rab5 binding, which may further explain the association of this variant with HSAN9 disease.

TECPR2 localizes to early endosomes in a Rab5-dependent manner, a process disrupted in HSAN-associated R1336W variant Previous studies have shown that epitope-tagged TECPR2 localizes to the cytosol, and partial overlap has been reported with ER markers, including COPII coat proteins SEC24D, SEC31, and VAPB (Nalbach et al., 2023; Stadel et al., 2015). We were unable to detect endogenous TECPR2 by immunofluorescence as the commercially available anti-TECPR2 antibodies did not recognize the protein in HeLa cells. Thus, we analyzed the localization of epitope-tagged constructs, ensuring that only weak to moderate TECPR2-expressing cells were taken for analysis. Interestingly, we observed that GFP-tagged TECPR2 (GFP-TECPR2), while having a mostly cytosolic distribution, also showed punctate localization, especially in weakly transfected cells (Fig. 2A). Immunostaining of GFP-TECPR2 expressing cells with anti-Rab5 antibodies showed that these GFP-TECPR2 punctae were indeed positive for Rab5 (Fig. 2A; Pearson’s colocalization coefficient (PCC) quantification is shown in Fig. 2C). The TECPR2 punctae were also positive for the early endosomal marker and Rab5 effector-EEA1 (Fig. 2B-C), while little or no colocalization was observed with late endosomal/lysosomal markers-Rab7 and LAMP1 (Supplementary Fig. S2A-B). Next, to analyze whether TECPR2 colocalizes with the ER tubular network, we co-expressed TECPR2 with the ER membrane contact site protein VAPB, previously shown to interact with TECPR2. GFP-TECPR2 in live cells showed a cytosolic localization with few punctate structures and was not observed on the VAPB-positive ER tubular network; however, a few TECPR2-positive vesicles were colocalized with VAPB (Supplementary Video S1). We found that TECPR2-positive vesicles were associated with ER tubules, and few were co-migrating in association with the ER (Supplementary Video S1). Indeed, the association of TECPR2 punctae and co-migration along ER tubules is reminiscent of Rab5-positive early endosome motility along ER tubules (Friedman et al., 2013).

To test whether endogenous TECPR2 localizes to early endosomal fractions, we used an approach wherein cells were incubated with 10 nm paramagnetic particles (Ferrofluid [FF]), and endocytosed paramagnetic particle-containing vesicles were isolated from cell homogenates at different time points post-incubation (Schleinitz et al., 2023; Walia et al., 2024). As shown in Fig. 2D, the early endosomal proteins, including Rab5 and EEA1, were enriched in the pulse-only sample, and their levels steadily decreased upon chase for 60 min and 120 min, whereas the late endocytic proteins Rab7 and LAMP1 showed a reverse trend, indicating maturation of the FF- loaded endosomes with increasing time points of chase. We did not observe the presence of Golgi marker TGN46 in any of the fractions, indicating that the fractions were endocytic compartments. We found that TECPR2 was enriched in the early endosomal fractions as compared to the late endosomal fractions, strengthening our conclusion that TECPR2 at the endogenous expression level in HeLa cells localizes to early endosomes (Fig. 2D).

Rab proteins direct vesicular transport by recruiting effectors to target membranes and orchestrating vesicle motility, tethering, and fusion. Next, we employed several independent approaches to test whether GTP-bound Rab5 recruits TECPR2 to early endosomes. First, co-expression of Rab5 WT and its constitutively dominant active mutant (Q79L) led to dramatic membrane recruitment of TECPR2, whereas TECPR2 was completely cytosolic in cells co-expressing the constitutively dominant negative mutant (S34N) (Fig. 2E-G). Importantly, in contrast to TECPR2 (WT), which was recruited on early endosomes upon co-expression with Rab5, the HSAN9 variants of TECPR2 that weakly interacted with Rab5 (D1000Y) or did not interact (W1140G and R1336W) remained cytosolic in the presence of Rab5 (Supplementary Fig. S2C-I). Consistent with its binding to Rab5, the HSAN9-associated variant T903M continued to colocalize with Rab5 (Supplementary Fig. S2D). The AlphaFold2-predicted Rab5 binding-defective mutant, i.e., TECPR2 (E1299A), also showed a highly reduced overlap with Rab5 endosomes, suggesting that Rab5 binding is required for TECPR2 membrane localization (Supplementary Fig. S2H-I).

Second, we employed a recently reported MitoID method for Rab effector and regulator identification to test whether Rab5 is sufficient to direct TECPR2 membrane localization (Gillingham et al., 2019). To this end, we expressed TECPR2 with constitutively dominant-active and dominant-negative Rab5 constructs fused to a mitochondrial-targeting sequence (Rab5 (Q79L)-mito and Rab5 (S34N)-mito). As a control, we first verified that no mitochondrial recruitment of the GFP-tag by itself was observed in the presence of Rab5 (Q79L)-mito (Fig. 2H- L). GFP-TECPR2 was recruited to mitochondria in cells co-expressing Rab5 (Q79L)-mito, while no recruitment was observed with Rab5 (S34N)-mito (Fig. 2I-J and 2L). Further, HSAN9- associated TECPR2 variant R1336W was not recruited to mitochondria in the presence of Rab5 (Q79L)-mito, indicating that GTP-bound Rab5 interacts with and recruits TECPR2 to intracellular membranes, and this process is disrupted upon HSAN9-associated R1336W mutation (Fig. 2K- L).

Next, we employed an in vitro minimal reconstitution assay to test whether Rab5 is sufficient to direct TECPR2 on membranes. To this end, constitutively active Rab5 (Q79L) was anchored on giant unilamellar vesicles (GUVs) through a covalent bond between the C-terminal cysteine residue of Rab5 and a maleimide-conjugated lipid in GUV, as recently described (Tremel et al., 2021). We confirmed Rab5 loading by immunostaining the Liss-Rhodamine-labeled GUV membranes with anti-Rab5 antibodies (Supplementary Fig. S2J). Next, we incubated the purified Rab5 binding fragment of TECPR2, i.e., TECPR2_TECPR (WT) and the Rab5 binding-defective version (TECPR2_TECPR (R1336W)), with GUVs containing GTP-loaded Rab5 and evaluated TECPR2 recruitment using immunostaining with anti-TECPR2 antibodies. We found that TECPR2_TECPR (WT) was recruited to GUVs in the presence of GTP-loaded Rab5 and no recruitment was detected on GUVs without Rab5 (Fig. 2M-N and 2P). Importantly, even in the presence of GTP-loaded Rab5, TECPR2_TECPR (R1336W) was not recruited to GUVs, confirming previous observations that this HSAN9-associated TECPR2 variant does not interact with Rab5 and therefore, fails to localize to membranes (Fig. 2O-P).

We found that the epitope-tagged TECPR2_TECPR fragment when expressed in cells was mostly punctate, and almost all punctae were colocalized with endogenous Rab5 (Supplementary Fig. S2L). Notably, in cells with moderate to high TECPR2_TECPR expression, the Rab5 effector EEA1 appeared to be cytosolic, indicating that TECPR2_TECPR competes for Rab5 binding with other endogenous effectors (Supplementary Fig. S2M). Consistent with the essential role of Rab5 in mediating TECPR2 membrane localization, depletion of all three isoforms of Rab5 (A, B, and C) led to a relocalization of the TECPR2_TECPR fragment from the membrane to the cytosol, corroborating that Rab5-binding is required for TECPR2 membrane association (Fig. 2Q; knockdown efficiency was >90%, as confirmed by immunoblotting and is shown in Supplementary Fig. S2K). Taken together, we conclude that TECPR2 is a Rab5 effector and the C-terminal TECPR repeats are essential and sufficient for Rab5 binding and, consequently, for TECPR2 recruitment to early endosomes. Importantly, impaired Rab5 binding and lack of membrane localization explain the loss-of-function phenotype of the HSAN9-associated TECPR2 R1336W variant.

TECPR2 localizes to and maintains the peripheral distribution of early and recycling endosomes To gain insights into TECPR2 function, we next decided to visualize the dynamics of TECPR2- positive early endosomes in live cells. We imaged only weakly transfected cells for this analysis, as cells with high TECPR2 expression showed large and aberrant clustering of Rab5-positive vesicles in the perinuclear region (data not shown). Interestingly, we observed two different populations of TECPR2 and Rab5 endosomes: peripheral, smaller endosomes that showed bidirectional motility near the cell surface, and relatively larger perinuclear ring-shaped endosomes that were less mobile (Supplementary Video S2). We also found that TECPR2- positive short tubules emanating from Rab5-positive endosomes occasionally undergo fission (Fig. 3A-B and Supplementary Video S3). TECPR2-positive endosomes also exhibited tethering and homotypic fusion (Supplementary Video S3). Intensity profile analysis over time showed that TECPR2-positive endosomes occasionally underwent both fission and fusion (Fig. 3C, and Supplementary Fig. S3). In a subset of transfected cells, we also observed TECPR2-positive, relatively long, and stable tubules (Supplementary Video S4). Notably, we did not observe tubulation of TECPR2 endosomes in fixed cells, likely because of poor preservation of the tubular structures upon fixation. As expected, owing to its lack of Rab5 binding, the HSAN9-associated TECPR2 (R1336W) point-mutant did not show localization to vesicular or tubular membranes when co-expressed with Rab5 (Supplementary Video S5). Next, we analyzed whether TECPR2 vesicular and tubular endosomes were accessible to endocytic cargo. To this end, we incubated TECPR2 and Rab5 co-expressing cells with labeled transferrin (Tfn-568), a cargo that recycles back to the cell surface along with its receptor. At 5 min post-internalization, Tfn-568 was present in TECPR2-positive endosomes. We also observed Tfn-568 localized on TECPR2-positive short and dynamic tubular endosomes, which eventually underwent fission (Fig. 3D and Supplementary Video S6).

**Figure 3:**
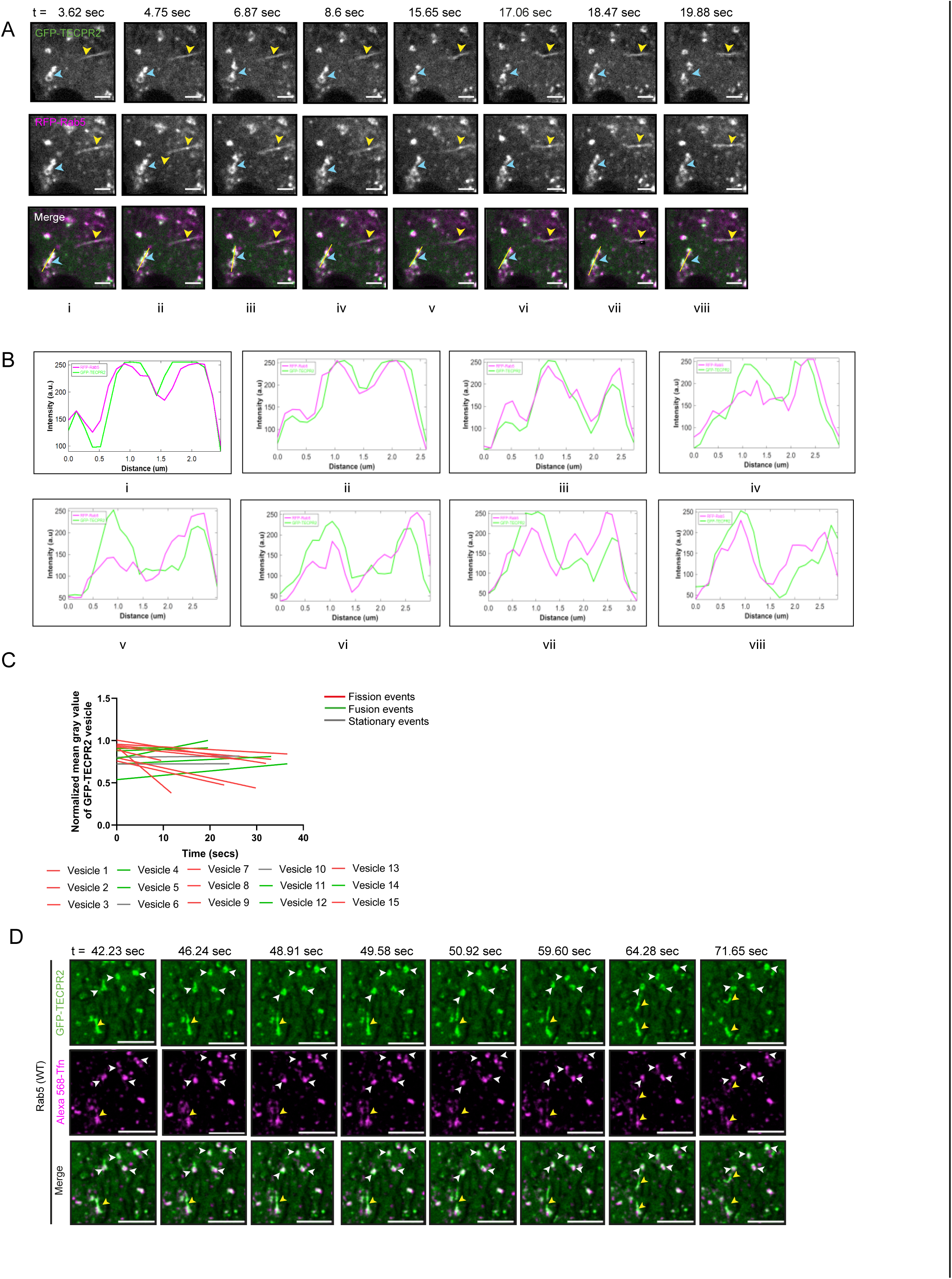
TECPR2 localizes to and maintains the peripheral distribution of early and recycling endosomes. A and B) Time-lapse images of HeLa cells expressing GFP-TECPR2 (green) and RFP-Rab5 (magenta) (see Supplementary Video S3). The blue arrowheads indicate GFP-TECPR2- and RFP-Rab5-positive vesicles forming short tubules and undergoing fission to form two separate endosomes. The yellow arrowheads indicate GFP-TECPR2-positive tubules. Scale bars: 5 µm. In (B), the line-scan analysis of the fluorescence intensity of GFP-TECPR2 and RFP-Rab5 across the indicated endosomes is shown. C) A graph showing fluorescence intensities of GFP-TECPR2-positive vesicles during fission and fusion processes. Linear regressions are shown for n=15 vesicles. Slightly positive and negative slopes are marked in green and red, respectively, and the flat lines are indicated in grey. D) Time-lapse images of HeLa cells expressing GFP-TECPR2 (green) and untagged-Rab5 and incubated with Alexa 568-Tfn ligand (magenta) (see Supplementary Video S6). The white arrowheads indicate GFP-TECPR2 vesicles containing Alexa 568-Tfn undergoing elongation and eventually fission. The yellow arrowheads indicate Alexa 568-Tfn ligand containing GFP-TECPR2-positive tubules undergoing fission event. Scale bars: 5 µm.

To analyze TECPR2 function in HeLa cells, we next employed an siRNA-based approach to reduce TECPR2 protein expression and investigate the status of early and recycling endosomal proteins (efficiency of knockdown as confirmed by western blotting was >90%, as shown in Supplementary Fig. S4A). Notably, we observed a striking loss of Rab5- and EEA1-positive endosomes from the cell periphery and relocalization of early endosomes to the perinuclear region (Fig. 4A-B). A prominent cytosolic pool of Rab5 was also visible, whereas EEA1-positive endosomes appeared to be enlarged in TECPR2-depleted cells (Fig. 4B and 4D). Surprisingly, TECPR2-depleted HeLa cells had significantly reduced cell spreading compared to control cells (compare cell spreading in Fig. 4 panel A versus panel B). Therefore, to quantify the distribution of Rab5 and EEA1, we measured the fractional distance of Rab5- and EEA1-positive early endosomes from the cell center, which is independent of the cell area (Jongsma et al., 2016). Indeed, we observed a significant shift in early endosome distribution towards the perinuclear region (Fig. 4E). The effect on early endosome distribution was rescued by the re-expression of TECPR2 (WT), indicating that this phenotype was specifically due to TECPR2 depletion (Fig. 4C and 4E).

**Figure 4:**
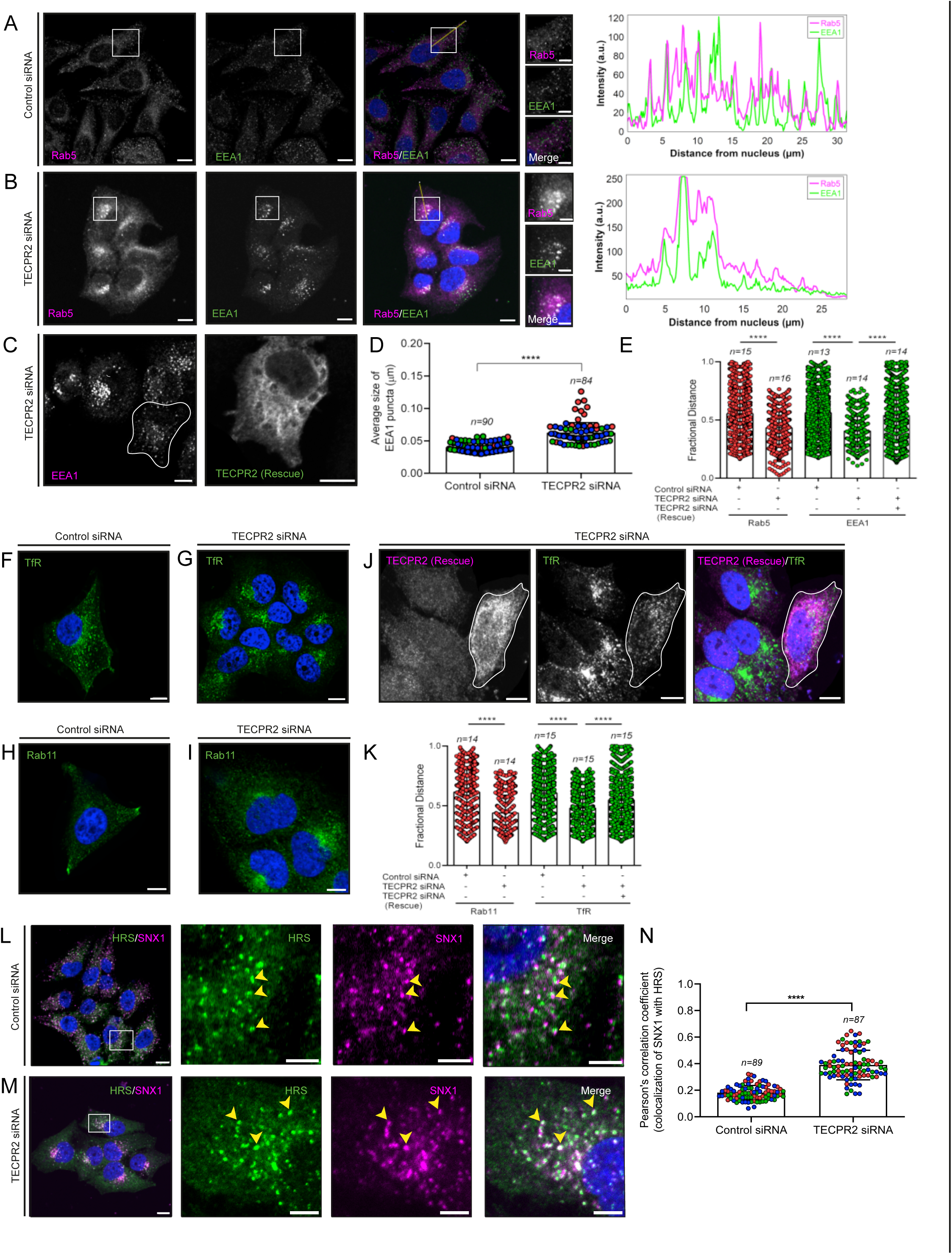
TECPR2 localizes to and maintains the peripheral distribution of early and recycling endosomes. A and B) Representative confocal micrographs of HeLa cells treated with the indicated siRNA, followed by immunostaining with anti-Rab5 and anti-EEA1 antibodies. On the right, the line-scan analysis of the fluorescence intensity of EEA1 (green) and Rab5 (magenta) along the yellow line is shown. Scale bars: 10 µm (main); 5 µm (inset). C) Representative confocal micrograph of HeLa cells treated with TECPR2 siRNA and transfected with the siRNA-resistant FLAG-tagged TECPR2 rescue construct (highlighted with a white line), followed by immunostaining with anti-FLAG and EEA1 antibodies. Scale bars: 10 µm. D) Quantification of EEA1 puncta size in HeLa cells treated with control siRNA and TECPR2 siRNA. The values plotted are the mean ± SD from three independent experiments. Experiments are color-coded, and each dot represents the individual data points from each experiment (****p<0.0001; unpaired two- tailed Student’s t test). E) The graph represents quantification of the distribution (shown as fractional distance) of Rab5-positive endosomes and EEA1-positive endosomes in HeLa cells with indicated siRNA treatment. The values plotted are mean ± S.D., and the total number of cells analyzed is indicated on the graph (****p<0.0001; unpaired two-tailed Student’s t test). F-I) Representative confocal micrographs of HeLa cells treated with the indicated siRNA followed by immunostaining with anti-TfR (F-G) and anti-Rab11 (H-I) antibodies. Scale bars: 10 µm. J) Representative confocal micrograph of HeLa cells treated with TECPR2 siRNA and transfected with siRNA-resistant FLAG-tagged TECPR2 rescue construct (highlighted with a white line), followed by immunostaining with anti-FLAG and anti-TfR antibodies. Scale bars: 10 µm. K) The graph represents quantification of the distribution (shown as fractional distance) of Rab11-positive endosomes and TfR-positive endosomes in HeLa cells upon indicated siRNA treatment. The values plotted are mean ± S.D., and the total number of cells analyzed is indicated on the graph (****p<0.0001; unpaired two-tailed Student’s t test). L and M) Representative confocal images of HeLa cells treated with either control siRNA or TECPR2 siRNA and immunostained with anti- SNX1 and anti-HRS antibodies. The yellow arrowheads denote co-localized pixels. Scale bars: 10 µm (main); 5 µm (inset). N) The graph shows quantification of the Pearson’s colocalization coefficient of SNX1 with HRS in HeLa cells treated with either control or TECPR2 siRNA. Experiments are color-coded, and each dot represents the individual data points from each experiment. The total number of cells analyzed is indicated on the top of each data set graph (****p<0.0001; unpaired two-tailed Student’s t test).

We also analyzed the distribution of other Rab5 effectors, Rabenosyn-5 and APPL1, in TECPR2-depleted cells. APPL1 endosomes form a distinct Rab5-positive sorting compartment that exchanges cargo via fusion and fission with EEA1 positive endosomes (Kalaidzidis et al., 2015). Notably, while both Rabenosyn-5- and APPL1-positive compartments accumulated in the perinuclear region, a subpopulation of these endosomes continued to remain near the plasma membrane in TECPR2-depleted cells (Supplementary Fig. S4E-F). We confirmed the effect on early endosome size and distribution upon TECPR2 depletion by treating cells with two different siRNA oligo sequences that were efficient at TECPR2 knockdown, as well as in other human cell lines, including HEK293T and hTERT RPE-1 cells (Supplementary Fig. S4B-D and S4G-K). The decrease in the cell spreading area was also evident upon TECPR2 depletion using different siRNA oligos in HeLa cells, as well as in hTERT RPE-1 and HEK293T cell lines (Supplementary Fig. S4G-K). We also assessed Rab5 protein levels in TECPR2-depleted cells, as TECPR2 knockdown reduced Rab5 membrane association and partially localized it to the cytosol. Indeed, Rab5 levels were reduced in total cell lysates following TECPR2 depletion in multiple cell lines (Supplementary Fig. S4L). Our results are consistent with a previous study showing increased degradation of TECPR2 partner proteins upon its depletion (Stadel et al., 2015).

As TECPR2-positive endosomes were accessible to endocytosed receptor such as transferrin receptor (TfR) which undergoes constitutive recycling, we next determined whether TECPR2 depletion also affects recycling endosomes. TECPR2-depleted cells showed a striking relocalization of TfR and recycling endosomal marker protein-Rab11 from the cell periphery to the perinuclear region (Fig. 4F-I and 4K, and Supplementary Fig. S4M and S4O). TfR distribution was rescued by the re-expression of TECPR2, indicating that this phenotype was specifically due to TECPR2 depletion (Fig. 4J and 4K). We also found that SNX1, which marks the early endosomal sorting vesicles that mediate cargo trafficking towards the TGN, also relocalized and collapsed in the perinuclear region. (Fig 4L-M and Supplementary Fig. S4N and S4P). Moreover, SNX1 overlap with the ESCRT-0 subunit HRS, which mediates cargo sorting to intraluminal vesicles for degradation, was significantly increased in TECPR2-depleted cells (Fig 4L-N). We also observed dramatic enlargement of late endosomes/lysosomes marked by LAMP1 in TECPR2-depleted cells (Supplementary Fig. S4Q and S4R).

Thus, TECPR2-positive vesicles and tubules are accessible to recycling cargo, such as transferrin, and TECPR2 depletion results in perinuclear collapse of the early and recycling endosomal proteins. TECPR2-depleted cells showed enhanced colocalization of SNX1, a sorting adaptor, and HRS, which binds ubiquitinated cargo for subsequent endolysosomal degradation. These findings suggest that TECPR2 depletion might shuttle membranes and their contents from recycling towards degradation, causing a higher influx of membranes and cargo towards the late endosomal and endolysosomal compartments.

TECPR2 mediates retrieval of β1 integrin receptor from lysosomal degradation To investigate TECPR2 role in cargo recycling, we first tested whether constitutive recycling of transferrin is altered by TECPR2 depletion. We noted that steady-state surface levels of TfR were modestly but consistently reduced by ∼20% in TECPR2-depleted cells (Supplementary Fig. S5A). In line with this, we observed reduced uptake of Tfn-568 in TECPR2-depleted cells. Notably, when normalized to the “pulse-only” signal intensity, we found a modest but significant decrease of ∼20% in Tfn recycling upon TECPR2 depletion at 20 and 30 min of chase (Supplementary Fig. S5B). Moreover, Tfn-568-containing recycling endosomes clustered in the perinuclear region, similar to the observed steady-state distribution of TfR and Rab11 in TECPR2- depleted cells (Supplementary Fig. S5C-D and also see Fig. 4G and 4I). Notably, after 60 min of chase, Tfn recycling was similar to that of the control in TECPR2-depleted cells, suggesting that TECPR2 does not play a major role in TfR recycling (Supplementary Fig. S5B).

A dramatic and consistent phenotype of TECPR2 knockdown in multiple cell lines was that these cells showed reduced spreading compared to control cells (Fig. 5A-B). Quantification of the surface area of control and TECPR2 depleted cells at steady state revealed a ∼1.6-fold decrease in the surface area of TECPR2-depleted cells (Fig. 5C). As cell spreading is mediated by integrin receptor-dependent focal adhesions and extracellular matrix connections, we quantified the number of focal adhesions upon TECPR2 depletion. Strikingly, the number of focal adhesions (labeled by Paxillin) was significantly reduced in TECPR2-depleted cells (Fig. 5D-F). Consistent with this, we found a significant decrease in the spreading and focal adhesion number of trypsinized TECPR2-depleted cells after reseeding for 90 min on fibronectin-coated coverslips, compared to the control (Fig. 5G-I). This led us to investigate whether TECPR2 regulates integrin trafficking, specifically, integrin recycling. To this end, we assessed the surface levels of active and inactive Itgβ1 (integrin β1) using flow cytometry in control and TECPR2-depleted cells. As shown in Supplementary Fig. S5E, the surface levels of active Itgβ1 were reduced by ∼ 25% upon TECPR2 depletion. We also found that the surface levels of EGFR, which, in the absence or low concentration of its ligand, are mostly sorted from early endosomes for constitutive recycling (Bakker et al., 2017), were strikingly reduced in TECPR2 depletion (Supplementary Fig. S5F). In contrast, surface levels of LAMP1 and cation-independent mannose-6-phosphate receptor (CI- M6PR), which are sorted from early endosomes towards intracellular compartments, i.e., late endosomes/lysosomes and the trans-Golgi network (TGN), respectively, were unaffected in TECPR2-depleted cells (Supplementary Fig. S5G-H).

**Figure 5:**
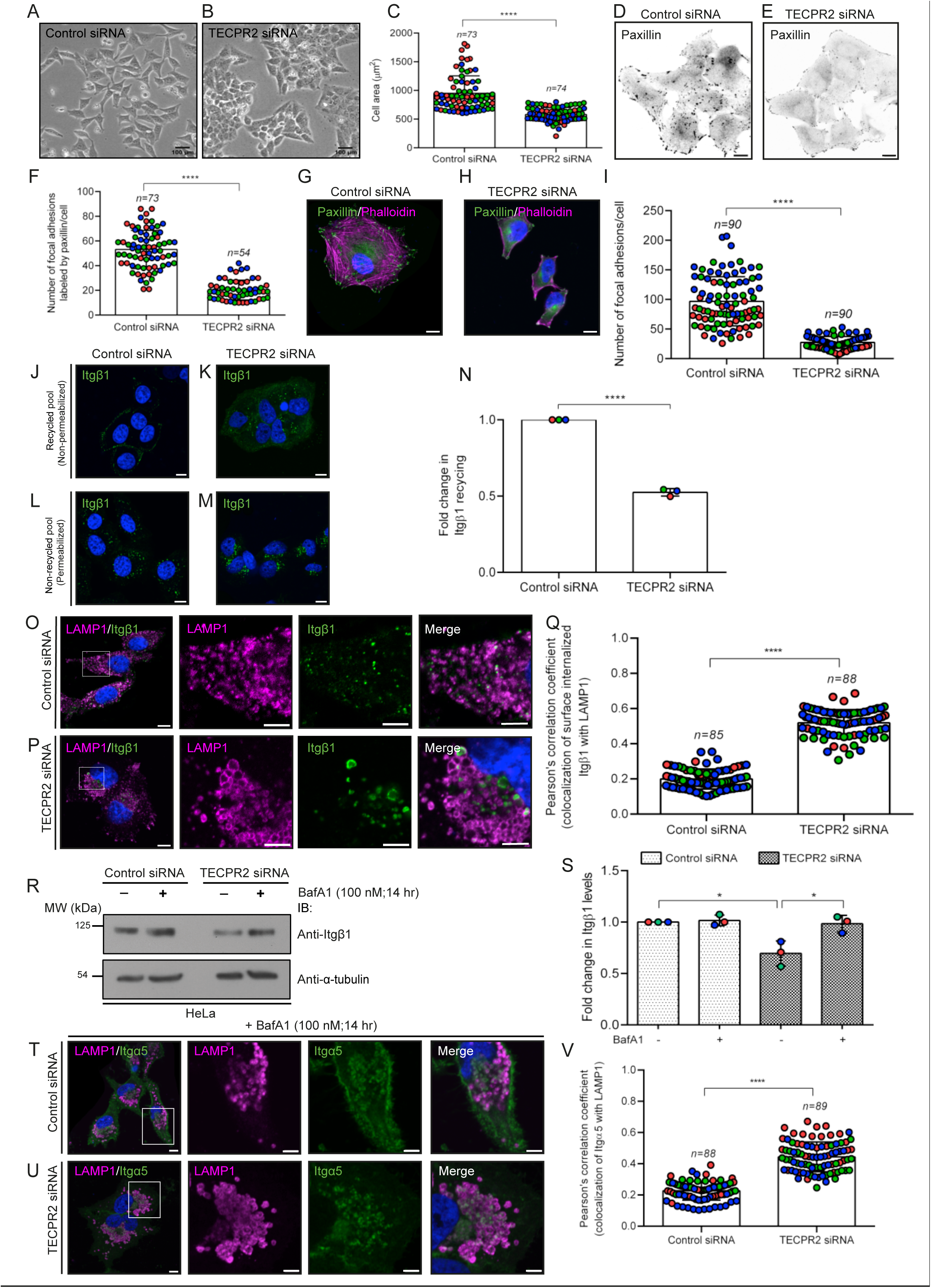
TECPR2 mediates retrieval of β1 integrin receptor from lysosomal degradation. A-C) Phase-contrast microscopy images of HeLa cells treated with control siRNA and TECPR2 siRNA. As shown, upon silencing of TECPR2 expression, reduction in cell spreading was observed. Scale bar: 100 µm. The quantification of cell area in control and TECPR2 siRNA-treated HeLa cells is shown in (C). n denotes the total number of cells examined over three independent experiments. Experiments are color-coded, and each dot represents the individual data points from each experiment (****p<0.0001; unpaired two-tailed Student’s t test). D-F) Representative confocal images of HeLa cells treated with control siRNA and TECPR2 siRNA and immunostained with an anti-paxillin antibody. The inverted images are shown for better visualization of focal adhesions. Scale bar: 10 µm. The quantification of the number of focal adhesions labeled by paxillin in HeLa cells treated with the indicated siRNA is shown in (F). n denotes the total number of cells examined over three independent experiments. Experiments are color-coded, and each dot represents the individual data points from each experiment (****p<0.0001; unpaired two-tailed Student’s t test). G-I) Representative confocal images of HeLa cells treated with indicated siRNA, followed by trypsinization and re-seeding on fibronectin- coated coverslips. After 90 min post-seeding, cells were fixed and stained with anti-paxillin antibody to determine cell spread area. Scale bar: 10 µm. In (I), quantification of the number of focal adhesions per cell labeled by paxillin is shown. n represents the total number of cells examined over three independent experiments. Experiments are color-coded, and each dot represents the individual data points from each experiment (****p<0.0001; unpaired two-tailed Student’s t test). J-N) A recycling assay with antibodies against active (12G10) Itgβ1 was performed on HeLa cells treated with indicated siRNA to determine the surface (recycled) pool (J-K) and internal (non-recycled) pool (L-M) of Itgβ1. Scale bar: 10 µm. In (N), the ratio of surface to internal active Itgβ1 pools in control and TECPR2-depleted HeLa cells is shown (****p<0.0001; unpaired two-tailed Student’s t test). O-Q) A recycling assay with antibodies against active (12G10) Itgβ1 was performed on HeLa cells treated with indicated siRNA, and localization of the internal (non-recycled) pool of Itgβ1 to lysosomes (LAMP1-positive) was visualized by confocal imaging. Scale bars: 10 µm (main); 5 µm (inset). In (Q), the Pearson Correlation Coefficient (PCC) of the internal (non-recycled) pool of active Itgβ1 with the lysosomal marker LAMP1 was measured in HeLa cells treated with control siRNA and TECPR2 siRNA. n denotes the total number of cells examined over three independent experiments. Experiments are color-coded, and each dot represents the individual data points from each experiment (****p<0.0001; unpaired two-tailed Student’s t test). R and S) Western blot analysis of HeLa cell lysates transfected with indicated siRNA, and treated with or without bafilomycin A1 (BafA1; 100 nM) for 14 hr, followed by IB with anti-Itgβ1 (9EG7) and anti-α-tubulin antibodies. In (S), quantitative western blot analysis of the effect of TECPR2 depletion on total levels of active Itgβ1 in HeLa cells upon BafA1 treatment is shown (*p=0.0121 for control and TECPR2 siRNA without BafA1 treatment; *p=0.0264 for TECPR2 siRNA with and without BafA1 treatment; unpaired two-tailed Student’s t test). T-V) Representative confocal images of HeLa cells transfected with the indicated siRNA and treated with BafA1 (100 nM) for 14 hr. Cells were fixed and immunostained with antibodies to Itgα5 and LAMP1. Scale bars: 10 µm (main); 5 µm (inset). In (V), the PCC of Itgα5 with the lysosomal marker LAMP1 was determined in HeLa cells transfected with indicated siRNA and treated with BafA1 (100 nM) for 14 hr. n denotes the total number of cells examined over three independent experiments. Experiments are color-coded, and each dot represents the individual data points from each experiment (****p<0.0001; unpaired two- tailed Student’s t test).

Next, we employed an antibody-based recycling assay to investigate the fate of endocytosed active Itgβ1 in the control and TECPR2-depleted cells. We found that the surface to internalize active Itgβ1 levels at 30 min of chase showed a ∼1.8-2-fold decrease upon TECPR2 depletion compared to the control (Fig. 5J-N). Previous studies have shown that blocking the retrieval of Itgβ1 receptors from early endosomes leads to their trafficking and degradation in endolysosomes (Steinberg et al., 2012). To this end, we next permeabilized and immunostained control and TECPR2-depleted cells with the late endosomal and lysosomal marker LAMP1 to investigate whether internalized active Itgβ1 receptors were trafficked to lysosomes instead of recycling to the cell surface. Notably, colocalization of internalized active Itgβ1 with LAMP1 was significantly increased in TECPR2-depleted cells (Fig. 5O-Q). To further investigate whether integrin stability at steady state is altered in TECPR2-depleted cells, we measured the total levels of Itgβ1 in control and TECPR2-depleted cell lysates. Indeed, we found reduced levels of Itgβ1 in TECPR2 knockdown cells, which was rescued by overnight treatment of cells with the lysosomal V-ATPase inhibitor BafA1 (Bafilomycin A1) (Fig. 5R and 5S). We corroborated these findings by immunostaining, where significantly increased colocalization of Itgα5 and LAMP1 was observed in TECPR2-depleted cells but not in control cells upon BafA1 treatment. (Fig. 5T-V). Taken together, these observations suggest that TECPR2 is required for efficient retrieval of integrin from the lysosomal degradation pathway and integrin-mediated cell spreading. TECPR2 may also be required for the constitutive recycling of EGFR, but this requires further exploration in future studies.

TECPR2 interacts with SNX17 and regulates its localization to Rab5-positive endosomes Previous studies have shown that knockdown of SNX17, the sorting adaptor that binds to the NPXY sorting motif on Itgβ1 and Itgβ5 tails, leads to a defect in β1 integrin retrieval from the lysosomal degradation pathway, which is similar to the phenotype we observed upon TECPR2 depletion (Cullen and Steinberg, 2018; Steinberg et al., 2012). Notably, SNX27, which is closely related to SNX17 (both members of the SNX-FERM family of sorting nexins), has been reported as a potential binding partner of TECPR2 (Nalbach et al., 2023; Stadel et al., 2015). Thus, we next investigated whether TECPR2 and SNX17 play a role in the same pathway of β1 integrin recycling. We found that mature β1 integrin (mol. wt. ∼130 KDa) was degraded to a similar extent in SNX17 or SNX17 and TECPR2 co-depleted cells, suggesting that TECPR2 and SNX17 do not compensate for each other’s loss and likely play a role in the same pathway of β1 integrin recycling (Fig. 6A). We also found an increase in the total levels of the immature unglycosylated Golgi resident form of Itgβ1 (∼100 KDa) in SNX17-depleted and SNX17 and TECPR2 co-depleted cells (Fig. 6A). It is unclear whether the increase in immature Itgβ1 is due to enhanced synthesis of Itgβ1 in these cells.

**Figure 6:**
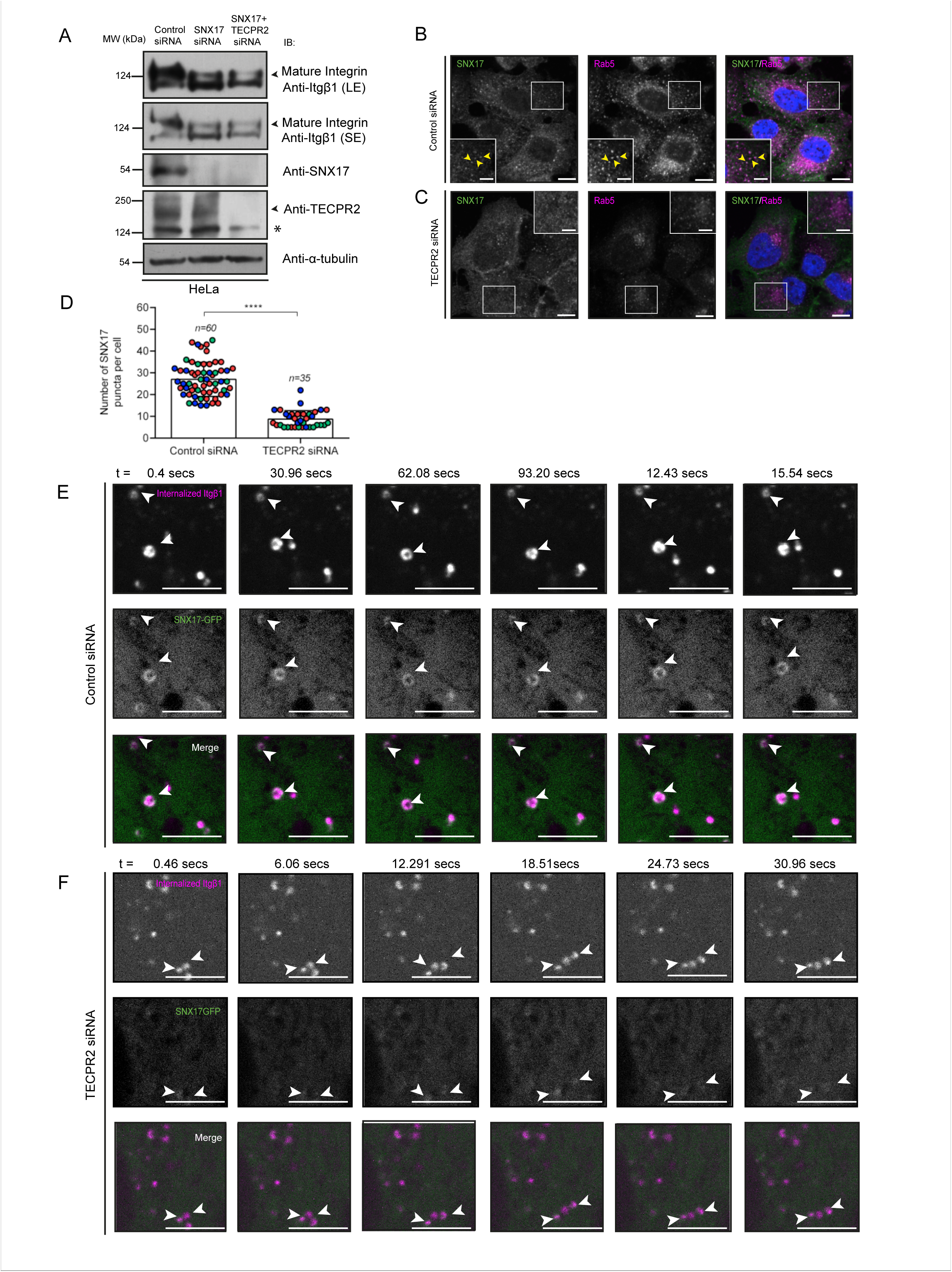
TECPR2 regulates sorting adaptor SNX17 localization on endocytosed β1-integrin- containing endosomes. A) Western blot analysis was performed on lysates of HeLa cells treated with indicated siRNA and immunoblotted with indicated antibodies. The signal of mature glycosylated Itgβ1 is marked with an arrowhead, and the lower band indicates the immature non- glycosylated form. In the TECPR2 blot, the (*) asterisk indicates a non-specific band detected by the anti-TECPR2 antibody. B-D) Representative confocal images of HeLa cells treated with the indicated siRNA, followed by immunostaining with anti-Rab5 and anti-SNX17 antibodies. The yellow arrowheads indicate colocalization of SNX17 with Rab5. Scale bars: 10 µm (main); 5 µm (inset). In (D), the graph represents the quantification of SNX17 puncta number in HeLa cells following indicated siRNA treatment. n denotes the total number of cells examined over three independent experiments. Experiments are color-coded, and each dot represents the individual data points from each experiment (****p<0.0001; unpaired two-tailed Student’s t test). E and F) Time- lapse images of HeLa cells treated with control siRNA (E) and TECPR2 siRNA (F) and expressing SNX17-GFP (green) and surface internalized Itgβ1 (magenta) (see Supplementary Video S7). To visualize the surface endocytosed Itgβ1 (magenta), cells were pre-incubated with a complex of primary antibodies against active Itgβ1 (12G10) labeled with Alexa Flour-568 conjugated secondary antibodies. The white arrowheads indicate endosomes positive for both SNX17-GFP and endocytosed Itgβ1. Scale bars: 5 µm.

To understand whether and how TECPR2 regulates SNX17 function, we investigated SNX17 localization on early endosomes in control and TECPR2-depleted cells. Strikingly, TECPR2 knockdown led to a striking reduction in SNX17 vesicular localization (Fig. 6B-D). Importantly, we found that SNX17 recruitment on endocytosed Itgβ1 receptor-containing endosomes was strongly reduced upon TECPR2 depletion (PCC in control siRNA-treated cells 0.44±0.06 versus in TECPR2 siRNA-treated cells 0.17±0.04, n=29 cells; see representative images in Fig. 6E-F and Supplementary Video S7).

This striking reduction in SNX17 vesicular localization upon TECPR2 depletion prompted us to investigate whether TECPR2 interacts with SNX17. Indeed, SNX17 was colocalized with TECPR2 on enlarged early endosomes in cells expressing Rab5 (Q79L), and both TECPR2 and SNX17 were colocalized on active Itgβ1 receptor-containing endosomes (Supplementary Fig. S6A and S7A and Supplementary Video S8). To test whether SNX17 interacts with TECPR2, we performed a GST pulldown assay using GST or GST-SNX17 as the bait. As shown in Fig. 7B and Supplementary Fig. S6B, we found that both overexpressed and endogenous TECPR2 interacts with SNX17. Next, to determine which region of TECPR2 binds to SNX17, we expressed C-terminal and N-terminal domain deletion mutants of TECPR2 with Halo-tagged-SNX17 (Halo-SNX17). We found that SNX17 colocalized with TECPR2_TECPR (935-1411 a.a.) but not with the N-terminal region (1-944 a.a.), indicating that SNX17 likely interacts with the C-terminal TECPR repeat-containing region (Supplementary Fig. S6C-D). To determine whether the C-terminal TECPR repeat-containing region of TECPR2 directly binds to SNX17, we incubated MBP-TECPR2_TECPR with GST or GST-SNX17 in a purified protein- protein interaction assay. Indeed, we found direct binding of SNX17 to the TECPR2 C-terminal TECPR repeat-containing region. Notably, SNX17 showed interaction with the HSAN9- associated Rab5 binding-defective version of TECPR2_TECPR (R1336W), although the binding was reduced compared to the WT protein (Fig. 7C). These observations suggest that TECPR2 has distinct binding interfaces for interactions with SNX17 and Rab5.

**Figure 7:**
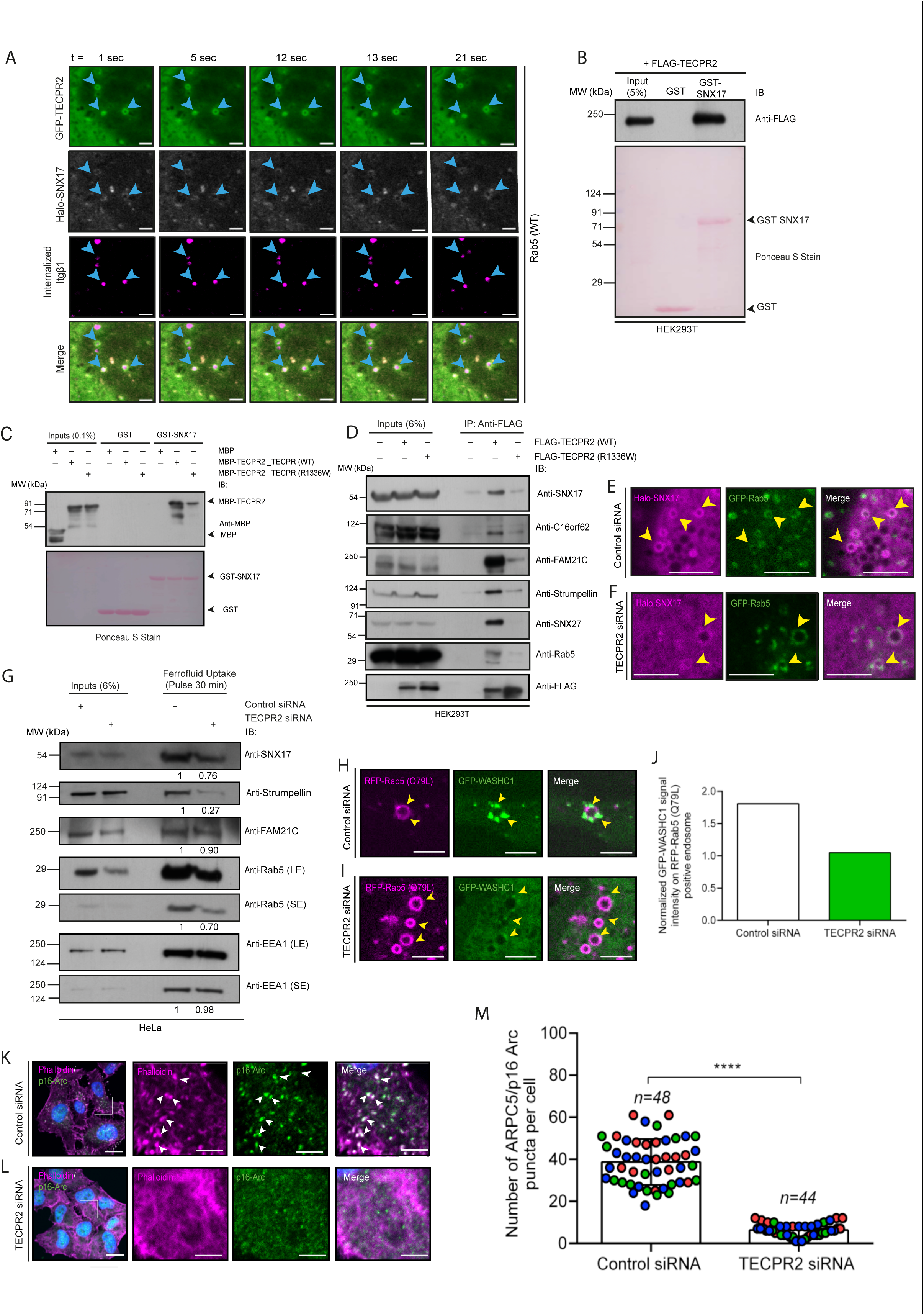
TECPR2 directly interacts with SNX17 and regulates association of SNX17, Retriever, and WASH Complex subunits with Rab5-positive endosomes. A) Time-lapse images of HeLa cells expressing GFP-TECPR2 (green), Halo-SNX17 (grey), untagged-Rab5 (WT), and surface internalized Itgβ1 (magenta) (see Supplementary Video S8). To visualize the surface endocytosed Itgβ1 (magenta), cells were pre-incubated with a complex of primary antibodies against active Itgβ1 (12G10) labeled with Alexa Flour-568 conjugated secondary antibodies. To visualize Halo-SNX17, cell-permeant HaloTag TMR ligand was used. The arrowheads indicate endosomes positive for GFP-TECPR2, Halo-SNX17, and internalized Itgβ1. Scale bars: 5 µm. B) Recombinant GST and GST-SNX17 proteins immobilized on glutathione- conjugated beads were incubated with lysates of HEK293T cells expressing FLAG-TECPR2. The precipitates were IB with an anti-FLAG antibody, and Ponceau S staining was done to visualize the purified proteins. C) Recombinant GST and GST-SNX17 proteins immobilized on glutathione- conjugated beads were incubated with purified MBP or MBP-TECPR2_TECPR (WT or R1336W). The precipitates were IB with an anti-MBP antibody, and Ponceau S staining was done to visualize the purified proteins. D) Lysates of HEK293T cells expressing FLAG-TECPR2 (WT or R1336W) were immunoprecipitated (IP) with anti-FLAG antibody-conjugated beads and IB with the indicated antibodies. E and F) Representative single frame time-lapse images of HeLa cells expressing Halo-SNX17 (magenta) and GFP-Rab5 (green) and treated with control siRNA (E) and TECPR2 siRNA (F). To visualize Halo-SNX17, cell-permeant HaloTag TMR ligand was used. The arrowheads indicate endosomes positive for Halo-SNX17 and GFP-Rab5. Scale bars: 5 µm. G) The ferrofluid particle uptake for 30 min (pulse) at 37°C was performed in HeLa cells treated with control and TECPR2 siRNA. The cells were homogenized, and the ferrofluid- containing compartments were purified and IB for the presence of indicated proteins. The values represent the densitometric analysis of the indicated protein levels in the purified ferrofluid- containing endosomes, normalized to their input levels. H-J) Representative single frame time- lapse images of HeLa cells expressing RFP-Rab5 (Q79L) (magenta) and GFP-WASHC1 (green) and treated with control siRNA (H) and TECPR2 siRNA (I) (see Supplementary Video S9). The arrowheads indicate endosomes positive for RFP-Rab5 (Q79L) and GFP-WASHC1. Scale bars: 5 µm. The graph in (J) represents the quantification of normalized GFP-WASHC1 signal intensity on RFP-Rab5 (Q79L)-positive endosomes. K-M) Representative confocal images of HeLa cells treated with control siRNA (K) and TECPR2 siRNA (L), followed by immunostaining with anti- p16-Arc antibodies (green) and actin labeling using Alexa Fluor-647 conjugated phalloidin (magenta). The arrowheads denote localization on p16-Arc on actin patches. Scale bars: 10 µm (main); 5 µm (inset). In (M), the quantification of the number of the p16-Arc puncta in HeLa cells upon indicated siRNA treatment is shown. Experiments are color-coded, and each dot represents the individual data points from each experiment (****p<0.0001; unpaired two-tailed Student’s t test).

We envisioned that in intact cells, as the HSAN9-associated Rab5 binding-defective form of TECPR2 is cytosolic, it may not interact with SNX17. Indeed, in total cell lysates, we found a greatly reduced interaction between the TECPR2 (R1336W) variant and SNX17 compared to the WT TECPR2 (Fig. 7D). Previous studies have shown that SNX17 interacts with the Retriever- CCC-WASH complexes, which generate branched actin-enriched subdomains on early endosomes, leading to cargo sequestration and, together with the formation of transport carriers, cargo recycling (Cullen and Steinberg, 2018; McNally et al., 2017; Simonetti and Cullen, 2019). We found that the C16orf62 subunit of the Retriever complex and WASH complex subunits

FAM21C and Strumpellin also immunoprecipitated with TECPR. Furthermore, TECPR2 (R1336W) showed reduced interaction with the subunits of the Retriever and WASH complexes (Fig. 7D). TECPR2 also co-immunoprecipitated SNX27, confirming the previously reported high- throughput interaction data (Fig. 7D) (Nalbach et al., 2023; Stadel et al., 2015). We also found that TECPR2 colocalized with FAM21C and Strumpellin on enlarged early endosomes in cells expressing Rab5 (Q79L) (Supplementary Fig S6E-F). Furthermore, TECPR2 colocalized with the ARP2/3 complex (as observed by immunostaining for the p16-Arc subunit of ARP2/3) on enlarged early endosomes in cells expressing Rab5 (Q79L) (Supplementary Fig. S6G).

We next investigated the hypothesis that TECPR2 acts as a linker between Rab5 and SNX17 and regulates the recruitment of integrin retrieval machinery on early endosomes. Consistent with this, we found that localization of SNX17 on Rab5-positive early endosomes was strongly reduced upon TECPR2 knockdown (PCC in control siRNA-treated cells 0.48±0.09 versus in TECPR siRNA-treated cells 0.18±0.03, n=30 cells, Fig. 7E-F). Next, we purified FF-loaded endosomes from control and TECPR2-depleted homogenates after 30 min of pulse, when the fractions are enriched for the early endosomal marker protein EEA1 (Fig. 7G). Indeed, we observed decreased levels of SNX17 and WASH subunit Strumpellin, but not FAM21C, in early endosomal fractions upon TECPR2 depletion, suggesting that TECPR2 regulates early endosomal recruitment of specific proteins of the integrin retrieval machinery (Fig. 7G). In accordance with this, we found that TECPR2 depletion resulted in strikingly reduced localization of WASH subunit WASHC1 on Rab5-positive early endosomes. (Fig 7H-J and Supplementary Video S9). Consequently, recruitment of the ARP2/3 complex and the presence of actin on early endosomes was strikingly reduced upon TECPR2 depletion (Fig 7K-M).

TECPR2 knockdown in zebrafish induces defects in hatching and motility To elucidate the role of TECPR2 in vivo, we used zebrafish as a model organism to study the effects of its depletion at the organismal level. The zebrafish genome encodes two transcripts for tecpr2 (tecpr2-202 encoding a protein of length 1358 a.a. and tecpr2-201 encoding a protein of length 1308 a.a.), which are 96.3% identical. The sequence identity and similarity between the human and zebrafish Tecpr2 longer isoform is 53.6% and 64.3%, respectively, and with Tecpr2 shorter isoform is 55.4% and 66.3%, respectively. Importantly, we found that Tecpr2 (shorter isoform) interacted with the Rab5 ortholog in zebrafish, suggesting that this interaction is conserved across vertebrate evolution (Fig. 8A). We observed the expression of tecpr2 transcripts in zebrafish embryos at the one-cell stage, which sharply decreased 24 hours post-fertilization (hpf) and gradually increased between 72 and 96 hpf (Fig. 8B). The tecpr2 expression showed a peak at 4 days post-fertilization (dpf), a time when the larvae are freely swimming and feeding. Whole-mount in situ hybridization analysis revealed that tecpr2 was expressed in 0 dpf embryos; a faint expression in the brain and retina was observed at 1 dpf; at the forebrain-midbrain boundaries at 2 dpf; in the brain, retina, and anterior myotomes at 3 dpf; in the brain, swim bladder, and heart at 4 dpf; and tecpr2 expression was observed in the brain, myotomes, and retina at 5 dpf embryos (Fig. 8C).

**Figure 8:**
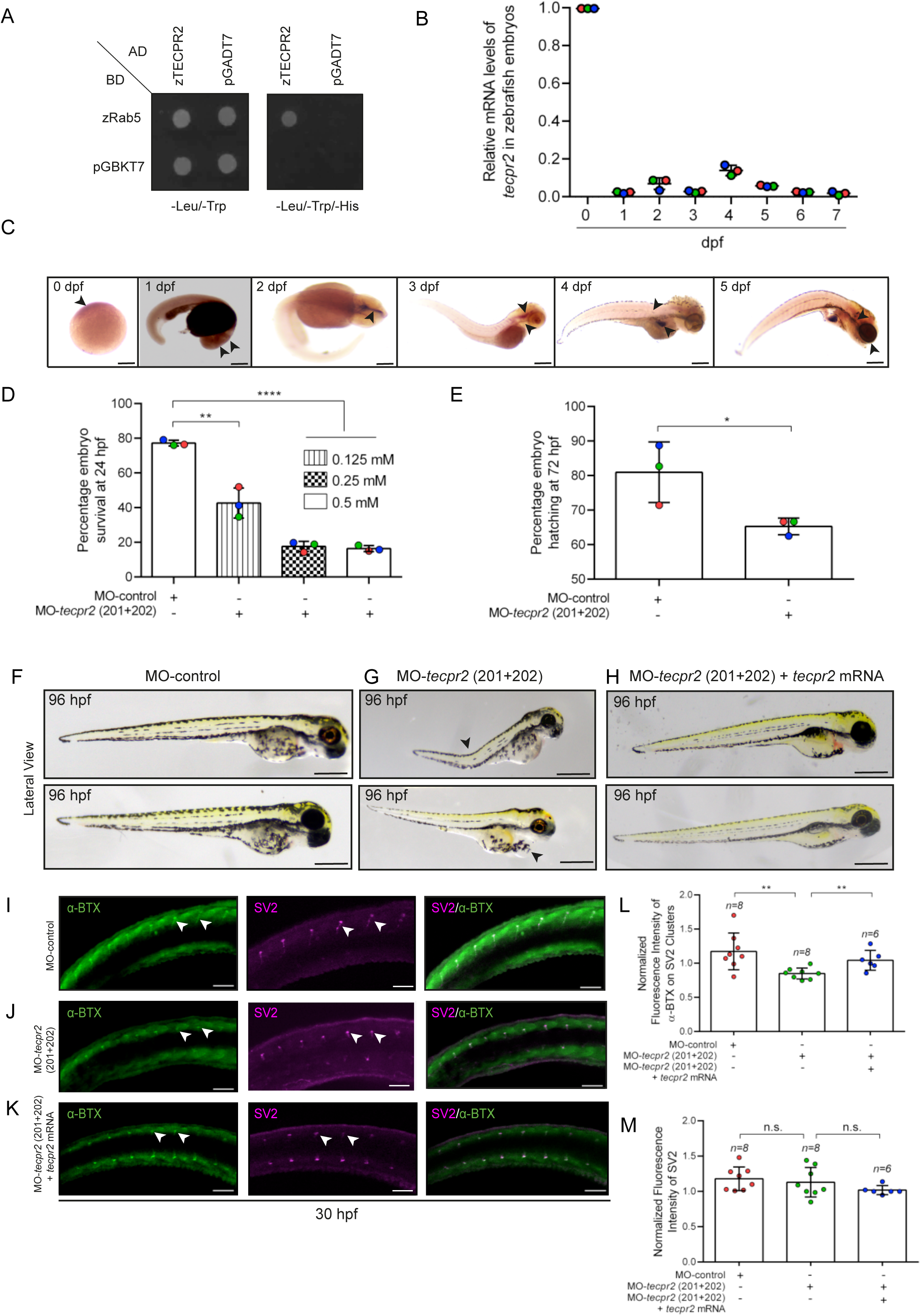
Tecpr2 depletion in zebrafish results in reduced survival, hatching defect, and impaired motility. A) Yeast two-hybrid interaction analysis of zebrafish TECPR2 with zebrafish Rab5. Co-transformants were spotted on -Leu/-Trp to confirm viability and -Leu/-Trp/-His media to detect interactions. B) Zebrafish embryos were collected at different days post fertilization (dpf), and the relative mRNA levels of tecpr2 were determined using qRT-PCR. C) Representative brightfield images of zebrafish embryos showing spatial expression of tecpr2 transcripts at different developmental stages. Scale bars: 200 µm. D and E) The graph represents dose-dependent survival of zebrafish embryos injected with MO-control and MO-tecpr2 (201+202) morpholinos at 24 hpf (D) and embryos hatching at 72 hpf (E) (****p<0.0001; **p<0.01; *p=0.040; unpaired two-tailed Student’s t test). F-H) Lateral view of zebrafish embryos injected with MO-control (F), MO-tecpr2 (201+202) (G) and MO-tecpr2 (201+202) + tecpr2 mRNA (rescue) (H) at 96 hpf. The tecpr2 morphants (G) showed tail curvature and developed pericardial oedema (marked by arrowheads), and this phenotype was rescued upon introduction of tecpr2 mRNA (H) into morphants. Scale bars: 200 µm. I-K) Representative confocal images of whole-mount immunostaining of MO-control (I), tecpr2 morphants (J), and morphants injected with tecpr2 mRNA (K) were fixed at 30 hpf and stained using α-BTX (green) to label the acetylcholine receptors and anti-SV2 antibodies (magenta) to stain the motor neurons. The arrowheads denote spots co-labeled with anti-SV2 antibodies and α-BTX. In tecpr2 morphants (J), intensity of α- BTX was reduced on SV2 clusters. Scale bars: 10 µm. L and M) The graph represents fluorescence intensity of α-BTX-positive spots on SV2 clusters (L) and fluorescence intensity of SV2 (M) in zebrafish embryos treated with MO-control, MO-tecpr2 (201+202), MO-tecpr2 (201+202) + tecpr2 mRNA (rescue) (**p<0.01; n.s., non-significant; unpaired two-tailed Student’s t test).

Next, we depleted both zebrafish Tecpr2 isoforms by targeted gene knockdown using morpholino (MO)-based antisense oligonucleotides against tecpr2 201 and 202 transcripts in single-cell stage embryos. The knockdown efficiency of the morpholinos was determined using GFP reporter constructs containing MO-target sequences that were co-injected into embryos along with their respective MO. Embryos injected with the control morpholino was used as controls. As shown in Supplementary Fig. S7A, control embryos showed GFP expression in 100% of the embryos, whereas almost no expression of the GFP reporter was observed in embryos co-injected with either MO-tecpr2 201 or MO-tecpr2 202, confirming the efficiency of MO in knocking down gene expression.

We next assessed embryo survival and any morphological changes in zebrafish one-cell- stage embryos injected with control or combined MO-tecpr2 (201 and 202). Compared to the control MO-injected embryos, the majority of the tecpr2 MO-injected embryos, the morphants, did not survive at 24 hpf (Fig. 8D). Furthermore, the surviving Tecpr2-morphants displayed a severe hatching defect at 72 hpf (Fig. 8E). Tecpr2-morphants displayed tail curvature, decreased body length, and pericardial oedema (Fig. 8F-G and Supplementary Fig. S7B-D). The tecpr2- 201 mRNA, when co-injected with MO-tecpr2 (201+202), rescued the viability and morphological defects of Tecpr2 depletion (Fig. 8H and Fig. S7E-F). Whole-mount immunostaining of the HA epitope confirmed the translation of the injected tecpr2 mRNA (Fig. S7G).

At 4 dpf, we observed impaired motility in tecpr2 (201+202) morphants compared to that in control MO-treated embryos in response to touch stimulation. Co-injection of tecpr2-201 mRNA restored normal motility in tecpr2 (201+202) morphants, indicating that motility defects are due to the reduced expression of tecpr2 transcripts (Supplementary Video S10). We speculated that the absence of Tecpr2 affected neuromuscular junction (NMJ) architecture, leading to defective locomotion in Tecpr2 morphants. To investigate this, we assessed changes in developing NMJ in control and Tecpr2-depleted embryos. We performed whole-mount immunostaining of control and MO-tecpr2 (201+202) morphants at an early time point of development (30 hpf), employing antibodies against SV2 (presynaptic vesicles) and α- bungarotoxin (α-BTX), which binds to postsynaptic acetylcholine receptors (AChRs). At 30 hpf, we observed SV2 as spots or dots colocalizing with α-BTX in the control morphants (Fig. 8I-J). Notably, we found reduced intensity of postsynaptic AChRs at the SV2 spots/dots in Tecpr2- depleted morphants compared to the control morphants, and this effect was rescued upon re- expression of tecpr2-201 mRNA. No significant change in SV2 intensity was observed in Tecpr2- depleted morphants (Fig. 8K-M). Our results suggest that Tecpr2 may regulate postsynaptic acetylcholine receptor levels at the site of NMJ formation, which is essential for normal motility of zebrafish larvae.

## Discussion

Previous studies on TECPR2 have not addressed how its subcellular localization is regulated. To address this question, we performed small-scale screening of Rab G proteins that might interact with TECPR2. The motivation behind such an approach was that the Rabs, Arfs, and Arl families of small G proteins generally act as recruiting agents to bring their partners to specific intracellular compartments. We found that TECPR2 interacts with Rab5, a master regulator of cargo traffic, to and from early endosomes. Based on the findings from this study, we conclude that TECPR2 is a Rab5 effector, implying that Rab5 directly interacts with and recruits TECPR2 on early endosomes. We also found that the HSAN9-associated R1336W TECPR2 variant showed reduced binding to Rab5, and accordingly, this variant did not show membrane localization.

TECPR2-positive early endosomes were dynamic, accessible to endocytic cargo, and undergo frequent budding and fusion events. Occasionally, we also found that TECPR2 was localized to relatively long and stable tubular membranes. We noted a dramatic collapse of early and recycling endosomes in the perinuclear region upon TECPR2 depletion. This was most prominently observed upon analysis of recycling endosomal markers, including TfR and Rab11, which showed dramatic redistribution from the periphery to the perinuclear region in TECPR2- depleted cells. TECPR2 depletion also led to an increased overlap between SNX1, a marker for the retrograde sorting subdomain, and HRS, a marker for the degradative subdomain on early endosomes, coupled with an increase in the size of LAMP1 positive late endosomal and lysosomal compartments. These findings suggest that upon TECPR2 depletion, there might be a defect in the segregation of the sorting/recycling and degradative subdomains of early endosomes.

The most striking phenotype of TECPR2 depletion in multiple cell types was the reduced cell spreading area. This prompted us to analyze whether surface integrin levels are affected by TECPR2 depletion. Indeed, we found that integrin recycling was impaired and degradation was increased upon TECPR2 depletion, leading to reduced levels of integrins at the cell surface. To understand the mechanism by which TECPR2 regulates cargo recycling, we analyzed the localization of molecular players known to mediate integrin retrieval from the lysosomal degradation pathways. SNX17 is a key player that recognizes the sorting motif in the integrin cytoplasmic tail and sorts integrins into the recycling subdomain of early endosomes. SNX17 interacts with the retriever complex that further interacts with the CCC and WASH complexes that mediate recruitment of the Arp2/3 complex, which promotes branched actin polymerization to promote the formation of tubular recycling domains (Cullen and Steinberg, 2018; McNally et al., 2017; Simonetti and Cullen, 2019; Steinberg et al., 2012).

Two observations prompted us to analyze whether TECPR2 and SNX17 interact and play a role in the same pathway. First, the defect in integrin retrieval from lysosomal degradation upon TECPR2 depletion was similar to the SNX17 depletion phenotype. No additive effect was observed upon co-depletion of TECPR2 and SNX17, suggesting that they act in the same pathway. Moreover, SNX17 belongs to the SNX-FERM subfamily similar to SNX27, which has been reported as a potential interaction partner of TECPR2 in high-throughput proteomics screening experiments (Nalbach et al., 2023; Stadel et al., 2015). Indeed, we found that TECPR2 directly interacts with SNX17 via its C-terminal region containing TECPR repeats, and SNX17 early endosomal recruitment was reduced upon TECPR2 depletion (Fig. 9). It will be insightful in future studies to determine the binding interface residues in TECPR2 that mediate binding to SNX17 and whether HSAN9-associated variants affect binding to SNX17. While in this study we have only analyzed integrin recycling, TECPR2-depleted cells likely have defects in recycling of other cargo receptors, such as low-density lipoprotein receptor-related protein 1 (LRP1), which are regulated by SNX17 (Burden et al., 2004; Farfan et al., 2013; McNally et al., 2017).

**Figure 9:**
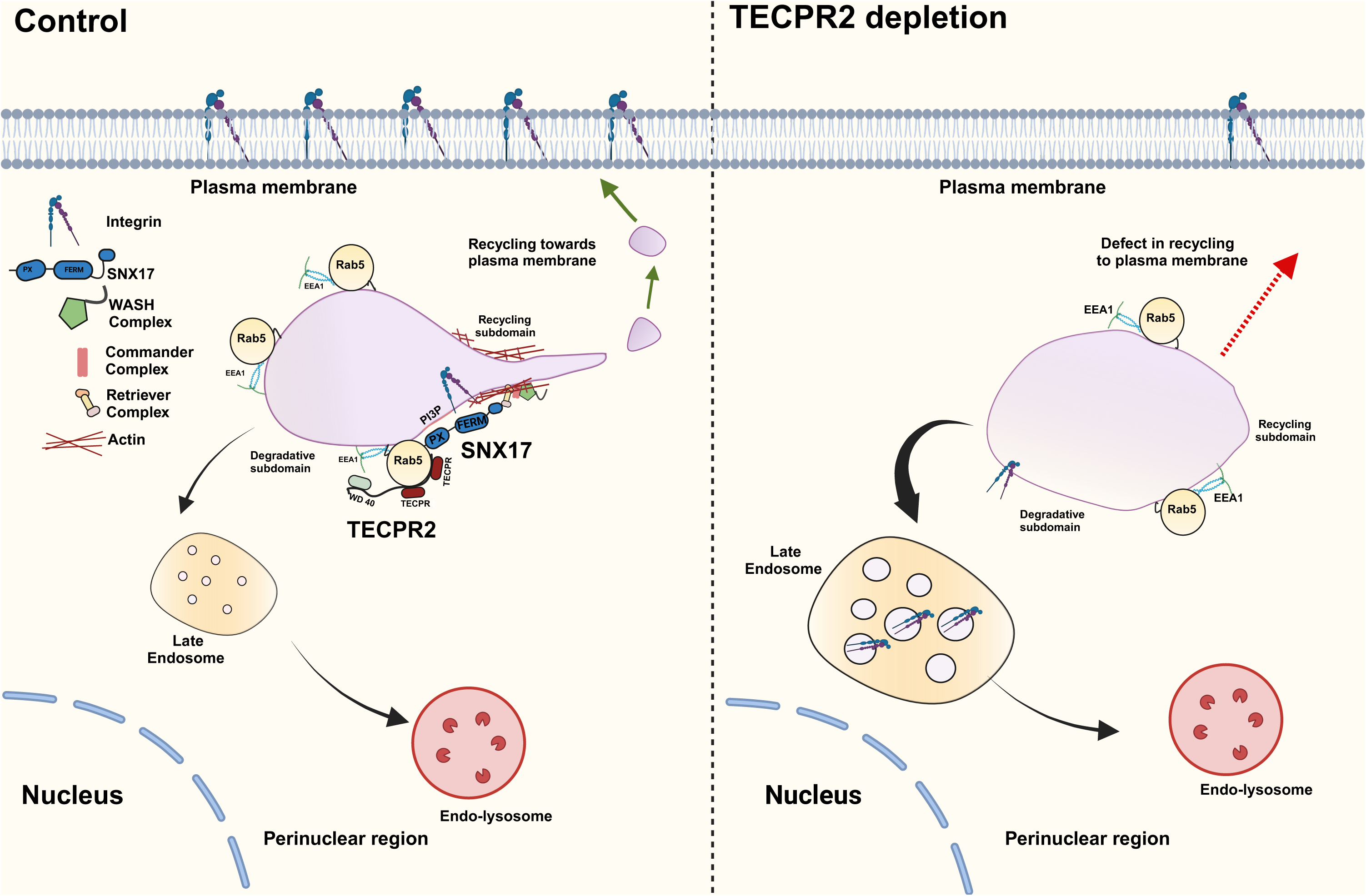
Proposed role of TECPR2 in regulating cargo recycling to the plasma membrane. TECPR2 is a Rab5 effector, implying that Rab5 directly interacts with and recruits TECPR2 on early endosomes. TECPR2 interacts with cargo (integrin receptors) retrieval machinery on early endosomes, including the sorting adaptor, SNX17, and associated protein complexes of Retriever and WASH, and mediates cargo recycling to the plasma membrane. TECPR2 depletion leads to reduced association of SNX17 and WASH complex subunits with early endosomes and impaired retrieval of integrin receptors from the lysosomal degradation pathway.

We also found reduced levels of the WASH subunits WASH and Strumpellin on Rab5- positive early endosomes in TECPR2 depleted cells. Consequently, early endosomal localization of the Arp2/3 subunit p16-Arc was reduced in TECPR2-depleted cells (Fig. 9). It remains to be tested whether TECPR2 directly interacts with any subunit of the WASH complex on early endosomes. Indeed, WASH-knockout cells also exhibit collapse of early endosomal markers in the perinuclear region, similar to TECPR2 depletion (Gomez et al., 2012). Furthermore, zebrafish embryos depleted of the WASH complex subunit, Strumpellin, show the same phenotypes as Tecpr2-depleted embryos, such as impaired motility and tail curvature (Clemen et al., 2010).

Our study does not address how TECPR2’s role as a Rab5 effector aligns with its previously described roles in maintaining ERES and regulating cargo export from the secretory pathway. We did not observe TECPR2 localization to either the ER or the Golgi membranes upon live-cell imaging in HeLa cells. Instead, epitope-tagged-TECPR2 was either cytosolic or present on vesicles that were in contact with the ER tubules, a behavior reminiscent of early endosomes in contact with the ER network. Notably, we observed that TECPR2-positive endosomes colocalized with VAPB, an ER membrane contact site protein that has been previously shown to interact with TECPR2 (Nalbach et al., 2023). The phenotypes observed in TECPR2-depleted cells, i.e., enlarged early endosomes and lysosomes, reduced localization of actin polymerization machinery on early endosomes, and impaired cargo recycling, might result from reduced ER-early endosome contact and ER-mediated endosomal tubule fission (Gomez et al., 2012; Rowland et al., 2014).

We hypothesize that impaired cargo recycling due to defects in ER-endosome contacts and ER- and actin-mediated endosomal tubule fission underlie the neurodevelopmental delay and neurodegeneration pathologies in patients with loss-of-TECPR2 mutations. In strong support of this hypothesis, mutations in ER-shaping (Spastin and REEP1) and endosomal fission proteins (WASH subunit-Strumpellin and Spastin) are associated with the most common forms of HSPs (Allison et al., 2017; Blackstone et al., 2011; Park et al., 2010). It is not known whether the contact sites between the ER and early endosomes also regulate ER shape and ERES formation; the latter has been shown to be reduced in TECPR2-depleted cells (Stadel et al., 2015).

While we analyzed TECPR2 function in non-neuronal cells, we predict that TECPR2 plays a similar role in regulating receptor retrieval and recycling in neurons. We propose that loss of TECPR2 function in neurons would likely affect the recycling of cargo recognized by both sorting adaptors SNX17 and SNX27, as the latter protein also interacts with TECPR2. Cargoes include receptors such as β1-integrin (Rivero-Rios et al., 2023), β2-adrenergic receptors (Lauffer et al., 2010), AMPA receptors (Hussain et al., 2014; Loo et al., 2014; Temkin et al., 2017), and neuroligin-2 (Halff et al., 2019), which are essential for neurite outgrowth, axon guidance, neuronal signalling, and function. Future studies on TECPR2 function and understanding HSAN9 disorder should investigate the endocytic recycling of SNX17 and SNX27-dependent cargo receptors in healthy and HSAN9 afflicted patient neurons. Indeed, as described in this study, zebrafish will be a useful in vivo model system for understanding and identifying the major cargo receptors dependent on TECPR2 for recycling. The locomotor defects observed in zebrafish embryos lacking Tecpr2 are reminiscent of the progressive spasticity observed in human patients.

Previous studies have shown that TECPR2 regulates autophagosome biogenesis and autophagosome-lysosome fusion (Fraiberg et al., 2021; Stadel et al., 2015; Tamim-Yecheskel et al., 2021). TECPR2 engagement with LC3 and Rab5 is likely via separate binding sites with LC3 binding mediated by “WEVI” motif in TECPR2 (1407-1411 a.a.) whereas the exact Rab5 binding site remains to be experimentally determined. A potential hypothesis for future research is that TECPR2 facilitates the maturation of autophagosomes to amphisomes by mediating the interaction and fusion of autophagosomes and Rab5-positive endosomes.

In conclusion, our study reveals that TECPR2 localizes to early endosomes in a Rab5- dependent manner, wherein it mediates the recruitment of molecular complexes that sequester specific cargo receptors and generate actin-rich retrieval domains on early endosomes required for the recycling of these cargo receptors back to the plasma membrane.

### Material and Methods

### Cell culture

HeLa and HEK293T (from ATCC) cells were cultured in DMEM media (Gibco) supplemented with 10% heat-inactivated fetal bovine serum (FBS; Gibco) in a humidified cell culture incubator (New Brunswick) with 5% CO2 at 37°C. hTERT-RPE1 cells (from ATCC) were cultured in DMEM/F-12 media (Gibco), supplemented with 10% FBS. For the live-cell imaging and flow cytometry experiments, cells were cultured in phenol red-free media (Gibco) supplemented with 10% FBS. All the cell lines were subcultured for not more than 18 passages and were routinely screened for the absence of mycoplasma contamination using the MycoAlert Mycoplasma Detection Kit (Lonza).

### DNA constructs, antibodies and chemicals

Supplementary Tables II and III list all the DNA constructs and antibodies used in this study, respectively. Most of the chemicals used in this study were purchased from Sigma-Aldrich. Alexa fluor-conjugated transferrin (Tfn), -phalloidin, -α-Bungarotoxin (BTX), and DAPI were purchased from Molecular Probes. All the lipids used in the preparation of Giant unilamellar vesicles (GUVs) were purchased from Avanti Research. Water-based ferrofluid (EMG 508) was purchased from Ferrotec Corporation. The cell-permeant HaloTag TMR ligand (G8252) and Janelia Fluor 646 HaloTag ligand (GA1120) were purchased from Promega. The avidin egg white (A2667) and TRIzol Reagent (15596026) were purchased from Invitrogen.

### Gene silencing

The siRNA oligos for gene silencing were purchased from Dharmacon (Horizon Discovery) and prepared according to the manufacturer’s instructions. Transient transfection of siRNAs was performed using the DharmaFECT 1 reagent according to the manufacturer’s instructions. The following siRNAs were used in this study: control siRNA, 5’-TGGTTTACATGTCGACTAA-3’; TECPR2 siRNA, 5’- CCAGATCAGTTTGCAGAAA-3’; TECPR2 siRNA oligo #2, 5’- GGAGATAGGAACATTATAA-3’ and TECPR2 siRNA oligo #3, 5’- GGAGTTTACTTCCGTGTAG-3’; and SNX17 siRNA, 5’-CCAGTGATGTCCACGGCAATT-3’. The following pool of siRNA oligos targeting all the three Rab5 paralogs, Rab5a, Rab5b and Rab5c, were used: 5’-AGGAATCAGTGTTGTAGTA-3’, 5’-GAAAGTCAAGCCTGGTATT-3’, and 5’- CAATGAACGTGAACGAAAT-3’.

For gene silencing in zebrafish embryos, the following morpholinos (MO), purchased from Gene Tools, LLC, were used: MO control, 5’-CCTCTTACCTCAGTTACAATTTATATA-3’; MO tecpr2-201, 5’-GATCAATAAAAGAGGTTGCTGTGTT-3’; and MO tecpr2-202, 5’- CGTTTACATTGATTCTGAAGCCTAC-3’.

Transfection, immunofluorescence and live-cell imaging Cells grown on glass coverslips (VWR) were transfected with desired plasmids using X- tremeGENE HP DNA transfection reagent (Roche) for 16 –18 hr. After transfection, cells were fixed with 4% paraformaldehyde (PFA) in PHEM buffer (60 mM PIPES, 10 mM EGTA, 25 mM HEPES, 2 mM MgCl2, pH 6.8) for 10 min at room temperature (RT). The cells were then blocked in blocking solution (0.2% saponin + 5% NGS in PHEM buffer) for 30 min at RT and washed three times with 1X PBS. Following blocking, cells were incubated with primary antibodies in staining solution (PHEM buffer + 0.2% saponin + 1% NGS) for 2 hr at RT or overnight at 4°C, washed three times using 1X PBS, and incubated with Alexa Fluor-conjugated secondary antibodies in PHEM + 0.2% saponin for 30 min at RT. Finally, coverslips were washed three times with 1X PBS and mounted on glass slides in Fluoromount-G (Southern Biotech). Single-plane confocal images were acquired on LSM710 confocal microscope using a 63×/1.4 NA oil immersion objective and Zen Black 2012 software (ZEISS). Super-resolution images were acquired on LSM 980 Elyra 7 super-resolution microscope equipped with a plan apochromat 63×/1.4 NA oil immersion objective and ZEN 2012 v. 8.0.1.273 (ZEISS) was used to acquire images. All the representative confocal images were adjusted for brightness and contrast using ImageJ (NIH). For phase contrast imaging, cells grown on glass coverslips were fixed with 4% PFA and imaged using the Nikon Eclipse TS100 inverted microscope using a 10× objective lens.

For live-cell imaging experiments, cells were grown on glass-bottom imaging dishes (Ibidi), followed by transfection of indicated plasmids. After transfection for 12 hr, complete DMEM media was replaced by phenol red-free complete DMEM (Gibco) media, and the dish was placed in a live-cell imaging chamber that was maintained at 37°C with a 5% CO2 supply. Time-lapse imaging was performed on an Olympus IXplore SpinSR Spinning Disk Super Resolution Confocal Microscope using a 100× Apo/1.45 NA oil objective. The videos were acquired using the cellSens dimension software (Olympus), and the final adjustments for brightness and contrast were done using the Fiji/ImageJ.

### Image analysis and quantification

#### Colocalization analysis

To analyze colocalization, images were opened in Fiji software. For quantifying the Pearson’s correlation coefficient (PCC) between GFP-TECPR2 (WT) with endogenous markers or FLAG- TECPR2 mutants with GFP-Rab5, manual thresholding was performed. For this, a background subtraction was first done by selecting a small area near the nucleus, the background intensity was subtracted by using the subtract option under the “Math tool” in the “Process” menu of Fiji. For quantification of PCC of other immunofluorescence images that were showing punctate staining, the “Max Entropy” thresholding was applied. Finally, after thresholding, JACoP plugin was used to determine the PCC.

#### Quantification of size and number of puncta

For calculating the size of EEA1 puncta, the images were opened in Fiji software and a threshold was applied by using the “Max Entropy” thresholding option. The number of EEA1 endosomes was then calculated by “Analyze Particles” option in Fiji, under the “Analyze” menu. To calculate the number of puncta of SNX17 and p16^Arc^, manual thresholding was applied such that only the vesicular staining was visible. The number of puncta was then calculated by “Analyze Particles” option in Fiji.

#### Fractional distance of endosomes

The distribution of early/recycling endosomal markers was performed by measuring the fractional distance of these endosomes. A line was drawn from the centre of the cell to the cell periphery using the “line tool” of Fiji software in the confocal micrograph. The plot profile tool was then used which selects all the endosomal marker fluorescent intensities along with their corresponding distance values on the drawn line. The background pixels and their corresponding distance values were not considered for analysis. The remaining distances corresponding to the endosomal pixels were then converted to fractional distance. This was done by dividing all these values by the total distance of the line drawn.

#### Quantification of fission and fusion events

For calculation of fission and fusion events, GFP-TECPR2-positive vesicles near the peripheral region were chosen. A straight line was drawn along the vesicles undergoing fission or fusion and the mean gray value was measured at each frame during the fission or fusion events. The line drawn was moved in each frame according to the movement of the vesicles in order to compensate small movements of vesicles during live cell imaging. The mean gray values of each vesicle at each frame were recorded and these values were normalized to its respective highest value. These normalized data for each vesicle was analyzed by determining their linear regression lines and the slopes were compared to determine the fission and fusion events of the vesicles. The positive and negative slopes in the graph (Fig. 3C) were marked in green and red lines, which indicate overall fission and fusion of a single vesicle.

### Yeast two-hybrid assay

To perform yeast two-hybrid assay, previously described methodology was used (Sharma et al., 2021). Briefly, the bait gene was cloned in the pGBKT7 vector in fusion with Gal4-DNA-binding domain, and the prey gene was cloned in the pGADT7 vector in fusion with Gal4-activation domain. The plasmids were co-transformed in the transformation-competent Y2H Gold strain of S. cerevisiae (Takara Bio Inc.). The transformants were plated on double-dropout plates that were lacking tryptophan and leucine (-Leu/-Trp) and were allowed to grow at 30°C for 3 days to select yeast cells containing both the bait and prey plasmids. The co-transformant colonies of yeasts were plated on non-selective (-Leu/-Trp) and selective media (-Leu/-Trp/-His) or (-Leu/-Trp/-His/-Ade) to assess viability and interaction between the bait and prey proteins, respectively.

### Cell lysates, co-immunoprecipitation, and immunoblotting

For lysate preparation, cells were collected from culture dishes by trypsinization and centrifugation at 400×g for 2 min. The pellet obtained was washed two times with DPBS buffer (Lonza) and lysed in ice-cold RIPA lysis buffer (10 mM Tris-Cl pH 8.0, 1 mM EDTA, 0.5 mM EGTA, 1% Triton X-100, 0.1% SDS, 0.1% sodium deoxycholate, 140 mM NaCl supplemented with 1 mM PMSF and a protease inhibitor cocktail (Sigma-Aldrich). The samples were incubated on ice for 2 min, followed by a 30-sec vortex, and this cycle was repeated five times prior to centrifugation at 16,000×g for 5 min at 4°C. The resultant clear supernatants were transferred to a new microcentrifuge tube, and using a bicinchoninic acid assay kit (Sigma-Aldrich), the amount of protein was quantified. Protein samples were prepared for immunoblotting analysis by boiling them in a 4X sample loading buffer and then loaded onto SDS-PAGE.

To perform the co-immunoprecipitation (co-IP) assay, the previously described methodology was used (Sharma et al., 2021). Briefly, HEK293T cells were transfected with the desired plasmids expressing FLAG-tag proteins. Post 20 hr transfection, cells were lysed in ice-cold TAP lysis buffer (20 mM Tris-Cl pH 8.0, 150 mM NaCl, 0.5% NP-40, 1 mM MgCl2, 1 mM Na3VO4, 1 mM NaF) supplemented with 1 mM PMSF and protease inhibitor cocktail, and kept on rotation (HulaMixer, Thermo Scientific) for 30 min at 4°C. The cell lysate was centrifuged at 16,000×g for 5 min at 4°C, and the post-nuclear supernatant (PNS) was collected. PNS was incubated for 3 hr at 4°C with an anti-FLAG antibody-conjugated resin (Biolegend), followed by four washes with TAP wash buffer (20 mM Tris-Cl pH 8.0, 150 mM NaCl, 0.1% NP-40, 1 mM MgCl2, 1 mM Na3VO4, 1 mM NaF, and 1 mM PMSF). Protein complexes were eluted by boiling the beads in a 4X sample loading buffer at 100°C for 10 min. The samples were then subjected to SDS-PAGE for western blotting.

For the co-IP experiments involving the use of an anti-GFP nanobody, a recombinant GST-tagged anti-GFP nanobody was first expressed and purified from the E. coli Rosetta (DE3) strain using the protein purification methodology described below, and ∼5 µg of the nanobody-conjugated to the glutathione beads was added to the PNS, and the rest the same protocol was followed.

For the endogenous co-IP of TECPR2 with Rab5, anti-Rab5 primary antibody (2 µg; BD Biosciences) was first incubated with the PNS for overnight at 4°C with rotation, followed by addition of Protein A/G beads to the PNS for 3 hr at 4°C to allow the binding of primary antibody to the beads. The beads were gently washed three times with 1X PBS containing 0.1% NP-40 (Sigma-Aldrich) to remove non-specific protein binding, and samples were then prepared for western blotting analysis by boiling them in 4X sample loading buffer.

For immunoblotting, protein samples separated by SDS-PAGE were transferred onto a 0.2 µm PVDF blotting membrane (Amersham), followed by overnight incubation at 4°C in blocking buffer (10% skim milk (BD Difco) prepared in 1X PBS containing 0.05% Tween 20 (0.05% PBST)). After washing with 0.05% PBST, the blot was incubated with a primary antibody solution prepared in 0.05% PBST for 2 hr at RT or overnight at 4°C. The membranes were washed three times for 10 min each with 0.05% PBST and further incubated with an HRP-conjugated secondary antibody solution prepared in 0.05% PBST for 1 hr at RT. Following the secondary antibody step, the membranes were washed twice for 10 min with 0.3% PBST. The blots were developed by a chemiluminescence-based method (ECL Prime Western Blotting Detection Reagent; Amersham) using medical X-ray films (Retina). For certain blots, membranes were stripped for 2 min at RT and washed three times for 5 min with 0.05% PBST before being blocked and probed with a new antibody. ImageJ software was used to perform densitometry analysis of the immunoblots.

Recombinant protein purification, purified protein-protein interaction assay, and pulldown assay using recombinant proteins In this study, all the His-, GST- or MBP-tagged recombinant proteins were expressed and purified from E. coli Rosetta (DE3) strain. For the setting up of primary cultures, a single transformed colony was inoculated into Luria–Bertani (LB) broth containing the appropriate antibiotics and incubated at 37°C in a shaking incubator. After 12 hr of incubation, 1% of primary cultures were used as inoculum to establish secondary cultures, which were then incubated at 37°C with shaking until the absorbance at 600 nm reached 0.4-0.6. To induce protein expression, 0.3 mM IPTG (Sigma-Aldrich) was added to the cultures, followed by 16 hr of shaking incubation at 16°C. Bacterial cultures were centrifuged at 3220×g for 10 min, washed once with 1X PBS, and resuspended in bacterial cell lysis buffer (20 mM Tris-Cl pH 8.0, 150 mM NaCl) in case of His- and GST-tagged proteins, and with buffer (20 mM Tris-Cl pH 8.0, 150 mM NaCl, 1 mM EDTA, 1 mM DTT) for MBP-tagged proteins. Both the lysis buffers were supplemented with 1 mM PMSF and a protease inhibitor tablet (Roche). The bacterial cells were lysed by sonication followed by 30 min of centrifugation at 10,062×g at 4°C. The clear supernatants were incubated with either His-cobalt resin (Thermo Scientific) for His-tagged proteins or with amylose resin (New England Biolabs) for MBP-tagged proteins, or with glutathione resin (Gbiosciences) for GST-tagged proteins on a HulaMixer for 2 hr at 4°C. To remove impurities, the resins were washed at least five to seven times with a wash buffer (20 mM Tris-Cl pH 8.0, 300 mM NaCl, 10 mM imidazole-for the His-tagged proteins; 20 mM Tris-Cl pH 8.0, 300 mM NaCl- for the GST-taaged proteins; and 20 mM Tris-Cl pH 8.0, 200 mM NaCl, 1 mM EDTA, 1 mM DTT-for the MBP-tagged proteins).

For performing purified protein-protein interaction assay, MBP-tagged proteins were eluted from amylose resin using elution buffer (20 mM Tris-Cl pH 8.0, 200 mM NaCl, 1mM EDTA, 1 mM DTT, 20 mM maltose monohydrate) and His-tagged proteins were eluted from the His-cobalt resin using elution buffer (50 mM Tris, pH 8.0, 300 mM NaCl, 300 mM imidazole). The eluted proteins were further concentrated using ultra centrifugal filter units (Millipore).

For obtaining purified TECPR2_TECPR (935-1411 aa) protein fragment without the MBP tag for use in GUVs assays, the eluted MBP-TECPR2_TECPR (935-1411 aa) fragment was cleaved with Factor Xa enzyme (Sigma-Aldrich) by incubating with the protein at 4°C overnight. The cleaved MBP tag and Factor Xa enzyme was then removed using size exclusion chromatography (Sephadex G-75 column; Cytiva). The purification of recombinant human Rab5a protein that was used for maleimide labelling in GUVs assay was performed as previously described (Tremel et al., 2021).

To check binding of GST and GST-Rabs (WT or mutants) with MBP-TECPR2_TECPR (935- 1411 aa) or GST and GST-SNX17 with MBP or MBP-TECPR2_TECPR (WT or R1336W), 5 µg of GST or GST-fusion proteins immobilized on glutathione-conjugated-agarose beads were incubated with 2.5 µg of MBP or MBP-fusion proteins prepared in TAP lysis buffer (20 mM Tris pH 8.0, 150 mM NaCl, 0.5% NP-40, 1 mM MgCl2, 1 mM Na3VO4, 1 mM NaF, 1 mM PMSF, and protease inhibitor cocktail) for 2 hr at 4°C on a HulaMixer. Following the incubation step, beads were gently centrifuged, and washed three times with TAP lysis buffer containing 0.1% NP-40. The protein complexes were eluted by boiling samples in 4X sample loading buffer at 100°C for 10 min and are subjected to SDS-PAGE, followed by immunoblotting as described above.

For the pulldown assay using the purified recombinant GST- or MBP-tagged fusion proteins and mammalian cells as a source of lysates, the HEK293T cells alone or transfected with indicated plasmids were lysed in ice-cold TAP lysis buffer (20 mM Tris-Cl pH 8.0, 150 mM NaCl, 0.5% NP-40, 1 mM MgCl2, 1 mM Na3VO4, 1 mM NaF, 1 mM PMSF, and protease inhibitor cocktail) at 4°C for 10 min, followed by centrifugation at 16,000×g for 5 min at 4°C. The lysates were collected and incubated with GST and GST-fusion protein or with MBP and MBP-fusion protein bound to the glutathione and amylose beads, respectively, for 2-3 hr at 4°C with rotation using HulaMixer. After incubation, beads were washed at least six times with TAP lysis buffer containing 0.1% NP-40, and protein complexes were eluted by boiling samples in 4X sample loading buffer at 100°C for 10 min before loading them onto SDS-PAGE gel and immunoblotting, as previously described.

### Purification of ferrofluid (FF)-containing endosomes

Ferrofluid (FF)-containing endosomes were purified using a previously described protocol (Walia et al., 2024). For a fully confluent 60-mm cell culture dish, FF-containing media was prepared by adding 9 μL FF (EMG 508) to 1.5 mL of plain pre-warm DMEM media kept at 37°C. The suspension was sonicated for 30 sec and filter-sterilized using a 0.2 μm pore-size filter (Thermo Scientific). For uptake (pulse) of FF, the 60-mm dish containing HeLa cells were washed once with 1X PBS, and 1.5 mL of FF-containing DMEM was added to the cells for 30 min at 37°C in a cell culture chamber. After the pulse period, cells were washed twice with 1X PBS and incubated with 10% FBS-containing DMEM media at 37°C for the indicated time points (chase). At the end of the chase time points, cells were washed once with 1X PBS, harvested and homogenized (∼20 strokes) in homogenization buffer (250 mM sucrose, 20 mM HEPES, 0.5 mM EGTA pH 7.2, and protease inhibitor cocktail) using a Dounce homogenizer (Sigma-Aldrich) on ice. The homogenates were centrifuged at 800 × g for 5 min at 4°C and the resultant PNS were collected and placed on a DynaMag-2 magnet stand (Invitrogen) for 60 min at 4°C. The supernatants were carefully removed and the FF-containing compartments were gently washed once with the homogenization buffer and sedimented by centrifugation. The isolated FF-containing endosomes were suspended in 1X Laemmli buffer, boiled for 10 min at 100°C, and analyzed by SDS-PAGE and immunoblotting, as described earlier.

### Flow cytometry analysis of surface protein markers

To detect the surface level of different receptors (Itgβ1, CI-M6PR, and LAMP1) in HeLa cells treated with either control siRNA or TECPR2 siRNA, cells were detached from the cell culture dish using a 5 mM EDTA solution prepared in 1X PBS and added to the cells for 15-20 min on ice. The cells were collected by a gentle pipetting in a fresh microcentrifuge tube and centrifuged at 400×g for 2 min at 4°C. The cell pellets were washed once with ice-cold flow cytometry buffer (0.2% FBS in 1X PBS), followed by incubation with primary antibody solution prepared in flow cytometry buffer for 1 hr on ice. The cells were washed once to remove unbound primary antibody and allowed to incubate with Alexa Fluor-conjugated secondary antibody solution made in flow cytometry buffer for 30 min on ice. Finally, the cells were washed and resuspended in flow cytometry buffer containing 1% paraformaldehyde (PFA) (Sigma-Aldrich). The amount of surface receptor levels was measured by fluorescence flow cytometry using a BD FACS Aria Fusion Cytometer and BD FACS Diva software version 8.0.1 (BD Biosciences) were used to acquire the samples. Data analysis was performed using the BD FlowJo version 10.0.1.

For determining the surface level of EGFR, the above-described methodology was used with some modifications. Briefly, the cells were serum starved by incubating in DMEM media for 30 min at 37°C, followed by fixation on ice for 15 min using 2.5% PFA solution prepared in 1X PBS. The cells were scraped out and resuspended in flow cytometry buffer containing primary antibody against EGFR to label the surface EGFR. The cells were washed once to remove unbound primary antibody, and allowed to incubate with Alexa Fluor-conjugated secondary antibody solution made in flow cytometry buffer for 30 min on ice. Finally, the cells were washed and resuspended in flow cytometry buffer and analyzed by flow cytometry.

### Quantification of focal adhesion number

To quantify the number of focal adhesions (FA), HeLa cells grown on glass coverslips and treated with indicated siRNA were fixed and processed for immunofluorescence staining using the protocol described above. To label FA, cells were stained using an anti-paxillin antibody. To quantify the number of FA, MaxEntropy thresholding was applied to the confocal microscopy images of paxillin staining in ImageJ software. The FA number was then determined using the ‘analyze particles’ function.

To determine the number of FA after cell spreading, HeLa cells treated with indicated siRNA were trypsinized and re-seeded on glass coverslips coated with 10 µg/mL fibronectin (Sigma-Aldrich). After 90 min post-seeding, cells were fixed and stained with an anti-paxillin antibody, and the number of FA was measured as described above.

### Integrin recycling assay

Using a previously described methodology (Steinberg et al., 2012), a recycling assay with an antibody against active (12G10) Itgβ1 was performed. Briefly, HeLa cells growing on glass coverslips and transfected with indicated siRNA were incubated for 1 hr at 37°C with 5 µg/mL of primary antibody against active (12G10) Itgβ1 prepared in DMEM media containing 10% FBS. The cells were washed with ice-cold 1X PBS, and the surface-bound non-internalized antibodies were stripped using ice-cold citric acid buffer (pH 4.5) for 5 min. The cells were washed two times with ice-cold 1X PBS and incubated with pre-warmed 10% FBS containing DMEM media for 30 min to chase the internalized antibody. To determine the primary antibody-bound Itgβ1 pool that was recycled back to the cell surface, the cells were washed in ice-cold 1X PBS and fixed using a 4% PFA solution for 20 min on ice. After the fixation step, cells were incubated with blocking buffer (1% BSA in 1X PBS) for 10 min and stained using Alexa Fluor 488-conjugated secondary antibody prepared in blocking buffer for 30 min at RT. To determine the internal pool of primary antibody-bound Itgβ1, cells were subjected to a second acid wash step after a 30-min chase period to strip the recycled antibody from the cell surface. Cells were then fixed with 4% PFA on ice for 20 min, permeabilized with 0.1% saponin-containing blocking buffer, and stained for 30 min at RT using Alexa Fluor 488-conjugated secondary antibody and DAPI prepared in blocking buffer. Representative images were then acquired using an LSM 710 Zeiss confocal microscope.

To quantify recycling of Itgβ1 using confocal microscopy images, shell analysis was performed, as described previously (Williamson et al., 2022), with some modifications. Briefly, a region of interest (ROI) covering the periphery (shape) of each selected cell was drawn using the freehand selection tool. With the clear outside function of Fiji software, Itgβ1 signals from nearby cells were removed, and the total Itgβ1 signal intensity was measured for the first ROI. The same ROI was then reduced by 2 µm, and Itgβ1 signal intensity was measured for the second ROI. Finally, surface Itgβ1 signal intensity was calculated by subtracting the intensity of the second ROI from the first ROI. For the analysis of internal Itgβ1, an ROI covering the periphery of each selected cell was drawn using the freehand selection tool. With the clear outside function of Fiji software, Itgβ1 signals from nearby cells were removed and Itgβ1 signal intensity was measured. The ratio of the surface to internal Itgβ1 signal intensity was calculated to determine the fold change in recycling.

To determine the colocalization of internal Itgβ1 with LAMP1, the above-mentioned antibody-based recycling assay was performed, and PCC was determined using the JaCoP plugin of ImageJ software as described previously.

### Transferrin recycling assay

To measure transferrin (Tfn) recycling, HeLa cells treated with control and TECPR2 siRNA were serum starved for 30 min at 37°C by addition of DMEM media to culture dishes. The cells were washed once with 1X PBS and pulsed for 5 min at 37°C with 20 µg/mL of Alexa 568-Tfn (Molecular Probes) prepared in DMEM media. After the pulse period, unbound or non-internalized Alexa 568-Tfn was removed by incubating the cells in citric acid buffer, pH 4.5 (0.1 M citric acid anhydrous and 0.1 M tri-sodium citrate dihydrate) for 90 sec at 37°C. After one wash with 1X PBS, for pulse-only samples, cells were collected by trypsinization, pelleted, and resuspended in flow cytometry buffer (0.2% FBS in 1X PBS) for measurement of Alexa 568-Tfn fluorescence intensity by flow cytometry. For the “chase” samples, after the citric acid buffer (pH 4.5) wash step, pre-warmed DMEM media containing 10% FBS was added to the cells for different time periods at 37°C. After the indicated chase period, a second citric acid buffer (pH 4.5) wash was given to cells for 90 sec at 37°C to strip off surface recycled Alexa 568-Tfn at each time point. Cells were collected by trypsinization, pelleted, and resuspended in flow cytometry buffer (0.2% FBS in 1X PBS) for measurement of remaining intracellular Alexa 568-Tfn fluorescence intensity by flow cytometry using a BD FACS Aria Fusion Cytometer, and BD FACS Diva software version 8.0.1 (BD Biosciences) were used to acquire the samples. Data analysis was performed using the BD FlowJo version 10.0.1.

To visualize Tfn recycling by microscopy, cells seeded on coverslips were processed using the above-described methodology. At the end of the pulse and chase time points, cells were fixed, processed, and imaged using confocal microscope.

### GUVs preparation and Rab5 labelling of GUVs

For GUVs, the lipid mixture was made with the following composition: DOPC (83.39 mol%), DOPE-MCC (15.55 mol%), Liss Rhod PE (0.05%), and DSPE-PEG Biotin (1 mol%). To prepare GUVs, 10 μL of the lipid mixture (∼ 1 mM) dissolved in chloroform was placed on Indium-Tin-Oxide (ITO)-coated plates. The lipid-coated plates were then dried in a desiccator for 45 min in vacuum condition. The GUVs were then made in a swelling solution (10 mM HEPES) that is inserted inside the electroformation chamber with a syringe, and the function generator (Textronix AFG3022C) was connected to copper strips of slides. The lipid film was then electroswelled at 25°C using a sinusoidal electric field of a sequence as previously described (Bhatia et al., 2017). The GUV produced was then transferred to a 1.5 mL microcentrifuge tube coated with 5 mg/mL

BSA (Sigma-Aldrich). Before labeling the GUVs with Rab5, immobilization of the GUVs was performed. An O-ring chamber attached to a glass slide was coated with avidin solution (1 mg/mL of avidin egg white (Invitrogen) prepared in 1X PBS and 5 mg/mL of BSA) for 2 hr, then washed twice with 100 µL of observation buffer (25 mM HEPES (pH 7.4), and 271.4 mM NaCl). Finally, 100 µL of observation buffer was added into the chamber, followed by the addition of 250 µL of the prepared GUVs and kept for 4 hr for incubation. The human Rab5a construct that was used for maleimide labelling was generously provided by Prof. Roger Williams (MRC Laboratory of Molecular Biology, Cambridge, UK). The purification of recombinant Rab5a and its anchoring to GUVs was performed using a strategy described in a previous study (Tremel et al., 2021). Briefly, two surface-exposed cysteine residues of Rab5a were mutated to C19S and C63S to facilitate the formation of a stable thioether bond between the C-terminal cysteine residues (Cys 212 residue) of human Rab5a and maleimide-functionalized (18:1 PE-MCC) lipid vesicles. 10 μM of purified recombinant Rab5a-GTP in buffer (25 mM HEPES, 150 mM NaCl, 0.5 mM TCEP) was then added onto the immobilized GUVs and incubated at 25°C for 4 hr. The unbound Rab5 was then removed carefully by adding and taking out 100 µL of wash buffer (31.8 mM HEPES, pH 8.0, 172.7 mM NaCl, 5 mM β-mercaptoethanol, and 181.8 mM sucrose) four times. The covalently attached Rab5a-GTP was visualized by incubating GUVs with antibody against Rab5 for 2 hr at 25°C in wash buffer, followed by washing four times with wash buffer, and finally incubating with Alexa-Fluor 488 conjugated secondary antibody for 1 hr at 25°C. To visualize the recruitment of TECPR2_TECPR (WT) and TECPR2_TECPR (R1336W) onto Rab5a-GTP-bound immobilized GUV, 20 μM of these purified proteins were added to buffer (265.5 mM HEPES, 150 mM NaCl, 0.5 mM TCEP) and incubated at 25°C for 2 hr. After incubation, the unbound antibodies were removed by adding and taking out 100 µL of wash buffer. The binding of the TECPR2_TECPR (WT) and TECPR2_TECPR (R1336W) was observed by incubating with antibody against TECPR2 at 25°C for 2 hr in wash buffer, followed by washing and finally incubating with Alexa Fluor-488 conjugated secondary antibody for 1 hr at 25°C.

### Zebrafish experiments

Adult zebrafishes were maintained at 26°C–28°C on a 14:10 hr light/dark cycle. Embryos for all assays were obtained by natural breeding of the wild-type fish.

### Total RNA isolation extraction

To determine the relative mRNA levels of tecpr2 in zebrafish embryos, total RNA was isolated from 20 embryos per experiment. The embryos collected at different days post-fertilization (dpf) were washed once with RNAse-free water and then stored in 200 µL TRIzol reagent (Invitrogen) at -80°C. The total RNA was isolated by homogenizing the embryos with tissue homogenizer, followed by the addition of 40 µL chloroform. The mixture was allowed to stand for 3 min at RT followed by centrifugation at 12000×g for 15 min at 4°C. A 40 µL of upper aqueous phase was transferred to chilled tubes containing 100 µL isopropanol. The above mixture was allowed to stand at 4°C for 15 min, followed by centrifugation and 70% ethanol wash to get RNA pellet. The dried RNA pellet was dissolved in 10 µL of RNAse-free water. The 5 µg of total RNA was used for cDNA synthesis using an equal volume of random hexamers and oligo-dT in a Superscript III reverse transcriptase reaction (Invitrogen).

### Quantitative RT-PCR analysis

Quantitative RT-PCR was performed with SYBR Green PCR Master Mix (Thermo Scientific) and tecpr2-specific primers (that binds to both tecpr2-201 and -202 transcripts). PCR conditions were used as per recommendations on a real-time PCR detection system (Eppendorf Master Cycler RealPlex4). The relative expression of mRNAs in embryos at different dpf was determined using the ΔΔCt method and normalized to β-actin mRNA levels.

tecpr2 qRT-Fp: 5’-TGGGAGCACATTCCAGGACTTC-3’

tecpr2 qRT-Rp: 5’-ATCCATCGGGGTCACTGCG-3’

β-actin qRT-Fp: 5’-GCAGAAGGAGATCACATCCCTGGC-3’

β-actin qRT-Rp: 5’-CATTGCCGTCACCTTCACCGTTC-3’

In situ hybridization

The 3’-end regions corresponding to tecpr2-201 and -202 transcripts was amplified using the primer pairs, as shown below. The reverse primer was appended with T3 promoter region.

Fp: 5’-GGAAAGACCTGTTCTGTATTTG-3’

Rp*: 5’-ctgaattaaccctcactaaaggTTAAATCACCTCCCACTCTTCC-3’

### *Lowercase shows the T3 promoter sequence

The purified PCR amplicons were used for in vitro transcription with T3 RNA polymerase (New England Biolabs) and digoxigenin-labelled rNTPs (Roche, 57127421) as per manufacturer’s instructions. The DIG-labelled RNA probes were used in the whole mount in situ hybridization (WISH) for the detection of tecpr2-201 and -202 transcripts. The bound probes were detected with alkaline phosphatase (AP)-tagged anti-DIG antibodies (Roche). The presence of mRNA-DIG labelled RNA probe-AP antibody complex was detected with NBT-BCIP substrate (Roche). The representative images were captured using stereozoom microscope with a 2.5× objective (Nikon).

### Morpholino injection and rescue

Different morpholino (MO) (purchased from Gene Tools, LLC) concentrations (0.25–1.0 mM) were micro injected (∼1 nL) into the fertilized zebrafish eggs at a one cell-stage. The transcript ID of the zebrafish tecpr2 transcripts and the morpholino sequence with its binding site are mentioned below:

Transcript ID: tecpr2-201 ENSDART00000104224.5 Transcript ID: tecpr2-202 ENSDART00000133668.4

MO-tecpr2-201: 5’-GATCAATAAAAGAGGTTGCTGTGTT-3’ (binds at start of intron 2) MO-tecpr2-202: 5’-CGTTTACATTGATTCTGAAGCCTAC-3’ (binds at exon 1)

Live and dead embryos were counted manually at 24 hr post-fertilization (hpf) to calculate survival percentage. Hatching percentage was determined by manually counting hatched and unhatched embryos at 72 hpf. The motility defects and phenotypic changes were assessed at 96 hpf using stereo zoom microscope at 0.5× magnification (Nikon, SMZ45T, Michrome6 camera). The rescue of tecpr2 morphants was done by injecting the embryo with in vitro transcribed tecpr2-201 mRNA (100 ng). For in vitro transcription, the CDS region of tecpr2-201 transcript was cloned downstream of HA-tag in pGADT7 vector. The plasmid was linearized by digestion with XhoI restriction enzyme (NEB) and was used as template in T7-mediated in vitro transcription as per manufacturer’s instructions (MEGAscript T7 Transcription Kit) (Thermo Scientific).

### Determining the knockdown efficiency by morpholinos

To check the knockdown efficiency of MO tecpr2-201 and tecpr2-202, zebrafish embryos were injected with GFP-expressing reporter constructs (containing the ztecpr2 MO-target sequences upstream of the GFP coding sequence in pEGFP-N1 vector) along with the respective morpholinos at a one-cell stage. The GFP expression was detected under fluorescent stereo zoom microscope (Nikon) and images were captured using a 20× objective. All the representative confocal images were adjusted for brightness and contrast using the ImageJ software.

### Whole-mount immunofluorescence of zebrafish embryos

Zebrafish embryos at one cell-stage were injected with MO-control, MO-tecpr2 (201+202) and MO-tecpr2 (201+202) with ztecpr2 mRNA. The embryos at 30 hpf were dechorionated and fixed in 4% PFA in 1X PBS overnight at 4°C. The presynaptic vesicles were detected by incubating the embryos with anti-synaptic vesicle protein 2 (SV2) (SV2; Developmental Studies Hybridoma Bank) primary antibody overnight at 4°C in HulaMixer followed by incubation with Alexa Fluor- 568-conjugated secondary antibody. SV2 antibody was diluted in 5% BSA prepared in 0.1% PBST. To detect postsynaptic acetylcholine receptors, the α-Bungarotoxin (α-BTX)-conjugated Alexa Fluor-488 was made in 1X PBS with 0.1% PBST and incubated along with secondary antibody for 2 hr at RT using a HulaMixer. The imaging of zebrafish was performed on Olympus IXplore SpinSR Spinning Disk Super Resolution Confocal Microscope using a 100× Apo/1.45 NA oil objective. All the representative confocal images were adjusted for brightness and contrast using ImageJ.

### Statistical analysis

All the data are presented as the mean ± S.D. unless otherwise specified. Statistical significance was determined using an unpaired two-tailed Student’s t test (GraphPad Prism 8.0) to calculate p values (****p<0.0001, ***p<0.001, **p<0.01, *p<0.05, or n.s., not significant (p>0.05)).

## Data availability

All relevant information supporting the findings of this study is presented in the manuscript and supplementary materials. A source data file comprising raw data and western blot images that have not been cropped is included in the manuscript.

## Abbreviations

APPL1, Adaptor protein, phosphotyrosine interacting with PH domain and leucine zipper 1; Arl, Arf-like; Arl8b, Arf-like GTPase 8b; BafA1, Bafilomycin A1; CI-M6PR, cation-independent mannose-6-phosphate receptor; EGFR, Epidermal growth factor receptor; ER, Endoplasmic reticulum; GDP, Guanosine diphosphate; GFP, Green fluorescent protein; GST, Gluathione-S- transferase; GTP, Guanosine triphosphate; GUVs, Giant unilamellar vesicles; HOPS, Homotypic fusion and protein sorting; HSAN, Hereditary sensory and autonomic neuropathies; IB, Immunoblotting; IP, immunoprecipitation; Itg, Integrin; LAMP1, Lysosomal-associated membrane protein 1; LE, Late endosome; LIR, LC3-interacting region; MBP, Maltose-binding protein; MO, Morpholino; NAD, Neuroaxonal dystrophy; PLEKHM1, Pleckstrin homology domain-containing family M member 1; RT, Room temperature; siRNA, short-interfering RNA; SNX, Sorting nexin; TECPR2, Tectonin-β- propeller repeat containing protein 2; Tfn, Transferrin; WT, Wild-type

## Acknowledgements

1. S. P. and K.W. acknowledges fellowship support from IISER Mohali and the Council of Scientific & Industrial Research (CSIR), respectively. R.P. acknowledges fellowship support from Anusandhan National Research Foundation (ANRF). P.S. and K.K.B. acknowledges post-doctoral fellowship support from DBT [DBT-RA/20123/July/N/4351] and IISER Mohali, respectively. This research was supported by the Department of Biotechnology (DBT)/Wellcome Trust India Alliance Senior Fellowship [IA/S/19/1/504270] and DBT grant [BT/PR53883/BMS/85/123/2024], and intramural funding from IISER Mohali to M.S. This work was supported by grants from ANRF [CRG/2022/003266] and CSIR [HCP532401] to A.T. This work was supported by grants from DBT to R.R. [BT/PR36570/BRB/10/1976/2021] and T. B. [BT/RLF/Re-entry/06/2020]. The funders had no role in the study design, data collection, interpretation, or decision to submit the manuscript for publication. The authors acknowledge all colleagues who shared the plasmids used in this study. The authors would also like to acknowledge Prateek Arora (IISER Mohali FACS Facility) for technical help with flow cytometry related experiments and all lab members of the M.S. and A.T. for helpful discussions.

## Contributions

S.P. designed and performed the majority of the experiments, analyzed the results, and prepared the figures. M.S., and A.T. conceived and designed the study and wrote the manuscript. R.P., K.W., and A.A. performed some of the protein-protein interaction assays and assisted in other experiments. P.S. designed, performed and analyzed the zebrafish experiments. R.R. designed and analyzed zebrafish experiments. K.K.B, R.G., and Y.V. contributed in generation of molecular biology reagents and assisted in the yeast-two hybrid assays. S.G. and T.B. prepared the GUVs and assisted in the GUV assay.

## Supplementary Figure Legends

**Supplementary Figure S1:**
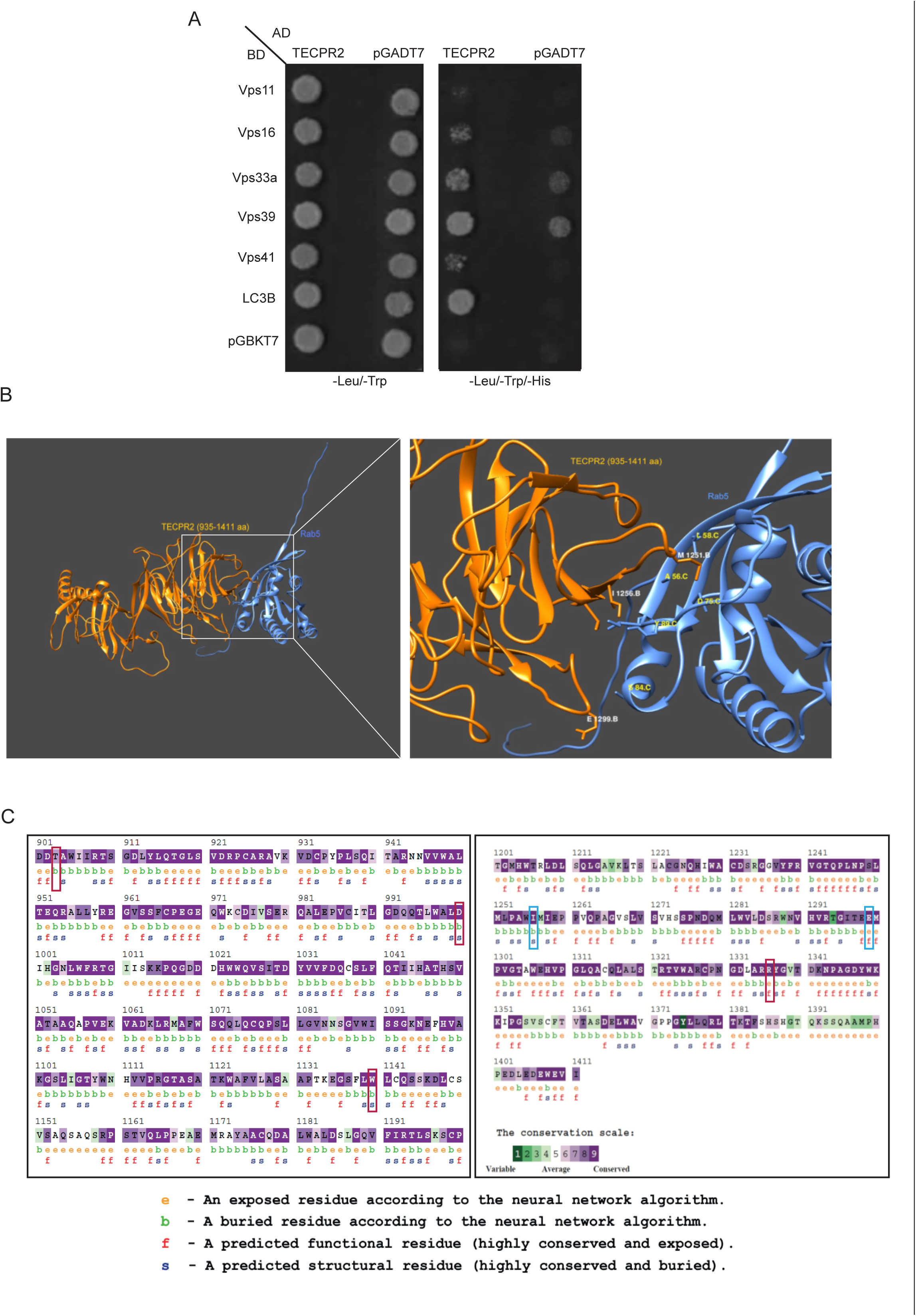
TECPR2 interacts with the HOPS complex subunits and AlphaFold2-based prediction of binding interface of TECPR2_TECPR with Rab5. A) Yeast two-hybrid interaction analysis of TECPR2 with HOPS complex subunits and LC3b. Co- transformants were spotted on -Leu/-Trp to confirm viability and -Leu/-Trp/-His media to detect interactions. B) Intramolecular interactions between Rab5 and the C-terminal region TECPR2 from 935-1411 amino acids were modeled using the AlphaFold2-ColabFold tool and visualized using Chimera software. The chain B in orange denotes TECPR2 (935-1411 aa fragment), and chain C in blue indicates Rab5. C) The Consurf Server tool was used to analyze conserved residues in the 901-1411 aa fragment of human TECPR2. The amino acid residues highlighted in red boxes are a subset of HSAN9-associated variants, and the highlighted blue boxes indicate residues identified from AlphaFold2 analysis described in (B).

**Supplementary Figure S2:**
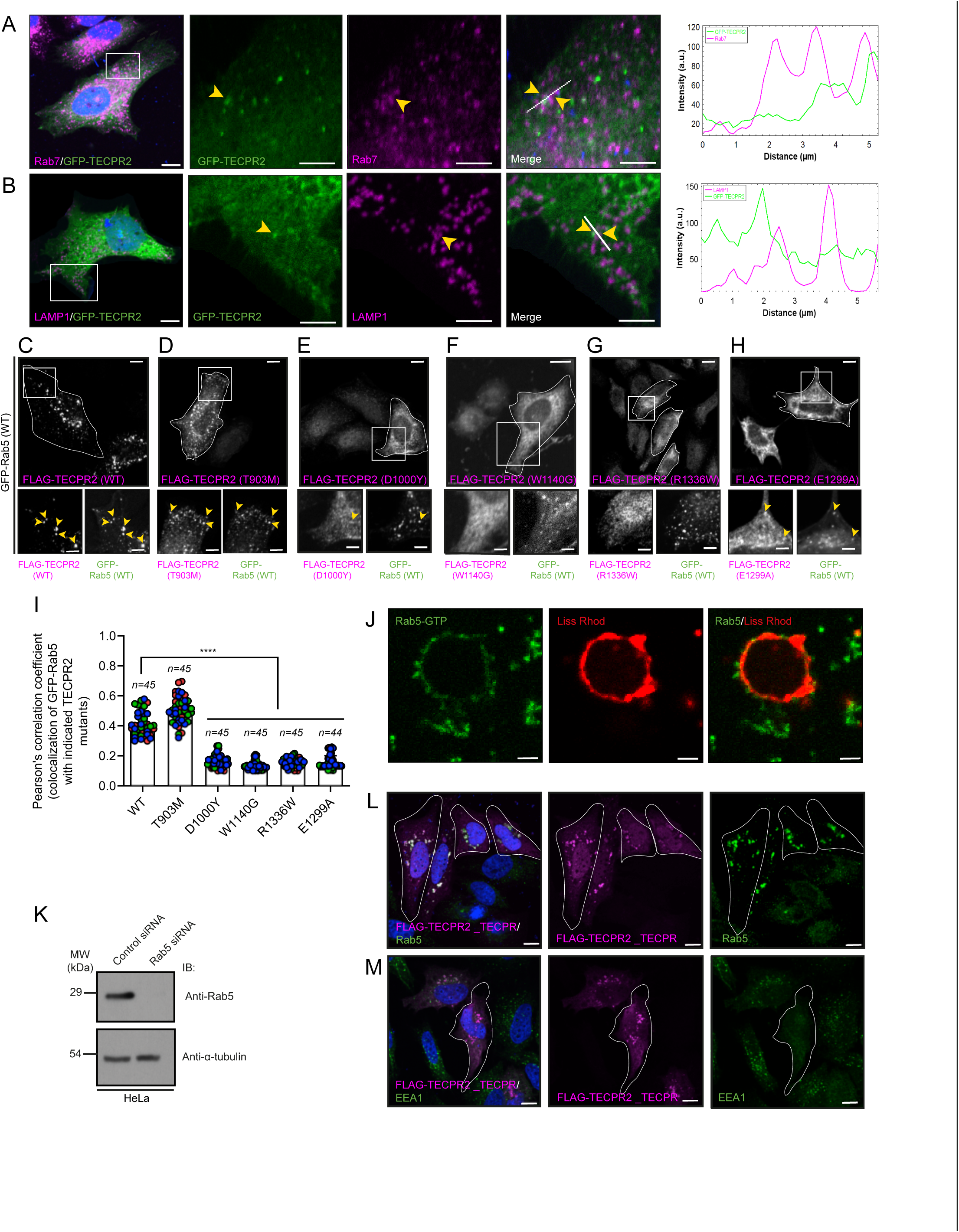
Rab5 binding is required for TECPR2 membrane localization. A and B) Representative confocal micrographs of HeLa cells transfected with GFP-TECPR2 and immunostained with anti-Rab7 or anti-LAMP1 antibodies. The line profiles on the right indicate fluorescence intensity along the white lines for both channels: GFP-TECPR2 (green) and Rab7 (magenta) (A) or LAMP1 (magenta) (B). The arrowheads in the insets denote the colocalized pixels. Scale bars: 10 µm (main); 5 µm (inset). C-I) Representative confocal micrographs of HeLa cells co-transfected with GFP-Rab5 and FLAG-TECPR2 (WT) (C), FLAG-TECPR2 (T903M) (D), FLAG-TECPR2 (D1000Y) (E), FLAG-TECPR2 (W1140G) (F), FLAG-TECPR2 (R1336W) (G), or FLAG-TECPR2 (E1299A) (H). Cells were fixed and immunostained using an anti-FLAG antibody (magenta). The arrowheads in the insets denote the colocalized pixels of GFP-Rab5 with FLAG-TECPR2 (WT) or indicated mutant. Scale bars: 10 µm (main); 5 µm (inset). In (I), quantification of the Pearson’s colocalization coefficient for GFP-Rab5 with FLAG-TECPR2 (WT) or indicated mutant is shown. n denotes the total number of cells examined in three independent experiments. Experiments are color-coded, and each dot represents the individual data points from each experiment (****p<0.0001; unpaired two-tailed Student’s t test). J) Confocal micrographs of Liss-Rhodamine-labeled GUVs incubated with purified GTP-loaded Rab5. The GUVs were immunostained with an anti-Rab5 antibody to determine the recruitment of Rab5 on GUVs. Scale bars: 10 μm. K) To determine knockdown efficiency of Rab5 siRNA, lysates of HeLa cells treated with control siRNA and Rab5 siRNA were immunoblotted with the indicated antibodies. L and M) Representative confocal images of HeLa cells expressing FLAG- TECPR2_TECPR (935-1411 aa) (highlighted with a white line) and immunostained using the anti- Rab5 (L) and anti-EEA1 (M) antibodies. Scale bars: 10 µm.

**Supplementary Figure S3:**
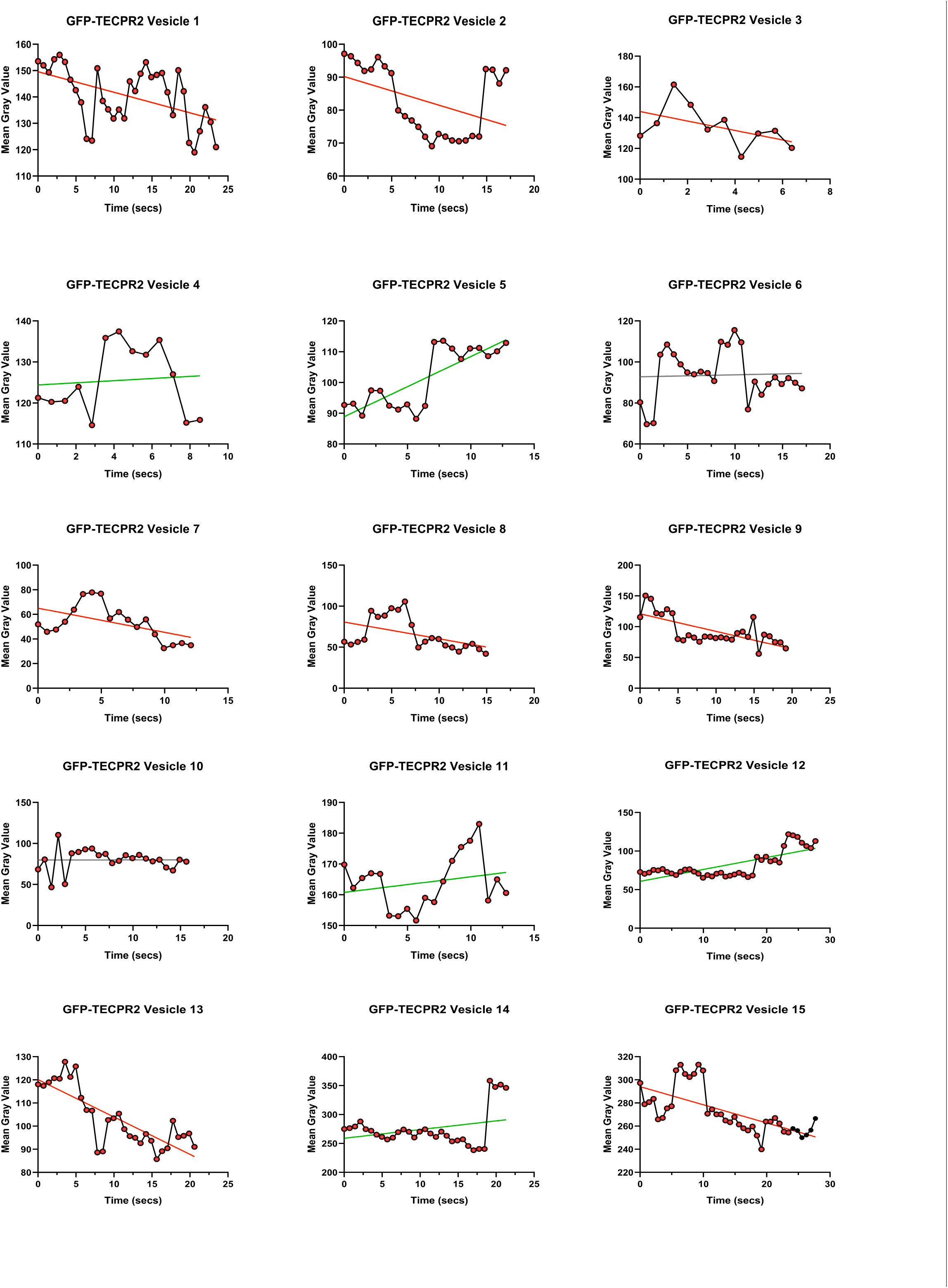
GFP-TECP2 vesicle intensity profile over time showing fission and fusion events. Graph showing changes in the fluorescence intensities of individual GFP- TECPR2 vesicles during fission and fusion processes (see Figure 3A-B and Supplementary Video S2). Linear regressions are shown for n=15 vesicles. Slightly positive and negative slopes are marked in green and red, respectively, and the flat lines are indicated in grey.

**Supplementary Figure S4:**
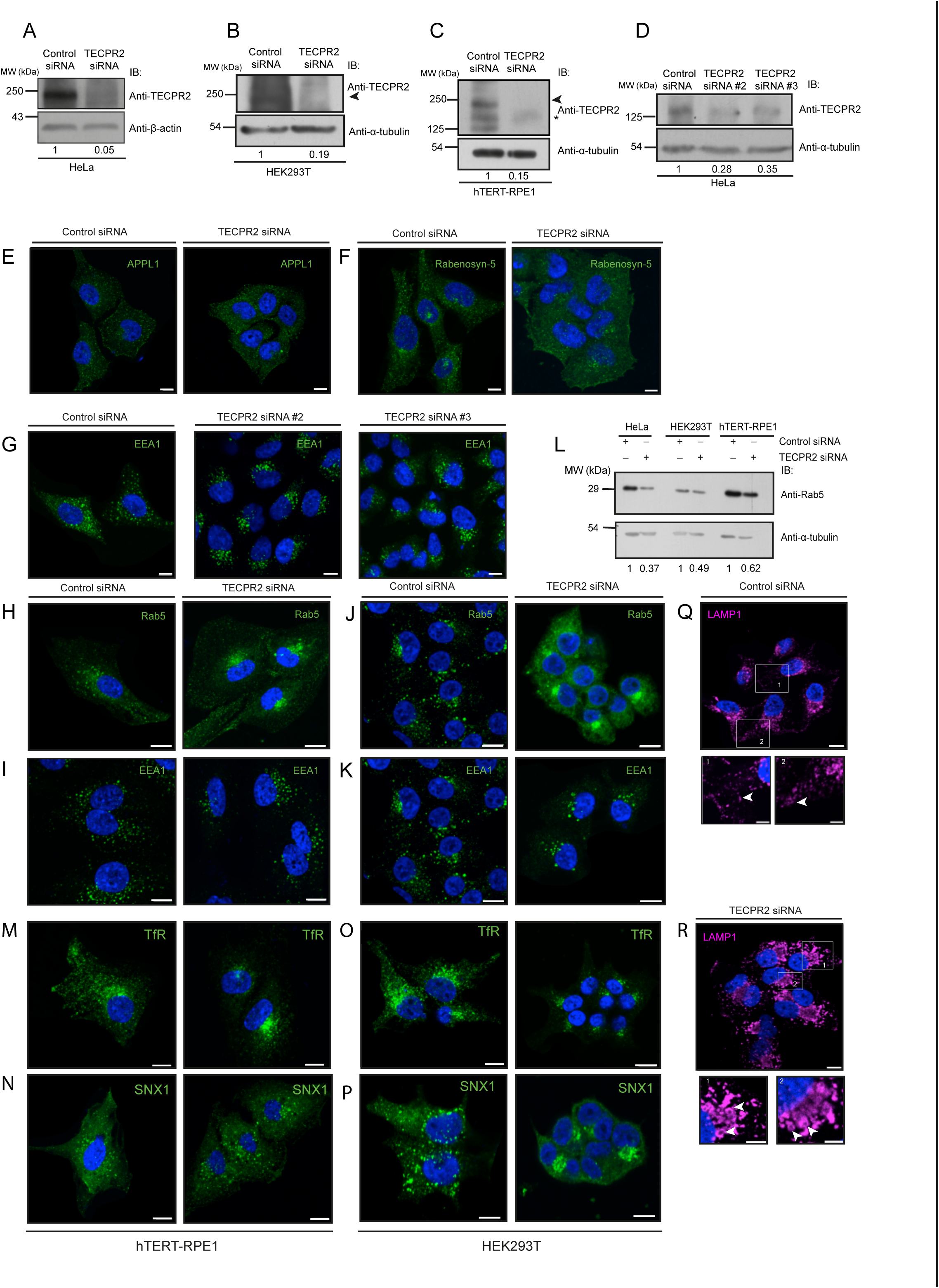
TECPR2 maintains the cellular distribution of early endosomes. A-D) Lysates of indicated cell type treated with control siRNA and TECPR2 siRNA were immunoblotted (IB) with anti-TECPR2 antibody for determining the knockdown efficiency, and the levels of β-actin or α-tubulin were used as a loading control. The band of endogenous TECPR2 is marked with an arrowhead, and the (*) asterisk indicates a non-specific band detected by the anti-TECPR2 antibody. The values represent a densitometric analysis of the TECPR2 signal intensity normalized to loading controls. E and F) Representative confocal micrographs of HeLa cells treated with the indicated siRNA, followed by immunostaining with the anti-APPL1 (E) or anti-Rabenosyn-5 (F) antibodies. Scale bars: 10 µm. G) Representative confocal micrographs of HeLa cells treated with the indicated siRNA, followed by immunostaining with anti-EEA1 antibody. Scale bars: 10 µm. H-K) Representative confocal micrographs of hTERT-RPE1 cells (H-I) and HEK293T cells (J-K) treated with control siRNA and TECPR2 siRNA, followed by immunostaining with the anti-Rab5 or anti-EEA1 antibodies. Scale bars: 10 µm. L) Lysates of indicated cell types transfected with control siRNA and TECPR2 siRNA were IB with indicated antibodies. The values represent the densitometric analysis of the Rab5 signal intensity normalized to the signal intensity of α-tubulin used as a loading control. M-P) Representative confocal micrographs of hTERT-RPE1 cells (M-N) and HEK293T cells (O-P) treated with control siRNA and TECPR2 siRNA, followed by immunostaining with the anti-TfR or anti-SNX1 antibodies. Scale bars: 10 µm. Q and R) Representative confocal micrographs of HeLa cells treated with the indicated siRNA, followed by immunostaining with an anti-LAMP1 antibody. The arrowheads in the insets denote the LAMP1-positive endosomes. Scale bars: 10 µm (main); 5 µm (inset).

**Supplementary Figure S5:**
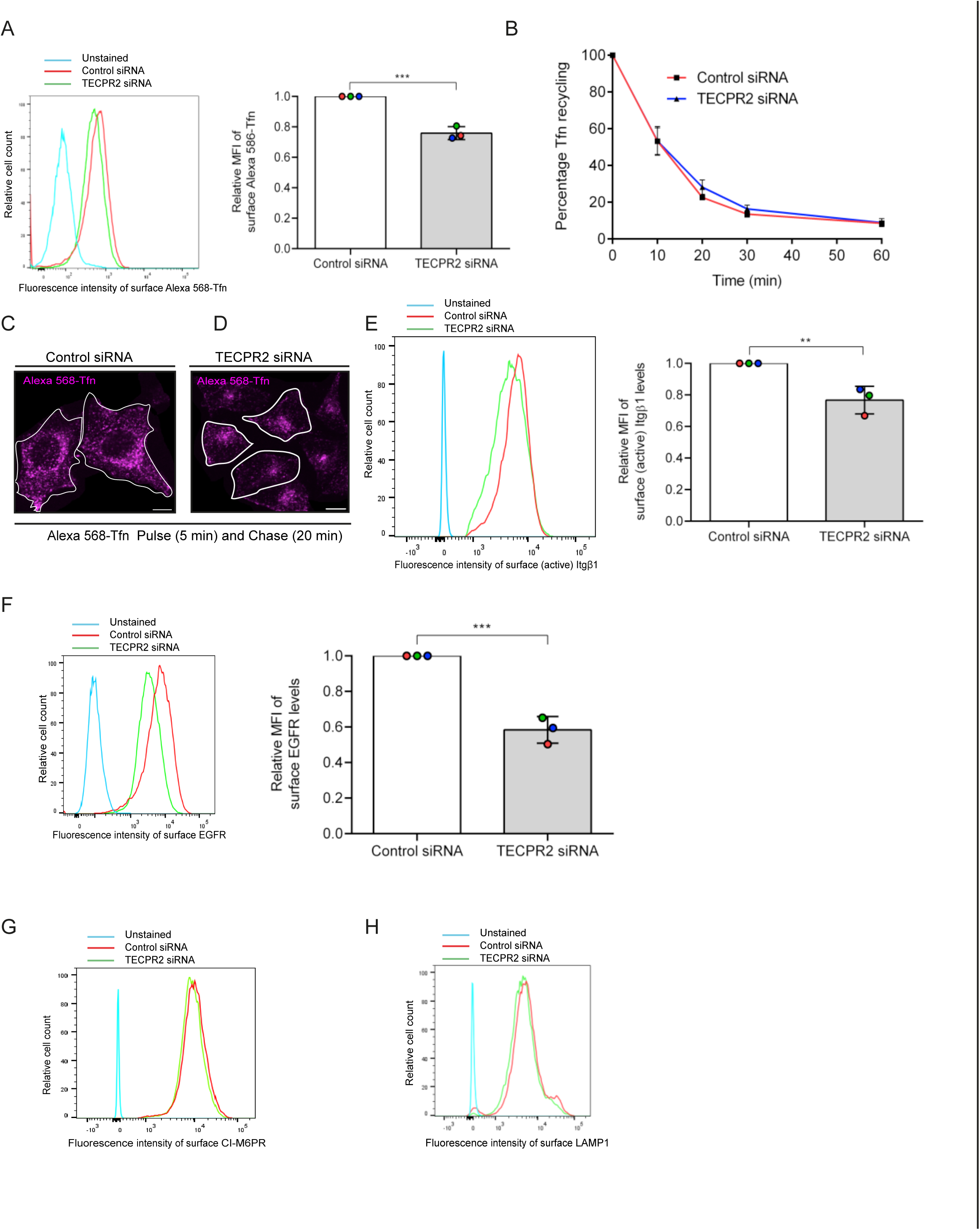
TECPR2-depleted cells exhibit reduced levels of Itgβ1 and EGFR at the cell surface. A) A histogram showing surface levels of TfR (labeled by the Alexa 568-Tfn ligand) in HeLa cells upon control and TECPR2 siRNA treatment, analyzed by flow cytometry. The graph on the right represents Mean Fluorescence Intensity (MFI) for surface-bound Alexa 568-Tfn signal (***p<0.001; unpaired two-tailed Student’s t test). B) Transferrin recycling assay. HeLa cells treated with either control siRNA or TECPR2 siRNA were serum starved for 30 min. Cells were incubated with Alexa 568-Tfn for 5 min (pulse only) or chased in complete media for the indicated times. Flow cytometry analysis was then performed to measure remaining Alexa 568- Tfn at the indicated times. C and D) HeLa cells treated with control siRNA and TECPR2 siRNA were serum starved for 30 min and pulsed with Alexa 568-Tfn for 5 min and chased in complete media for 20 min at 37°C prior to fixation. Scale bars: 10 µm. E) A histogram showing fluorescence intensity of surface Itgβ1 (active; 12G10) levels in control siRNA and TECPR2 siRNA-treated HeLa cells as analyzed by flow cytometry. The graph on the right represents relative MFI of surface Itgβ1 levels (**p=0.01; unpaired two-tailed Student’s t test). F) A histogram showing fluorescence intensity of surface EGFR levels in control siRNA and TECPR2 siRNA-treated HeLa cells as analyzed by flow cytometry. The graph on the right represents relative MFI of surface EGFR levels (***p<0.001; unpaired two-tailed Student’s t test). G and H) Representative histograms showing fluorescence intensity of surface CI-M6PR and LAMP1 levels in control siRNA and TECPR2 siRNA-treated HeLa cells as analyzed by flow cytometry.

**Supplementary Figure S6:**
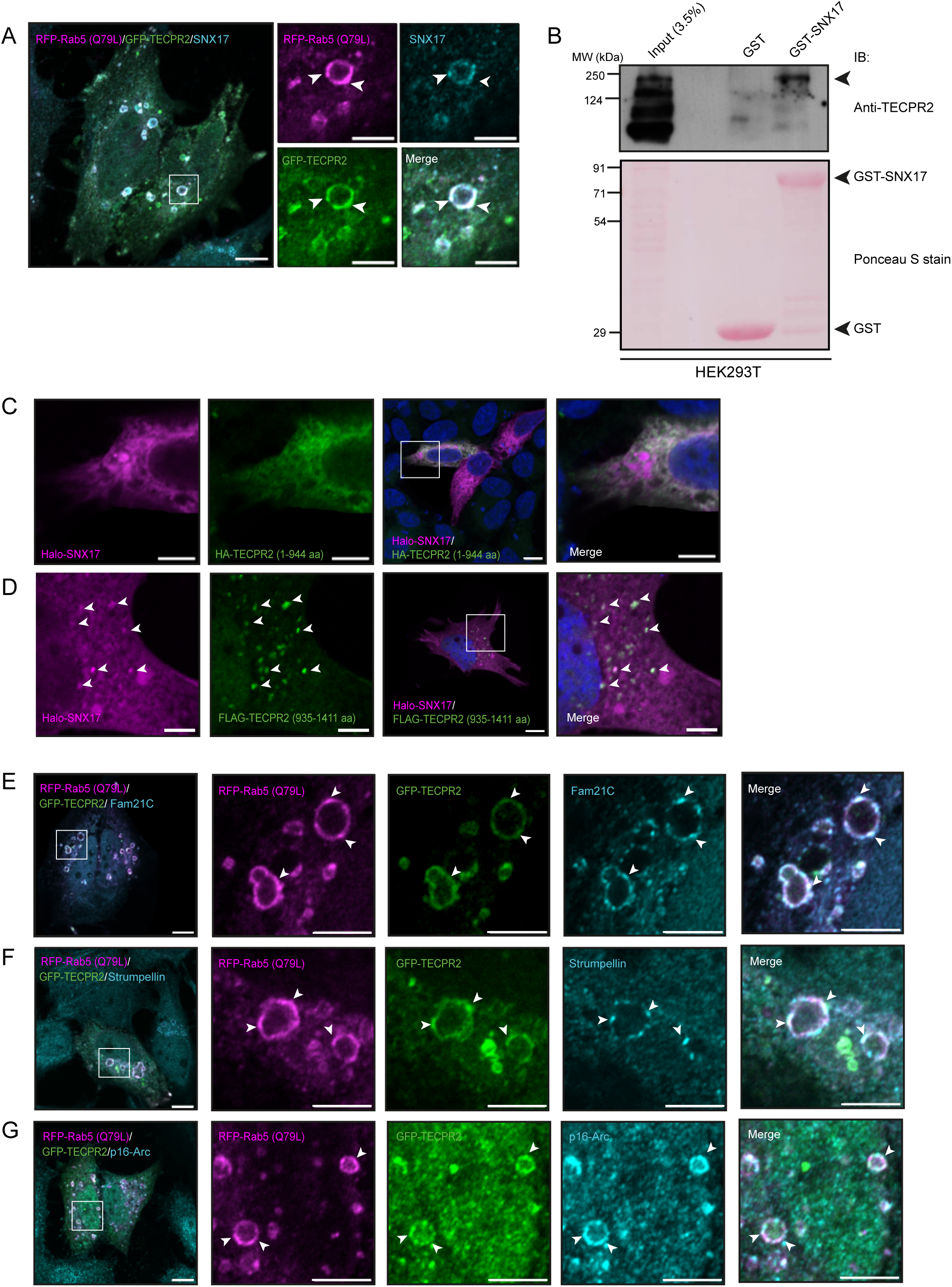
TECPR2 interacts with SNX17 and colocalizes with SNX17 and WASH Complex subunits on early endosomes. A) Representative Airyscan super-resolution image of HeLa cells co-transfected with GFP-TECPR2 and RFP-Rab5 (Q79L) expressing plasmids and immunostained with anti-SNX17 antibody. The arrowheads denote co-localized pixels for all three proteins. Scale bars: 10 µm (main); 5 µm (inset). B) Recombinant GST and GST-SNX17 proteins immobilized on glutathione-conjugated beads were incubated with the HEK293T cell lysates. The precipitates were IB with an anti-TECPR2 antibody, and Ponceau S staining was done to visualize the purified proteins. C and D) Representative confocal images of HeLa cells co-transfected with plasmids expressing Halo-SNX17 with HA-TECPR2 (1-944 aa) (C) or with FLAG-TECPR2 (935-1411 aa) (D). The cells were fixed and immunostained with the anti-HA or anti-FLAG antibodies. To visualize Halo-SNX17, cell-permeant HaloTag TMR ligand (71 nM) was used. The arrowheads denote co-localization of Halo-SNX17 with FLAG-TECPR2 (935-1411 aa). Scale bars: 10 µm (main); 5 µm (inset). E-G) Representative Airyscan super- resolution images of HeLa cells co-transfected with GFP-TECPR2 and RFP-Rab5 (Q79L) expressing plasmids and immunostained for Fam21C (E), Strumpellin (F), and p16-Arc (G). The arrowheads denote co-localized pixels for all three proteins. Scale bars: 10 µm (main); 5 µm (inset).

**Supplementary Figure S7:**
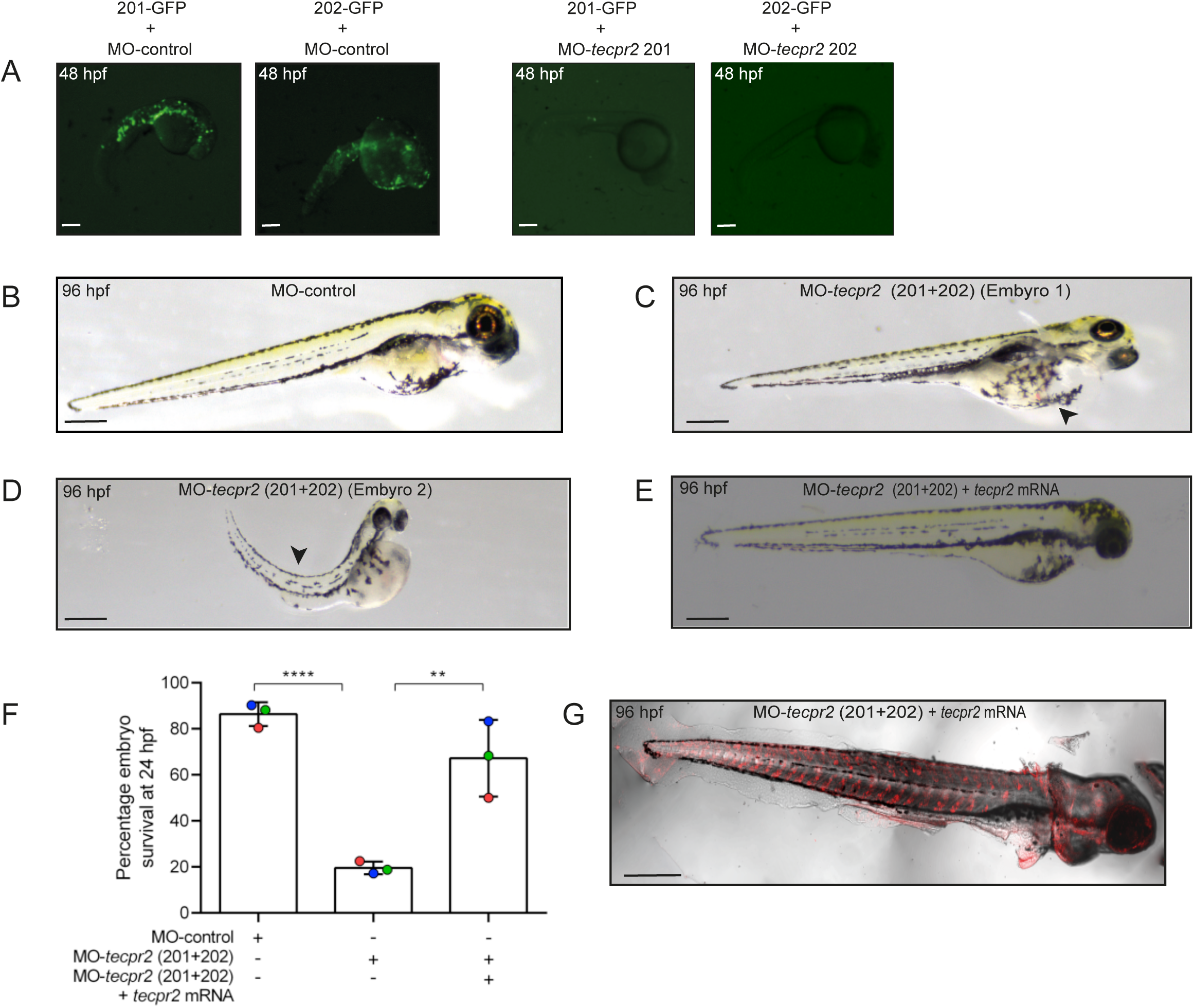
TECPR2-depleted zebrafish embryos show tail curvature and pericardial oedema. A) Representative images of zebrafish embryos acquired at 48 hpf after injection with a GFP-expressing reporter construct (containing the ztecpr2 MO-target sequences (201 or 202) upstream of the GFP tag) along with the respective morpholinos (targeting the ztecpr2) at a single-cell stage to determine the knockdown efficiency of the morpholinos. Scale bars: 200 µm. B-E) Lateral view of zebrafish embryos at 96 hpf upon injection with MO-control (B), MO-tecpr2 (201+202) (C and D), and MO-tecpr2 (201+202) + tecpr2 mRNA (rescue) (E). The tecpr2 morphants exhibited tail curvature and developed pericardial oedema, and the phenotype was rescued upon injection of tecpr2 mRNA into these morphants. Scale bars: 200 µm. F) The graph represents dose-dependent survival of zebrafish embryos at 24 hpf upon injection with MO-control, MO-tecpr2 (201+202), and MO-tecpr2 (201+202) + tecpr2 mRNA (rescue). G) At 96 hpf, whole-mount immunostaining of zebrafish embryos was performed with an anti-HA antibody to confirm the expression of tecpr2 mRNA (fused with a HA tag coding sequence) in embryos injected with MO-tecpr2 (201+202) along with tecpr2 mRNA (rescue). Scale bar: 10 µm.

## Supplementary Video Legends

Supplementary Video S1: Time-lapse imaging of HeLa cells expressing GFP-TECPR2 and mcherry-VAPB. The video is captured at 3.34 frames per second with no time interval between the frames. The video is shown at 5 frames/sec (the total number of frames displayed is 100). The yellow arrow indicates GFP-TECPR2-positive endosomes migrating on mcherry-VAPB tubules, and the blue arrow indicates the association and dynamics of GFP-TECPR2 endosomes positive for mcherry-VAPB at the ER-endosome contact sites. Scale bar: 10 µm.

Supplementary Video S2: Time-lapse imaging of HeLa cells showing peripheral and perinuclear vesicles of GFP-TECPR2. The video of HeLa cells transfected with GFP-TECPR2 and RFP-Rab5-expressing plasmids is captured at 2.84 frames per second with no time interval between the frames. The video is shown at 5 frames/sec (the total number of frames displayed is 50). The yellow and blue arrows indicate the dynamics of peripheral and perinuclear endosomes, respectively. The video does not include the RFP-Rab5 channel. Scale bars: 10 µm (main); 5 µm (inset).

Supplementary Video S3: Time-lapse imaging of HeLa cells expressing GFP-TECPR2 and RFP-Rab5, and displaying fission and fusion events of GFP-TECPR2-positive vesicles. The video is captured at 2.84 frames per second with no time interval between the frames. The video is shown at 5 frames/sec (the total number of frames displayed is 40 for fission events and 21 for fusion events). The yellow arrow denotes fission and fusion events of GFP-TECPR2-positive vesicles. Scale bar: 10 µm.

Supplementary Video S4: Time-lapse imaging of HeLa cells expressing GFP-TECPR2 and RFP-Rab5 and displaying stable tubules positive for GFP-TECPR2. The video is captured at 3.88 frames per second with no time interval between the frames. The video is shown at 5 frames/sec (the total number of frames displayed is 50). The yellow arrow indicates the dynamics and tubulation of GFP-TECPR2-positive vesicles. Scale bar: 10 µm.

Supplementary Video S5: Time-lapse imaging of HeLa cells expressing GFP-TECPR2 (R1336W) mutant and RFP-Rab5. The video is captured at 3.72 frames per second with no time interval between the frames. The video is shown at 5 frames/sec (the total number of frames

displayed is 30). The GFP-TECPR2 (R1336W) mutant primarily display cytosolic distribution. Scale bar: 10 µm.

Supplementary Video S6: Time-lapse imaging of HeLa cells expressing GFP-TECPR2 and untagged Rab5 and incubated with Alexa 568-Tfn. HeLa cells expressing GFP-TECPR2 and untagged-Rab5 were pulsed with 20 µg/mL of Tfn ligand conjugated to Alexa Fluor-568 (Alexa 568-Tfn) for 5 min prior to the start of the imaging. Following the pulse period, cells were washed, and imaging of internalized Alexa 568-Tfn for 30 min (chase) in complete media at 37°C was performed. The video is captured at 3 frames per second with no time interval between the frames. The video is shown at 5 frames/sec (the total number of frames displayed is 116). The yellow arrow indicates the dynamics of GFP-TECPR2 endosomes containing the cargo Alexa 568-Tfn, and the blue arrow indicates the tubulation and fission event of GFP-TECPR2 endosomes containing Alexa 568-Tfn. Scale bar: 5 µm.

Supplementary Video S7: Time-lapse imaging of control siRNA and TECPR2 siRNA-treated HeLa cells expressing SNX17-GFP with surface internalized Itgβ1. HeLa cells (control or TECPR2-siRNA-treated) were transfected with SNX17-GFP-expressing plasmids. Prior to the start of the imaging, the cells were incubated on ice for 1 hr with a complex of anti-Itgβ1 (12G10) primary antibody labeled with Alexa 568-conjugated secondary antibody to surface label Itgβ1. The cells were washed once and transferred to complete media for 10 min at 37°C to allow internalization of antibody-bound surface Itgβ1. Post-internalization step, the non-internalized antibody complex was removed by two washes with citric acid buffer (pH 4.5). After the acid wash step, imaging of surface internalized antibody-bound Itgβ1 was performed in complete media at 37°C. The control siRNA treated cell video is captured at 3.2 frames per second with no time interval between the frames, and the TECPR2 siRNA treated cell video is captured at 3.22 frames per second with no time interval between the frames. The videos are shown at 5 frames/sec (the total number of frames displayed is 31). While TECPR2 depletion reduces the membrane recruitment of SNX17-GFP on endocytosed Itg1-positive endosomes, the yellow arrows in control siRNA-treated cells indicate the recruitment of SNX17-GFP on endocytosed Itg1-positive endosomes. Scale bar: 5 µm.

Supplementary Video S8: Time-lapse imaging of HeLa cells expressing Halo-SNX17, GFP- TECPR2, and untagged Rab5 with surface internalized Itgβ1. HeLa cells were transfected with Halo-SNX17, GFP-TECPR2, and untagged Rab5-expressing plasmids. After 14 hr post- transfection, prior to the start of the imaging, the cells were incubated on ice for 1 hr with a complex of anti-Itgβ1 (12G10) primary antibody labeled with Alexa 568-conjugated secondary antibody to surface label Itgβ1. The cells were washed once and transferred to complete media for 10 min at 37°C to allow internalization of antibody-bound surface Itgβ1. Post-internalization step, the non- internalized antibody complex was removed by two washes with citric acid buffer (pH 4.5). After the acid wash step, imaging of surface internalized antibody-bound Itgβ1 was performed in complete media containing the cell-permeant Janelia Fluor 646 HaloTag ligand (40 nM) at 37°C. The video is captured at 1.28 frames per second with no time interval between the frames. The video is shown at 5 frames/sec (the total number of frames displayed is 30). The blue arrows indicate the endosomes positive for both GFP-TECPR2 and Halo-SNX17 and containing the surface-internalized Itgβ1. Scale bar: 10 µm.

Supplementary Video S9: Time-lapse imaging of control siRNA and TECPR2 siRNA-treated HeLa cells expressing GFP-WASHC1 and RFP-Rab5 (Q79L). HeLa cells (control or TECPR2- siRNA-treated) were transfected with GFP-WASHC1 and RFP-Rab5 (Q79L)-expressing plasmids. The control siRNA treated cell video is captured at 3.24 frames per second with no time interval between the frames, and the TECPR2 siRNA treated cell video is captured at 3.27 frames per second with no time interval between the frames. The videos are shown at 5 frames/sec (the total number of frames displayed is 31). Scale bar: 5 µm.

Supplementary Video S10: Zebrafish embryos injected with tecpr2-targeting morpholinos exhibit an impaired motility phenotype. As shown in the video, MO-control injected zebrafish embryos upon touch stimulation displayed normal flight response, whereas MO-tecpr2 (201+202) injected embryos displayed a delay in response to touch stimulation and a shiver response that does not generate enough force for the embryo to swim or execute a proper flight response. The injection of tecpr2 mRNA in tecpr2 morphants rescued this phenotype. The videos were captured using stereozoom microscope with 0.5X zoom.

## Supplementary Table Legends

**Supplementary Table I:** AlphaMissense score and pathogenicity prediction for TECPR2 clinical variants.

| TECPR2 variants | AlphaMissense Score | Pathogenicity Prediction |
| --- | --- | --- |
| p.Thr903Met | 0.13 | Likely benign |
| p.Asp1000Tyr | 0.958 | Likely pathogenic |
| p.Trp1140Gly | 0.903 | Likely pathogenic |
| p.Arg1336Trp | 0.968 | Likely pathogenic |

**Supplementary Table II:** List of DNA constructs used in this study.

| Plasmid Name | Description | Source |
| --- | --- | --- |
| ***Mammalian expression constructs:*** |  |  |
| pcDNA3.1(-) vector | Mammalian expression vector | Available in the lab |
| pcDNA3.1(-)-FLAG-TECPR2 (WT) | Full-length human TECPR2 (WT) with N-terminal FLAG tag cloned in pcDNA3.1(-) vector | This study |
| pcDNA3.1(-)-FLAG-TECPR2 (T903M) | Full-length human TECPR2 with T903M mutation and N-terminal FLAG tag cloned in pcDNA3.1(-) vector | This study |
| pcDNA3.1(-)-FLAG-TECPR2 (D1000Y) | Full-length human TECPR2 with D1000Y mutation and N-terminal FLAG tag cloned in pcDNA3.1(-) vector | This study |
| pcDNA3.1(-)-FLAG-TECPR2 (W1140G) | Full-length human TECPR2 with W1140G mutation and N-terminal FLAG tag cloned in pcDNA3.1(-) vector | This study |
| pcDNA3.1(-)-FLAG-TECPR2 (I1256A) | Full-length human TECPR2 with I1256A mutation and N-terminal FLAG tag cloned in pcDNA3.1(-) vector | This study |
| pcDNA3.1(-)-FLAG-TECPR2 (E1299A) | Full-length human TECPR2 with E1299A mutation and N-terminal FLAG tag cloned in pcDNA3.1(-) vector | This study |
| pcDNA3.1(-)-FLAG-TECPR2 (R1336W) | Full-length human TECPR2 with R1336W mutation and N-terminal FLAG tag cloned in pcDNA3.1(-) vector | This study |
| pcDNA3.1(-)-FLAG-TECPR2 (935-1411 aa) | N-terminal FLAG-tagged human TECPR2 from 935-1411 amino acid cloned in pcDNA3.1(-) vector | This study |
| pcDNA3.1(-)-HA-TECPR2 (1-944 aa) | N-terminal HA-tagged human TECPR2 from 1-944 amino acid cloned in pcDNA3.1(-) vector | This study |
| pcDNA3.1(-)-FLAG-TECPR2 (Rescue) | Full-length human TECPR2 rescue construct designed against TECPR2 siRNA using silent mutations and with N-terminal FLAG tag cloned in pcDNA3.1(-) vector | This study |
| pcDNA3.1(+)-Untagged-Rab5a (WT) | Full-length human Rab5 (WT) cloned in pcDNA3.1(+) vector | This study |
| pEGFP-C1 vector | Mammalian GFP expression vector | Gift from Prof. Peter Cullen (University of Bristol, UK) |
| pEGFP-C1-TECPR2 (WT) | Full length human TECPR2 (WT) with N-terminal GFP tag cloned in pEGFPC1 vector | This study |
| pEGFP-C1-TECPR2 (R1336W) | N-terminal GFP-tagged human TECPR2 with R1336W mutation and cloned in pEGFP-C1 vector | This study |
| pEGFP-C1-Rab5a (WT) | Full-length Rab5a (WT) with N-terminal GFP-tag cloned in pEGFP-C1 vector | Gift from Prof. Steve Caplan (University of Nebraska Medical Center, USA) |
| pEGFP-C1-Rab5a (Q79L) | Full-length Rab5 with Q79L mutation (GTP-locked) and cloned in pEGFP-C1 vector | Gift from Prof. Steve Caplan (University of Nebraska Medical Center, USA) |
| pEGFP-C1-Rab5a (S34N) | Full-length Rab5 with S34N mutation (GDP-locked) and cloned in pEGFP-C1 vector | Gift from Prof. Steve Caplan (University of Nebraska Medical Center, USA) |
| pMRFP-C3-Rab5a (WT) | Full-length Rab5a (WT) with N-terminal RFP tag cloned in pmRFP-C3 vector | Gift from Prof. Ari Helenius (Addgene plasmid # 14437) |
| pMRFP-C3-Rab5a (Q79L) | Full-length Rab5a (Q79L) with N-terminal RFP tag cloned in pmRFP-C3 vector | This study |
| pMRFP-C3-Rab5a (S34N) | Full-length Rab5a (S34N) with N-terminal RFP tag cloned in pmRFP-C3 vector | This study |
| pcDNA3.3-Rab5A (Q79L)-BirA*-HA-MAO | Rab5a lacking last three amino acids (Δcys form) with Q79L mutation cloned in JB28 vector | Gift from Prof. Sean Munro (Addgene plasmid # 128900) |
| pcDNA3.3-Rab5A (S34N)-BirA*-HA-MAO | Rab5a lacking last three amino acids (Δcys form) with S34N mutation cloned in JB28 vector | This study |
| pEGFP-N1-SNX17 | Full-length SNX17 cloned in pEGFP-N1 vector | Gift from Prof. Lawrence Banks (ICGEB, Italy) |
| pmCherry-C1-VAPB | Full-length VAPB cloned in pmCherry-C1 vector | Gift from Prof. Gerald Rv Hammond (University of Pittsburgh School of Medicine, USA) |
| pHTN Halo Tag CMV-neo vector | Mammalian expression plasmid | Available in the lab |
| pHTN Halo Tag CMV-neo-SNX17 | Full-length SNX17 with N-terminal Halo tag cloned in pHTN Halo Tag CMV-neo vector | This study |
| pEGFP-C1-WASH | Full-length murine WASH with N-terminal GFP tag cloned in pEGFP-C1 vector | Gift from Prof. Robert Insall (University of Glasgow, UK) |
| pEGPN1-zebrafish *tecpr2*-201 | Zebrafish *tecpr2*-201 morpholino binding site cloned upstream of GFP tag in pEGFPN1 vector | This study |
| pEGPN1-zebrafish *tecpr2*-202 | Zebrafish *tecpr2*-202 morpholino binding site cloned upstream of GFP tag in pEGFPN1 vector | This study |
| ***Bacterial expression constructs:*** |  |  |
| pGEX-4T3 vector | Bacterial protein expression vector for making N-terminal GST proteins | Available in the lab |
| pGEX6P1-Rab5a (WT) | Full-length human Rab5a (WT) with N-terminal GST tag cloned in pGEX6P1 vector | Gift from Prof. Sunando Datta (IISER Bhopal, India) |
| pGEX6P1-Rab5a (Q79L) | Full-length human Rab5a with Q79L mutation and cloned in pGEX6P1 vector | This study |
| pGEX6P1-Rab5a (S34N) | Full-length human Rab5a with S34N mutation and cloned in pGEX6P1 vector | This study |
| pMAL-C2X vector | Bacterial protein expression vector for making N-terminal MBP-tagged proteins | Available in the lab |
| pMAL-C2X-TECPR2_TECPR (935-1411 aa) | Human TECPR2 from 935-1411 amino acid with N-terminal MBP tag cloned in pMAL-C2X vector | This study |
| pMAL-C2X-TECPR2_TECPR (935-1411 aa; R1336W) | Human TECPR2 from 935-1411 amino acid with R1336W mutation and cloned in pMAL-C2X vector | This study |
| pGEX6P1-GFP-Nanobody | Expresses GST-tagged anti-GFP Nanobody in *E. coli* cells | Gift from Prof. Kazuhisa Nakayama (Addgene plasmid # 61838) |
| Rab5a-Q79L-1-212-(C19S, C63S) | Bacterial protein expression vector for making N-terminal His- and SUMO-tagged Rab5a (1-212 aa) with Q79L, C19S, and C63S mutations | Gift from Prof. Roger L. Williams (MRC Laboratory of Molecular Biology, UK) |
| pGEX6P2-SNX17 | Full-length human SNX17 with N-terminal GST tag cloned in pGEX6P2 vector | This study |
| ***Yeast two-hybrid constructs:*** |  |  |
| pGBKT7 vector | GAL4-DNA binding domain yeast two-hybrid vector | Available in the lab |
| pGBKT7-Rab1a (Q70L) | Rab1a with Q70L mutation cloned in pGBKT7 vector | Gift from Prof. Marci Scidmore (Cornell University, USA) |
| pGBKT7-Rab2a (Q65L) | Rab2a with Q65L mutation cloned in pGBKT7 vector | Gift from Prof. Coenraad Kuijl (Amsterdam UMC, Amsterdam) |
| pGBKT7-Rab3b (Q81L) | Rab3b with Q81L mutation cloned in pGBKT7 vector | Gift from Prof. Coenraad Kuijl (Amsterdam UMC, Amsterdam) |
| pGBKT7-Rab4a (Q70L) | Rab4a with Q70L mutation cloned in pGBKT7 vector | Gift from Prof. Marci Scidmore (Cornell University, USA) |
| pGBDC1-Rab5a (WT) | Rab5a (WT) cloned in pGBDC1 vector | Gift from Prof. Tamotsu Yoshimori (Osaka University, Japan) |
| pGBDC1-Rab5a (Q79L) | Rab5 with Q79L mutation cloned in pGBDC1 vector | Gift from Prof. Tamotsu Yoshimori (Osaka University, Japan) |
| pGBDC1-Rab5a (S34N) | Rab5 with S34N mutation cloned in pGBDC1 vector | Gift from Prof. Tamotsu Yoshimori (Osaka University, Japan) |
| pGBKT7-Rab6a (Q72L) | Rab6a with Q72L mutation cloned in pGBKT7 vector | Gift from Prof. Marci Scidmore (Cornell University, USA) |
| pGBDC1-Rab7a (Q67L) | Rab7a with Q67L mutation cloned in pGBD-C1 vector | Gift from Prof. Tamotsu Yoshimori (Osaka University, Japan) |
| pGBKT7-Rab8a (Q67L) | Rab8a with Q67L mutation cloned in pGBKT7 vector | Gift from Prof. Coenraad Kuijl (Amsterdam UMC, Amsterdam) |
| pGBDC1-Rab9a (Q67L) | Rab9a with Q67L mutation cloned in pGBD-C1 vector | Gift from Prof. Tamotsu Yoshimori (Osaka University, Japan) |
| pGBKT7-Rab10 (Q48L) | Rab10 with Q48L mutation cloned in pGBKT7 vector | Gift from Prof. Marci Scidmore (Cornell University, USA) |
| pGBKT7-Rab11a (Q70L) | Rab11a with Q70L mutation cloned in pGBKT7 vector | Gift from Prof. Marci Scidmore (Cornell University, USA) |
| pGBKT7-Rab14 (Q70L) | Rab14 with Q70L mutation cloned in pGBKT7 vector | Gift from Prof. Coenraad Kuijl (Amsterdam UMC, Amsterdam) |
| pGBKT7-Rab18 (Q67L) | Rab18 with Q67L mutation cloned in pGBKT7 vector | Gift from Prof. Coenraad Kuijl (Amsterdam UMC, Amsterdam) |
| pGBKT7-Rab20 (Q67L) | Rab20 with Q67L mutant cloned in pGBKT7 vector | Gift from Prof. Coenraad Kuijl (Amsterdam UMC, Amsterdam) |
| pGBKT7-Rab23 (Q67L) | Rab23 with Q67L mutant cloned in pGBKT7 vector | Gift from Prof. Coenraad Kuijl (Amsterdam UMC, Amsterdam) |
| pGBKT7-Rab24 (S67L) | Rab24 with S67L mutation cloned in pGBKT7 vector | Gift from Prof. Coenraad Kuijl (Amsterdam UMC, Amsterdam) |
| pGBKT7-Rab29 (Q67L) | Rab29 with Q67L mutation cloned in pGBKT7 vector | This study |
| pGBKT7-Rab30 (Q68L) | Rab30 with Q68L mutation cloned in pGBKT7 vector | This study |
| pGBKT7-Rab33a (Q95L) | Rab33a with Q95L mutation cloned in pGBKT7 vector | This study |
| pGBKT7-Rab39 (Q72L) | Rab39 with Q72L mutation cloned in pGBKT7 vector | This study |
| pGBKT7-Vps11 | Vps11 cloned into pGBKT7 vector | Available in the lab |
| pGBKT7-Vps16 | Vps16 cloned into pGBKT7 vector | Available in the lab |
| pGBKT7-Vps33a | Vps33a cloned into pGBKT7 vector | Available in the lab |
| pGBKT7-Vps39 | Vps39 cloned into pGBKT7 vector | Available in the lab |
| pGBKT7-Vps41 | Vps41 cloned into pGBKT7 vector | Available in the lab |
| pGBKT7-LC3b | LC3b cloned into pGBKT7 vector | Available in the lab |
| pGBKT7-zRab5 | Zebrafish *Rab5* cloned into pGBKT7 vector | This study |
| pGADT7 vector | GAL4-activation domain yeast two-hybrid vector | Available in the lab |
| pGADT7-TECPR2 (WT) | Full-length human TECPR2 wild-type cloned into pGADT7 vector | This study |
| pGADT7-TECPR2 (1-944 aa) | Human TECPR2 from 1-944 amino acid cloned into pGADT7 vector | This study |
| pGADT7-TECPR2 (935-1411 aa) | Human TECPR2 from 935-1411 amino acid cloned into pGADT7 vector | This study |
| pGADT7-TECPR2 (I1256A) | Full-length human TECPR2 with I1256A mutation cloned into pGADT7 vector | This study |
| pGADT7-TECPR2 (E1299A) | Full-length human TECPR2 with E1299A mutation cloned into pGADT7 vector | This study |
| pGADT7-TECPR2 (T903M) | Full-length human TECPR2 with T903M mutation cloned into pGADT7 vector | This study |
| pGADT7-TECPR2 (D1000Y) | Full-length human TECPR2 with D1000Y mutation cloned into pGADT7 vector | This study |
| pGADT7-TECPR2 (W1140G) | Full-length human TECPR2 with W1140G mutation cloned into pGADT7 vector | This study |
| pGADT7-TECPR2 (R1336W) | Full-length human TECPR2 with R1336W mutation cloned into pGADT7 vector | This study |
| pGADT7-zTECPR2 | Zebrafish *TECPR2* (201 transcript; NCBI NP_001038644.1) cloned into pGADT7 vector | This study |

**Supplementary Table III:**
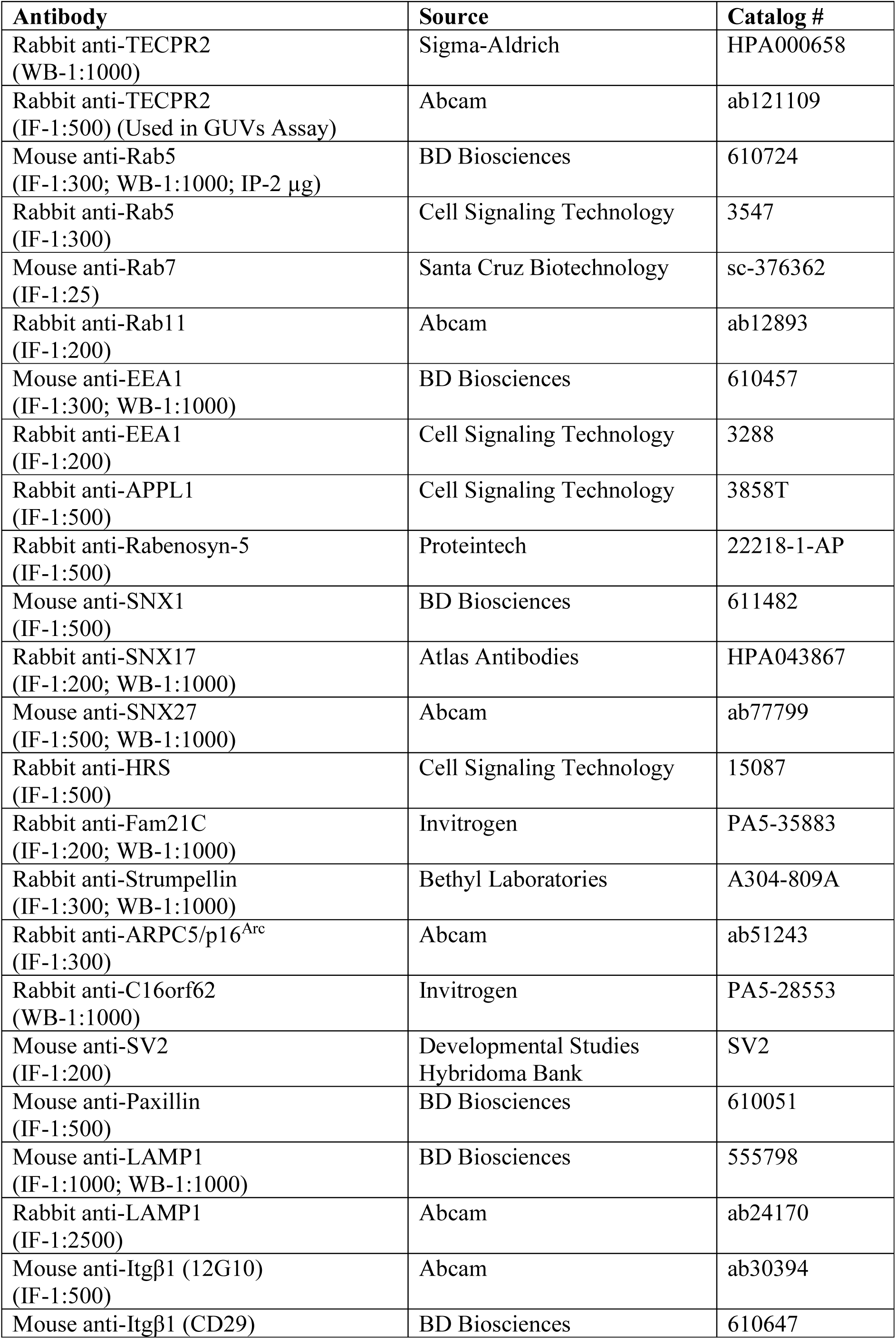
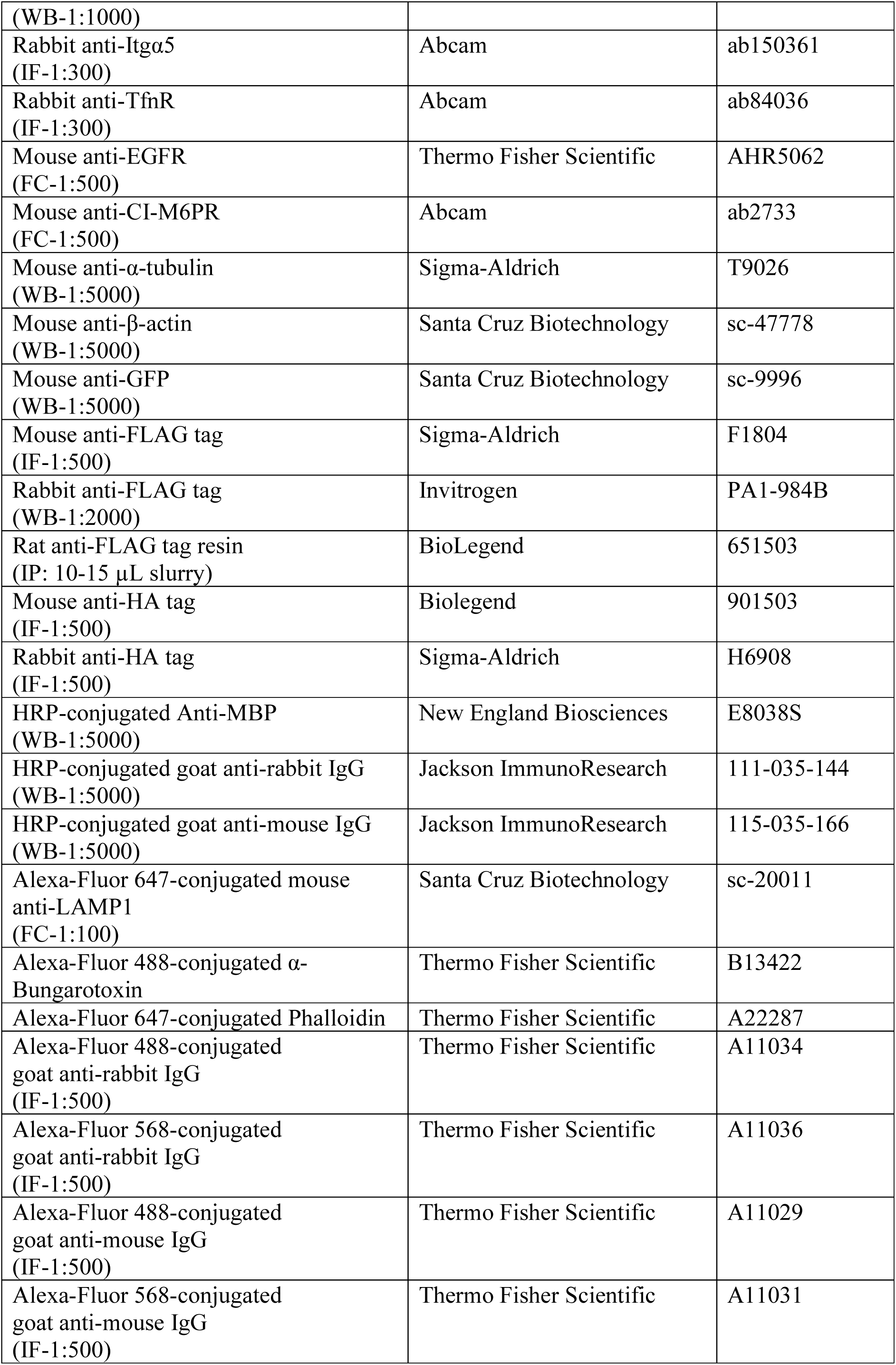
List of antibodies used in this study.

